# Split inteins for generating combinatorial non-ribosomal peptide libraries

**DOI:** 10.1101/2025.10.02.680031

**Authors:** Patrick Gonschorek, Christopher S. Wilson, Christian Schelhas, Kenan A. J. Bozhüyük, Peter Grün, Helge B. Bode

**Author notes:** These authors contributed equally to this work (Patrick Gonschorek, Christopher S. Wilson).

## Abstract

Engineering Non-Ribosomal Peptide Synthetases (NRPS) is a promising strategy for discovering new bioactive compounds, which can serve as valuable leads for drug development, such as new antibiotics. However, their engineering is hampered by the limited availability of molecular tools for the efficient heterologous expression of their large biosynthetic gene clusters. In fact, a single NRPS gene can already exceed the size limits of standard cloning vectors.

In this study, we establish split inteins as a novel tool for NRPS engineering to enable the expression of single, covalently linked NRPS proteins from multiple plasmids and to perform cloning-free module swapping. Using the xenotetrapeptide synthetase as model system, we show that an NRPS can be split into three parts and reconstituted via *trans*-splicing using two orthogonal inteins at four different engineering sites.

Based on this tripartite platform we build a library comprising 21 plasmids and generated 324 hybrid NRPS by combinatorial transformation. More than half were catalytically active, producing over 200 novel peptides. This intein-based technology provides a modular platform for generating natural product-like peptide libraries, expanding biocatalytically accessible chemical space.

## INTRODUCTION

Antibiotics such as penicillin, vancomycin, daptomycin, and colistin are natural products produced by complex biosynthetic gene culsters (BGC) involving non-ribosomal peptide synthetases (NRPS) as core enzymes. NRPS can incorporate structural features like cyclizations, acyl chains, D-amino acids or N-methylations, which confer favorable drug-like properties such as high affinity, target selectivity, proteolytic stability, and cell permeability^1,2^. Due to these diverse and biologically-validated scaffolds, natural products have historically been a rich source for drug discovery, accounting for around 30% of small-molecule drugs approved between 2000 and 2020^3-5^.

NRPS are large, multimodular enzymes that assemble peptide natural products through a stepwise condensation of amino acid building blocks. They have a repetitive modular structure, with a canonical module containing each of the three main domain-types: A condensation (C) domain, an adenylation (A) domain and a peptidyl carrier proteins (PCP), also referred to as a thiolation (T) domain. First, the A domains specifically select and activate amino acid substrates using ATP and covalently link them to the phosphopentatheine (Ppant) arm on the T domains. The C domains then catalyse the formation of the peptide bonds between two covalently bound amino acid substrates. NRPS assembly lines can also contain a variety of tailoring enzymes such as heterocylisation (Cy), epimerization (E), methyl transferases (MT), oxidation (Ox) and reduction (R) domains which can further modify and add to the structural diversity of the peptide. Finally, the peptide is released from the assembly line by a thioesterase (TE) domain which either hydrolyzes or cyclizes the peptide^6^.

Due to their modular structure, NRPS have long appealed as an opportunity to utilize synthetic biology’s toolkit to recombine domains to produce tailor-made peptides. Recent advances have brought this goal closer by adopting the strategy of splitting NRPS into exchangeable, tri-domain units either in front (XUT^I^) or within T domains (XUT^IV^)^7^, within condensation domains (XUC)^8^, or between C and A domains (XU)^9^. A systematic comparison of those four fusion sites has revealed that three sites - XUT^I^, XUT^IV^, and XU - consistently enable efficient generation of hybrid NRPS across a broad phylogenetic range, whereas the XUC site exhibits more limited compatibility^7^. Moreover, fusion sites located within linker regions, particularly XUT^I^ and XU, have been identified as optimal for maximizing production rates. Crucially, the XUT approaches have been successfully applied to the production of bioactive compounds, such as proteasome inhibitors^7^.

High throughput combinatorial shuffling of NRPS modules is a promising strategy to generate libraries of non-ribosomal peptides (NRP) for screening of novel bioactive compounds. While this has already been achieved at the DNA-level using Golden Gate cloning^10,11^, the size of the modules limits the number of modules that can be expressed from a single plasmid. Additionally, the large size of such plasmids makes them difficult to clone and transform. On average, each NRPS module is encoded by approximately 3 kilobases (kb) of DNA, contributing around 100 kDa to the overall molecular weight of the final enzyme. Low-copy expression plasmids, such as pACYC, can accommodate DNA inserts of up to 15 kb, which is sufficient for the insertion of up to five NRPS modules^12^. While some NRPS, such as the four modular xenotetrapeptide synthetase (XtpS, 12 kb, 458 kDa) fit on a single plasmid, the heterologous expression of larger known NRPS proteins like the kolossin syntetase (KolS, 49 kbp, 1.8 MDa, would require even three plasmids or other vectors like bacterial artificial chromosomes (BAC)^13,14^.

In contrast to KolS, some NRPS assembly lines naturally consist of multiple proteins that interact through native docking domains^15^. In these cases, the BGC can be readily divided across multiple plasmids for heterologous expression. However, as the architecture of each BGC is unique and cluster-specific, docking domains do not consistently appear at the same positions, preventing this approach from being applied as a universal strategy to split clusters onto separate plasmids. Moreover, natural docking domains are frequently located downstream of T domains, a position that is unfavorable for the construction of artificial NRPS hybrids.

One strategy to overcome these limitations has been the use of SYNZIPs (SZ), which enable recombination of NRPS at the protein level^16^. SZ are synthetic, coiled-coil peptides that are designed to specifically interact with high affinity^17^. Even though their affinity lies in the nanomolar range, their interaction is non-covalent and the 60 Å-long zipper pair remains part of the protein complex, possibly sterically impairing the NRPS assembly line and reducing production rates. Furthermore, to achieve the closest possible proximity between the C- and N-termini of a split protein, anti-parallel SZ are required. However, only one such pair (SZ17 and SZ18) has been described to date, meaning there is a lack of orthogonal, anti-parallel SZ pairs for efficiently constructing multi-partite systems^18^.

A well-established protein engineering tool that could address all of the aforementioned NRPS engineering challenges are split inteins^19^. Inteins (internal proteins) are small, self-excising protein segments that are found within the genomes of diverse organisms, being most abundant in prokaryotes, particularly in archaea. They possess the remarkable ability to catalyze their own removal from a precursor protein, fusing the remaining external protein fragments (exteins) covalently together by forming a new amide bond, a process named protein splicing. Inteins often occur in essential proteins and restore the protein’s activity after excision^20^. They exist either within a single continuous protein or are naturally split into two halves attached to two independently expressed proteins. Split inteins have two exteins attached to each intein half and can self-associate driven by electrostatic interactions, catalysing *trans* protein splicing^21^. The catalytic activity of the intein is affected by the adjacent residues in the extein sequences, particularly requiring a nucleophilic side chain (serine, threonine or cysteine) in the first position of the C-terminal extein for catalysis. For engineering purposes, typically three amino acids from the native extein sequence are incorporated into the target protein, along with the intein on both sides^22^. Inteins have already successfully been used for various biotechnological applications, such as the generation of more proteolytically- and thermostable proteins^23^, the generations of cylic peptide libraries^24^, the incorporation of noncanonical amino acids into protein in living cells^25^, or the delivery of large protein like dystrophin via AAV gene therapy^26^.

In this work, we have established split inteins for the expression of a single NRPS protein from multiple plasmids. We started by testing five orthogonal inteins to resconstitute a model NRPS system, split at one of four established NRPS engineering sites (XUT^I^, XUT^II^, XUC, XU). For each site we then selected the two best performing inteins and used them to reconstitute an NRPS assembly line encoded on three separate plasmids. Finally, using the tripartite system split at the XUT^I^ site, we built a NRPS library, that through simple combinatorial plasmid transformation without additional cloning, yields 324 possible hybrid NRPS combinations. Analyzing the production of all combinations, we gained novel insights into NRPS module compatibility. Furthermore, the library produces a structurally diverse set of over 200 NRP, highlighting the robustness of the intein-based platform for generating peptide libraries.

## RESULTS

### NRPS reconstitution at multiple fusion sites

To evaluate the potential of inteins as tools for NRPS engineering we chose the xenotetrapeptide (XTP) producing NRPS called xenotetrapeptide synthetase (XtpS) from the Gram-negative, entomopathogenic bacterium *Xenorhabdus nematophila* as model system^27^.

The NRPS was heterologusly expressed in *E. coli* DH10B::*mtaA* strain, which constitutively expresses the broad-spectrum Ppant transferase MtaA, for posttranslational activation of the T domain^28^.

First, we intended to split the NRPS into two parts and restore its activity by *trans*-splicing using a single split intein, while comparing various fusion sites and intein variants. Four sites within the NRPS were chosen for splitting and intein insertion based on their demonstrated suitability for NRPS engineering, ensuring their applicability for subsequent NRPS library generation^7-9^.

Using either the XUT^I^, XUT^II^, XUC or XU fusion site, all located between the second and third A domain (**Fig. 1a**), both halves were cloned on separate expression vectors. To reconstitute the NRPS after expression via protein-splicing, we selected the five inteins gp41-1, gp41-8, NrdJ-1, IMPDH-1 and SspGyrB based on splicing efficiency and orthogonality as characterized previously^29^. The N-terminal fragment of the inteins (intein_N_) was fused to C-terminus of the first half of the NRPS (extein_N_) and the C-terminal fragments of the inteins (intein_C_) N-terminally to the second half of the NRPS (extein_C_). To visualize the splicing complex at the XUT^I^ position we generated a structural model using AlphaFold3 (**Fig. 1b**).

**Figure 1.**
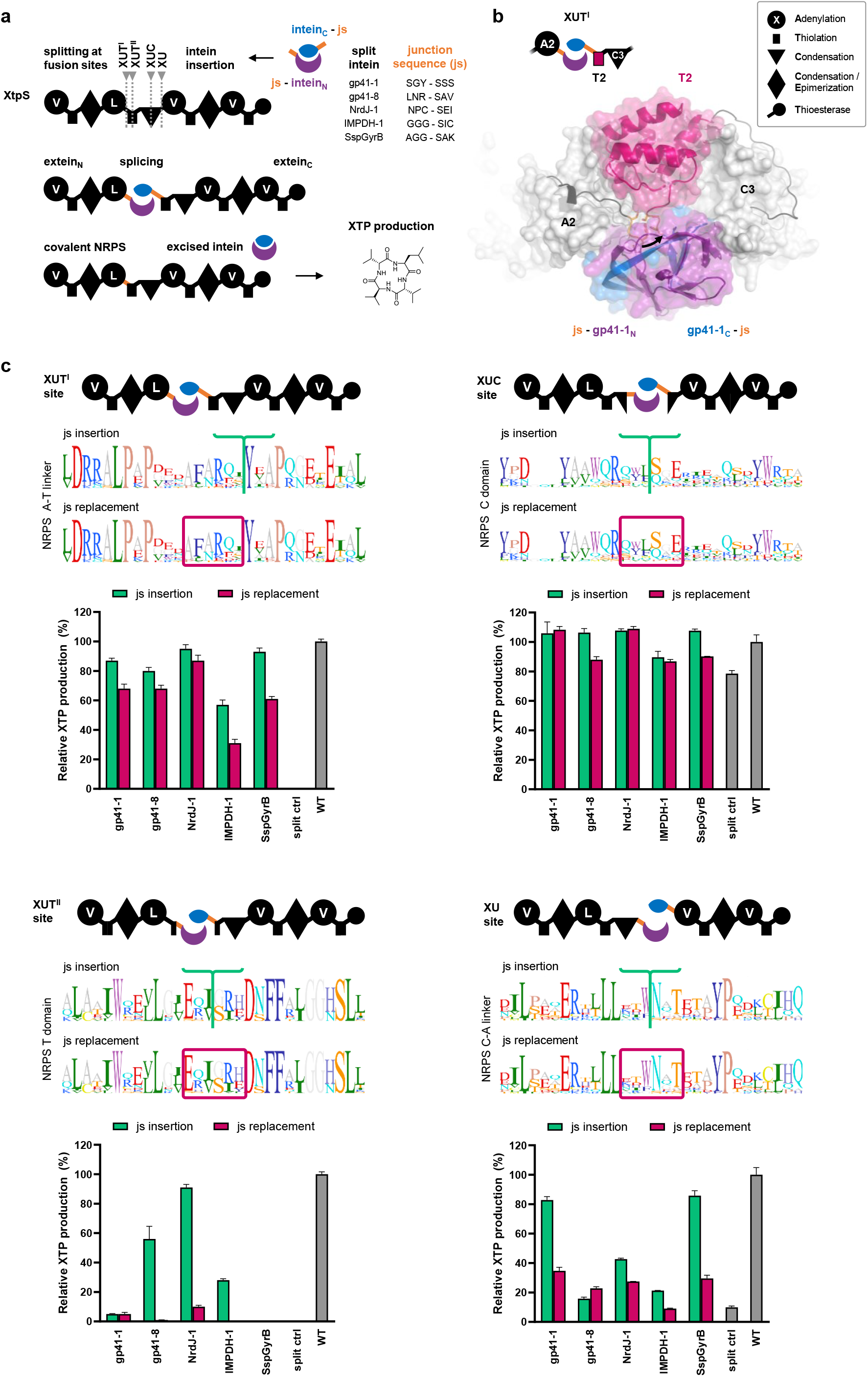
Establishing NRPS splicing at four different engineering sites. (**a**) Schematic representation of the Xenotetrapeptide synthetase (XtpS) NRPS (shown in black), used as a model system, with the four sites used for intein insertion indicated by gray arrows. The NRPS gene was split at one of the four fusion sites, and each half was linked to the N-terminal (purple) or C-terminal half (blue) one of five different split inteins. To ensure efficient splicing, three residues from the native junction sequences of the intein (js, orange) were used as linkers. These js residues remain in the spliced NRPS, resulting in a six-amino acid scar, with three residues coming from each side (js residues shown for each intein). The two NRPS-intein fusion proteins were expressed from separate plasmids. Successful *trans*-splicing should result in Xenotetrapeptide (XTP) production. (**b**) Structure model of the splicing complex at the XUT^I^ site between the A2 and T2 domains of XtpS generated with AlphaFold 3. The new peptide bond formed through splicing of the gp41-1 intein is shown as black arrow. (**c**) Sequence logos of the four NRPS regions locating the tested fusion sites (XUT^I^, XUT^II^, XUC and XU) are shown. For each site, two intein-insertion strategies were compared: Insertion without amino acid deletion, which preserves all native NRPS residues but results in a six-residue longer NRPS after splicing (insertion sites indicated with green brackets, green bars); versus replacement of six NRPS residues with the intein scar while maintaining the original protein length (relaced residues shown in red box, red bars). Bars represent the relative XTP production of intein-spliced XtpS compared with native XtpS (“WT”) as determined by LC-MS analysis. Identically split NRPS constructs but without attached inteins were used as negative controls (“split ctrl”, gray bars). Data is presented as mean ± SD (n = 3, biological replicates). NRPS domain explanations are shown in the box.

Previous studies have shown that incorporating the three extein residues directly adjacent to the splicing site as a linker to the intein in a non-native protein is crucial for maintaining optimal splicing efficiency^22^. These residues are commonly referred to as junction sequence (js), are specific to each intein and, unlike the intein itself, remain in the spliced protein, resulting in a six-residue ‘scar’. We decided to adopt this strategy and use three-residues native js for each intein as linker to the NRPS. To accommodate the resulting scar sequences in the spliced NRPS, we compared two intein-insertion approaches: (i) ‘js insertion’, involved inserting the inteins without deleting any native NRPS residues. While this approach preserves all NRPS residues, it results in an NRPS protein that is six residues longer after splicing. (ii) ‘js replacement’ involved deleting six NRPS residues at the splicing site to make room for the intein scar, thereby maintaining the original protein length and domain spacing.

Covalent protein formation was confirmed by Western blot, which showed the expected 458 kDa band corresponding to full-length XtpS at all four fusion sites and only weak or no bands for the individual non-spliced proteins (**Fig. S5**).

In order to evaluate the efficiency of different inteins at various fusion sites, XTP production by the intein-spliced NRPS was compared to that of the full-length NRPS directly expressed as single full-length protein (**Fig. 1c**). For all fusion sites, we observed that at least one intein was able to almost fully restore catalytic activity. However, the relative production varied considerably depending on the fusion site and the intein used. At the XUT^I^ and XUC sites, most inteins were able to restore a high level of production, with little variation between the specific intein used. In contrast, at the XUT^II^ and XU sites, intein performance varied, indicating that the observed differences are likely due to the sensitivity of the NRPS protein to the insertion or replacement of js residues, rather than differences in the splicing efficiency of the inteins themselves. In particular the XUT^II^ site, which is located within the T domain, seems to tolerate the gp41-8 and NrdJ-1 js residues better than those of the other three tested inteins. Furthermore, for all four fusion sites and nearly all inteins, the ‘js insertion’ strategy led to better productivity than the ‘js replacement’ approach, showing that preserving all NRPS residues is more critical for catalytic activity than preserving domain distances. This important insight guided the decision to exclusively use the ‘js insertion’ strategy in all subsequent constructs.

A rather surprising finding emerged from analyzing the production of the split NRPS constructs lacking attached inteins, which were generated as negative controls: While the NRPS split at the XUT^I^ and XUT^II^ did not produce any XTP and only 10 ± 1 % split at the XU site, the protein split at the XUC site still showed 79 ± 2 % productivity. This reveals that the N- and C-terminal lobes of C domains interact strongly enough to maintain all catalytically relevant conformations without the need for a covalent linkage between the lobes.

### Combining orthogonal inteins to build tripartite NRPS systems

We next sought to go one step further by splitting the NRPS into three parts and reconstituting it using two orthogonal split inteins. Such a system would be a valuable tool for NRPS engineering, as it would allow the simultaneous recombination of initiation, elongation, and termination units within NRPS assembly lines, without the need for cloning. We chose the intein pair gp41-8 and NrdJ-1 for the XUT^I^ and XUT^II^ sites, and used gp41-1 together with SspGyrB at the XUC and XU sites. Our model NRPS was expressed from three orthogonal plasmids, with splits introduced between the 2^nd^ and 3^rd^, and the 3^rd^ and 4^th^ A domains.

When testing the tripartite systems, we observed XTP production at all four fusion sites, with 59 ± 5% at XUT^I^, 25 ± 1% at XUT^II^, 98 ± 7% at XUC, and 65 ± 7% at XU (**Figure 2**). This demonstrates that NRPS proteins can be successfully reconstituted through trans-splicing from three parts at all established NRPS engineering sites, including the XUT^II^ site within the T domain, where the use of previously published SYNZIPs would be difficult.

**Figure 2.**
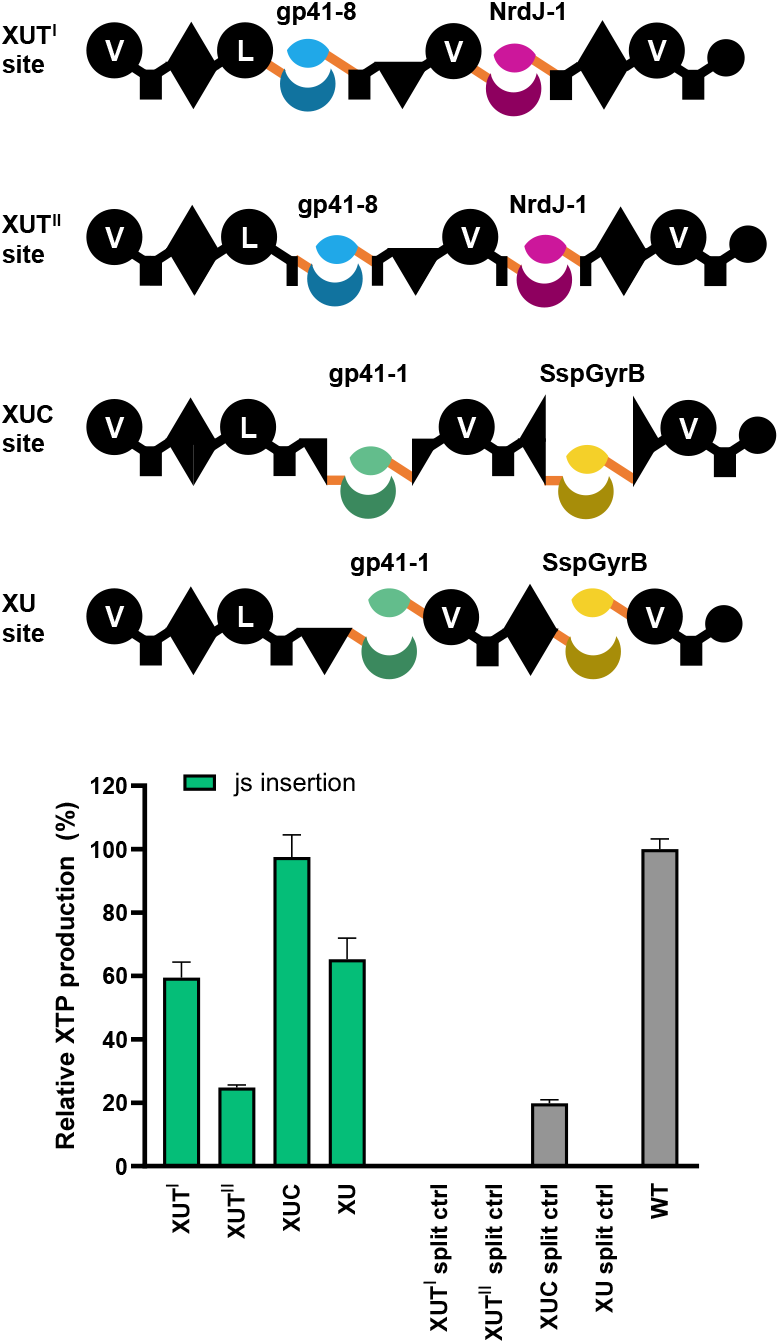
NRPS splicing from three parts by combining two orthogonal split inteins. Two orthogonal split inteins were used to reconstitute XtpS split into three parts at four different fusion sites. For the XUT sites, gp41-8 was combined with NrdJ-1. For the XUC and XU sites, gp41-1 was combined with SspGyrB. Bars represent the relative XTP production of intein-spliced XtpS compared with native XtpS (“WT”) as determined by LC-MS analysis. For the XU and XUC sites, identically split NRPS constructs but without attached inteins were used as negative controls (“split ctrl”, gray bars). Data is presented as mean ± SD (n = 3, biological replicates). For NRPS domain explanations see Fig. 1.

Among the four sites, the XUC site showed the highest production rates at 98% ± 7%. However, 20 ± 1% of the activity was also observed in the XUC negative control, which suggests a contribution from the interaction of the C domain lobes to the production rate as previously observed in the single split system.

### Generating a proof-of-concept NRPS library

After successfully establishing the split intein technology at various fusion sites, we wanted to demonstrate its applicability for the cloning-free generation of NRPS libraries. In order to design a proof-of-concept library, we chose the tripartite system split at the XUT^I^ site reconstituted with the two inteins gp41-8 and NrdJ-1 for all constructs. This fusion site had worked very effectively for restoring XTP production at 59 ± 5% and additionally had been shown in previous studies to have greater success at shuffling NRPS units from a broad taxonomic range^7^.

NRPS modules were derived from a variety of different BGCs originating from 12 different strains, mostly *Xenorhabdus*, one *Photohabdus* species of bacteria, as well as one module from the fungus *Mortierella alpine* (**Tab. S2, Fig. S3, Fig. S4**). Modules were selected with compatibility considerations in mind, focusing on modules that had previously demonstrated functionality when engineered in the context of XtpS or related NRPS systems^7-9,16,30^.

We designed, six plasmids encoding initiation modules, nine encoding elongation modules, and six termination modules, resulting in a library of 21 plasmids. Simple combinatorial transformation of sets of three plasmids allows the expression of 324 recombinant NRPS, that were anticipated to produce a structurally diverse range of NRP.

The samples were measured by high-resolution liquid chromatography–tandem mass spectrometry (HR-LC-MS/MS) searching for the expected peptide masses. A NRPS combination was considered catalytically active, if at least one of the expected linear or cyclic NRP could be found. Peptides that did not match expected products, like truncated peptides, were not considered in this analysis. Using this analysis strategy we found that out of the 324 expressed NRPS, 173 (53%) were catalytically active, among which 46 produced cyclic, and 148 produced linear peptides. Some modules can incorporate multiple building blocks, for example, module E can produce peptides with either a pentanoyl or hexanoyl acyl chain, and modules M, N, and U can each incorporate two different amino acids. Therefore, multiple peptide derivatives can be found in some NRPS combinations. Considering also these derivatives, a total of 258 NRP were identified - 201 linear and 57 cyclic (**Tab. S7, Fig. S2**) - without searching for all possible derivative combinations.

As the detectability of an NRP can vary depending on the particular LC-MS system used, we analyzed all 324 combinations using two different LC-MS systems to validate the results. While the data shown in **Figure 3** were obtained on a HR-LC-MS/MS system we also measure all samples on low-resolution LC-MS/MS ion-trap system and results are shown in **Fig. S6**. For 272 (84%) out of the 324 of the NRPS combinations, identical peptides as identified with the HR system were detected.

**Figure 3.**
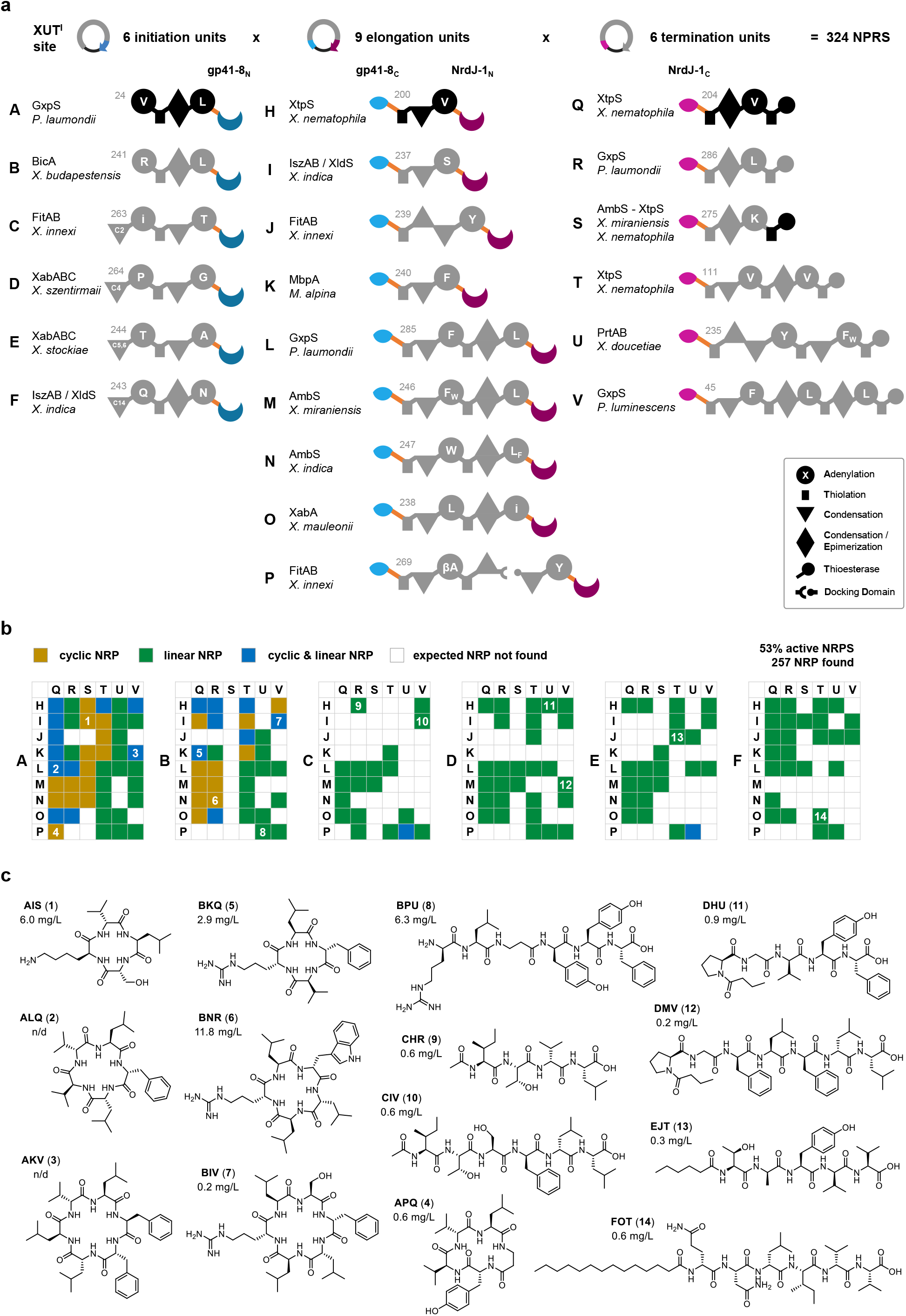
Combinatorial NRPS Library with 324 variants generated via split-intein technology. (**a**) NPRS library was generated based on the tripartite system split at the XUT^I^ site and reconstituted by splicing using the orthogonal split inteins gp41-8 and NrdJ-1. 21 NRPS exchange units were derived from biosynthetic gene clusters of twelve *Photorhabdus* and *Xenorhabdus* strains, as well as the fungal species *Mortierella alpina*. The library comprised six initiation units (A–F), nine extension units (H–P), and six termination units (labeled Q–V), each cloned into one of three compatible plasmid vectors (plasmid numbers in grey). Co-expression of all possible triplet combinations yielded 324 distinct *E. coli* strains expressing a single NRPS assembled by *trans*-splicing. (**b**) Each of the 324 strains was individually analyzed for production of the expected linear or cyclic NRP using HR-LC-MS. For each initiation unit (A-F), a matrix visualizes whether any of the expected cyclic NRP (yellow), linear NRP (green), or both (blue) were found. (**c**) Chemical structures of fourteen peptides (compounds 1-14) that were selected for chemical synthesis for to confirm their structures and quantify production titers. The selected peptides are indicated by numbers in the matrices in panel (b) and the three-letter codes refers to the respective NRPS constructs. Peptides were selected at random from among the active NRPS variants, ensuring representation of at least one product from each of the 21 modules used. Production titers could be determined for twelve peptides. For NRPS domain explanations see box.

To quantify production titers and to confirm predicted NRP structures, fourteen peptides (1-14, three-letter code refers to the respective NRPS module combination) were selected for chemical synthesis. The peptides were chosen from among the functional NRPS variants, ensuring that at least one product from each of the 21 modules used is represented. Production titers could be determined for twelve peptides (**1** and **4-14**), while titers for peptide **2** (ALQ) and **3** (AKV) could not be reliably determined due to poor solubility. Production titers varied between 0.2 and 12 mg/L, corresponding to total yields ranging from 1 to 47 µg from the 4 mL production cultures. Chemical structures were confirmed by HR-LC-MS/MS and comparison of retention times with synthetic standards. For all fourteen peptides (**1-14**), retention times, HR-MS masses and fragmentations did match the synthetic standards (**Fig. S1**). For peptides **9** (CHR) and **10** (CIV), two peaks with identical masses but slightly different retention times were observed. For both, the earlier-eluting peak matched the synthetic standard, suggesting that the second peak corresponds to a stereoisomer arising from a partially deficient epimerization activity of one of the dual C/E domains in modules R and V. In summary, all fourteen chemical structures were confirmed, demonstrating that NRP structures in hybrid NRPS libraries can be reliably predicted.

### Accessing natural product-like chemical space

To assess the potential of such a NRP library for bioactive compound discovery, we analyzed the chemical space covered by this library. While originally Lipinski’s rule of five (Ro5) defined a chemical space for molecules with favorable drug-likeness in particular in terms of oral bioavailability, natural product cyclic peptides often are found in the beyond Ro5 (bRo5) chemical space^31^. We assumed that NRP libraries explore the chemical space surrounding the NP used for their design, and that they may also occupy the bRo5 chemical space and thus might provide favorable pharmacokinetic properties by design. To confirm this hypothesis we generated a chemical space map to visualize the relationships between cyclic, linear NRP, and a reference set of 2’584 FDA-approved drugs^32^ by creating Morgan fingerprints for each molecule and applying Uniform Manifold Approximation and Projection (UMAP) for dimensionality reduction (**Figure 4**). This map revealed that linear NRP did cluster according to the starter unit used. Each of the six clusters corresponding to a specific starter module was found in a distinct region of the map rather at the edge of the space defined by approved drugs, but still overlapping with some. Interestingly, all cyclic NRP of the library were found in another distinct region of the chemical space, which corresponds to the bRo5 space, were natural product drugs such as capreomycin, romidepsin, colistin and pasireotide were located. While some of the NPs used for library generation (fitayylide^7^, protegomycin^33^, ambactin^34^) were found in this space as well, all novel peptides are distributed in the bRo5 space, sampling this region way beyond their parent NPs, highlighting that such cyclic NRP libraries can be used for accessing bRo5 chemical space. Additionally, from the split intein library 11 new cyclic tetra-peptides were produced, which corresponds to ∼17% of the known natural cyclic tetra-peptides described previously^35^.

**Figure 4.**
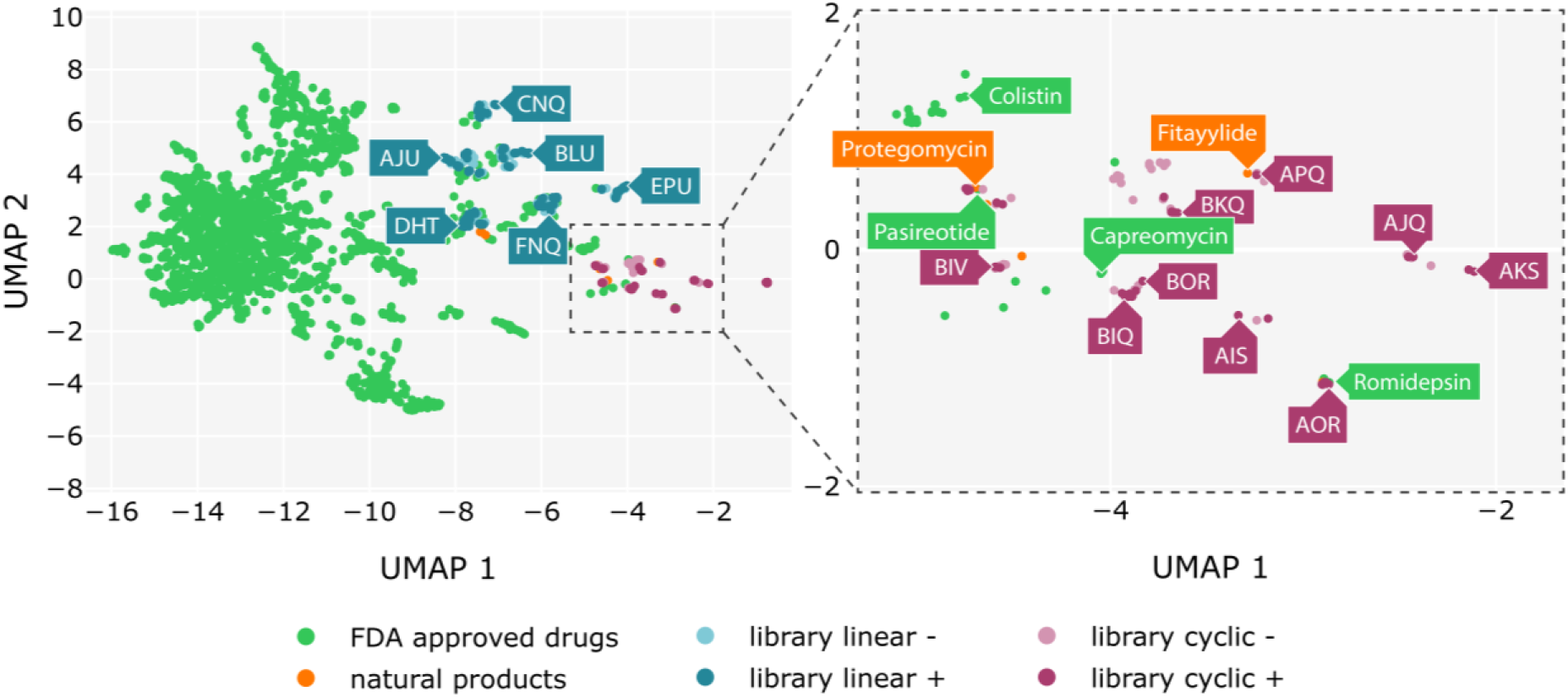
Chemical space accessible with the generated NRP library. Visualization of the chemical space occupied by all NRPs produced from the NRPS library (Figure 3) relative to a reference set of 2,584 FDA-approved drugs. Chemical space was mapped by generating Morgan fingerprints for each molecule and applying Uniform Manifold Approximation and Projection (UMAP) for dimensionality reduction. FDA-approved drugs (green dots), natural products from native BGCs used for library generation (orange), linear NRP are depicted in blue (detected peptides in dark blue, predicted but not detected in light blue), and cyclic NRPs are shown in red (dark red for detected and light red for predicted but not detected). A zoomed-in view highlights the region commonly referred to as beyond rule of five (bRo5) chemical space.

## DISCUSSION

Engineering NRPS is a compelling strategy for expanding the repertoire of accessible natural products, yet widespread application has been limited by technical challenges in cloning and expressing the large BGCs encoding those enzymes. Our study demonstrates for the first time that split inteins can be systematically applied as tools for NRPS engineering, offering a modular, robust, and cloning-free approach to assemble and diversify those large enzyme machineries once the different NRPS fragments are available on plasmids.

We have shown that split inteins can efficiently reconstitute catalytically active NRPS enzymes from split gene constructs expressed from up to three separate plasmids. Our systematic analysis of four established NRPS engineering fusion sites, combined with five orthogonal split inteins, revealed a functional pair of orthogonal intein for each fusion site. This is a notable advancement over previous protein engineering approaches, such as SYNZIP-mediated non-covalent assembly, which is limited by both the number of available orthogonal pairs and potential steric hindrance of the persistent SYNZIPs. By contrast, intein-mediated trans-splicing results in native-like, covalently linked enzymes, minimizing interference with catalytic function. Our comparison of two strategies for handling the six residue junction sequences revealed that preserving all residues of the NRPS is more important for catalytic activity than retaining precise domain spacings, which is a general principle that can guide future NRPS engineering efforts. Unexpectedly, we observed significant residual activity for C-domain–split constructs without inteins, revealing strong natural docking between C-domain lobes, which will be analyzed in more detail in the future. While not the focus of this work, this phenomenon raises structural questions about mechanism of C domains and how their lobes interact during catalysis.

The intein-XUC system allows splitting NRPS without losses in production titers, exploiting the intrinsic C-lobe interaction to promote splicing. This makes this system good choice for heterologous expression and other applications where only the native NRPS should be restored. The XUT^I^ system showed to be ideal for combinatorial NRPS library generation. As the XU system allows generation of hybrid NRPS with similar success rates, this system is likely equally suited for the generation of NRPS libraries and would need to be systematically compared again to the XUT^I^ site with a bigger data set.

We have shown the application of split inteins for generating NRPS libraries through combinatorial plasmid transformation, bypassing cloning challenges and size limits of classical DNA-level shuffling. Using only 21 plasmids, we biocatalytically produced 258 NRP in *E. coli*, which is to our knowledge the largest NRP library reported by now.

Notably, the peptides generated here effectively explore regions in the bRo5 chemical space, where many successful natural-product-derived drugs reside^14^. This work demonstrates that genetically encoded NPR libraries allow for the fast, cost-effective and sustainable generation of peptide libraries, which can be used for hit-identification within a unique, natural product-like chemical space. In contrast to synthetic macrocycle libraries, such genetically encoded compound libraries make the production and screening of natural product-like macrocycles easier and more broadly accessible to the research community and allow upscaling simply by using larger cultivation vessels.

Despite the high overall success rate, approximately half of the recombined NRPS hybrids were inactive, suggesting context-dependence and module compatibility issues in hybrid NRPS. While we relied on pre-characterized modules in this work, this technology also allows fast identification of compatible modules for future libraries. Furthermore, it facilitates the generation of large datasets needed to tackle the longstanding module-compatibility problem using deep learning approaches^36^. Finally, translation to other systems, such as polyketide synthases, hybrid PKS-NRPS, or even other modular enzymes, appears promising.

With a set of fully compatible exchange units, such libraries have the potential to unlock the vast chemical space of NRP for bioactive compound discovery. The benefit of the split intein library approach is that the library can easily be expanded by adding additional plasmids to generate greater chemical diversity in the future. The efficiency of this approach enables the exploration and expansion of chemical space beyond what is currently represented in existing datasets, as shown for cyclic tetrapeptides, increasing the likelihood of discovering compounds with new and diverse biological activities.

## ACKNOWLEDGEMENTS

This work was supported by the Max-Planck Society and an ERC Advanced Grant (835108) to HBB. We thank Tania Köbel from the MaxGENESYS Biofoundry for providing the initial high-throughput protocols. Tania Köbel, Timon A. Lindeboom and René Inckemann for supporting the operation of the equipment. Trinetri Goel and Chloé Puteaux for providing protocols and support for SPPS and purification.

## AUTHOR CONTRIBUTIONS

PGo and CSW designed and conducted most experiments and analyzed the data, PGr assisted with measuring and analyzing LC-MS data, PGo, CS and PGr wrote scripts for automated data analysis, CS, CSW and PGo conducted SPPS. KAJB was involved in early conceptual discussions and PGo created all figures and wrote the manuscript with input from all authors. PGo and HBB planned and supervised the project.

## Notes

### Competing Interest Statement

The authors have declared no competing interest.

### Summary of Updates

Figure 1 was not shown completely so we uploaded a new version of the manuscript.

