## Supplementary information_Tables_Figures for "Split inteins for generating combinatorial non-ribosomal peptide libraries"

#### MATERIALS AND METHODS

##### Cultivation of strains

*E. coli DH10B::mtaA* were cultivated in LB medium (10 g/L tryptone, 5 g/L yeast extract, 5 g/L NaCl and pH 7.5). Medium for overnight cultures was supplemented with kanamycin (50 µg/mL or 25 µg/mL, sterile in $ddH_2O$) chloramphenicol (34 µg/mL in ethanol) and spectinomycin (100 µg/mL, sterile in $ddH_2O$). For cryopreservation, overnight liquid cultures were mixed 1:1 (v/v) with 50% glycerol (v/v) and stored at -80°C.

##### Genomic DNA (gDNA) preparation

To obtain genomic DNA of various *Photorhabdus* and *Xenorhabdus* strains a single colony was inoculated into a 10 mL LB medium and incubated at 28°C for 24 h while shaking at 200 rpm. To extract the gDNA a Monarch Genomic DNA Purification Kit (NEB) kit was used and followed according the manufacturer's instructions.

##### Polymerase chain reaction (PCR)

PCR amplification of DNA fragments and vector backbones was performed with Q5 High-Fidelity DNA Polymerase (NEB) using primers with overhangs and the following reaction mix: 10 µL Q5 Reaction Buffer, 1 µL 10 mM dNTPs, 1 µL forward primer 10 µM, 1 µL reverse primer 10 µM, 1 µL (~ 20 ng/µL) template DNA, 0.5 µL Q5 High-Fidelity DNA Polymerase, 3 µL DMSO, 32.5 µL nuclease-free water. If plasmids were used as templates, template was digested using 1 µL DpnI and incubation 37°C for 20 min. PCR products were validated and isolated by 1% agarose gel electrophoresis in TAE buffer stained with MIDORI Green Advance (0.2 µL/100 mL, NIPPON Genetics) and extracted using the Monarch DNA Gel Extraction Kit (NEB).

##### Cloning of biosynthetic gene clusters

pACYC, pCOLA and pCDF with arabinose inducbe pBAD promoters were used as backbones for generating intein vectors. Intein gene fragments were ordered from Twist bioscience. XtpS was cloned onto a pCOLA based plasmid and used as a template for amplifying modules of XtpS. DNA sequences of gp41-1, gp41-8, NrdJ-1, IMPDH-1 and SspGyrB from Pinto *et al.* were ordered as synthetic DNA fragments from Twist[1]. All constructs were constructed with Gibson assembly using DNA fragments with 25-30 bp overlapping homologous arms. In the assembly reaction using 20 fmol backbone fragment and 4 molar equivalents of each insert fragment were used. Fragments were assembled with NEBuilder HiFi DNA Assembly Master Mix incubating at 50 °C for 1 h. Then 1 µL reaction mix was transformed into chemically competent E. coli DH10B cells. Plasmids were confirmed by overnight sequencing (Microsynth) using Sanger sequencing according to the companies protocol. Correct plasmids were isolated the next day from overnight cultures using the Monarch plasmid miniprep kit (New England Biolabs) using the manufacturers protocol.

#### Preparation of chemically competent cells

*E. coli DH10B::mtaA* colonies were inoculated into 5 mL CVM media (20 g/L tryptone, 5 g/L yeast extract, 0.7 g/L KCl, 4.9 g/L $MgSO_4 \cdot 7H_2O$) and incubated at 37°C overnight. Overnight culture was diluted 1:100 and incubated at 37°C till an $OD_{600}$ of 0.5. Culture was placed on ice and centrifuged at 4°C at 2500 rpm for 10 min. Supernatant was discarded and pellet was re-suspended in buffer 1 (15% glycerol (v/v), 2.94 g/L potassium acetate, 1.47 g/L $CaCl_2 \cdot 2H_2O$, 12.1 g/L RbCl, 7.2 g/L $MnCl_2 \cdot H_2O$, pH = 5.9). Cells were kept on ice for 90 min then spun at 2500 rpm at 4°C for 10 minutes. The supernatant was discarded and the pellet was resuspended in buffer 2 (15% glycerol (v/v), 0.42 g/L MOPS, 0.24 g/L RbCl, 2.21 g/L $CaCl_2 \cdot 2H_2O$, pH = 7.0) and aliquots were prepared for storage in −80 °C freezer.

#### Plasmid transformation

For transformation cells were thawed on ice and ~100 ng of plasmid DNA was added to the cells. Cells were incubated on ice for 20 minutes, subjected to a heat shock at 42 °C for 1 minute, placed on ice for an additional 3 minutes, and subsequently regenerated in SOC medium (10 g/L tryptone, 2.5 g/L yeast extract, 0.25 g/L NaCl, 0.09 g/L KCl, 1.0 g/L $MgCl_2 \cdot 6H_2O$, 1.23 g/L $MgSO_4 \cdot 7H_2O$, 0.9 g/L glucose) for 1 h at 37°C and then plated onto LB agar with respective antibiotics.

#### NRPS expression and NRP extraction

100 µl of overnight culture was inoculated into 10 mL XPP3 medium (10 g/L glycerol, 20 mL/L M9 salt A, 20 mL/L M9 salt B, 20 g/L Proteose Peptone No. 3; M9 salt A: 350 g/L $K_2HPO_4$, 100 g/L $KH_2PO_4$; M9 salt B: 29.4 g/L sodium citrate, 50 g/L $(NH_4)_2SO_4$, 5 g/L $MgSO_4$) complemented with 2 mL/L vitamins (10 mg/L folic acid, 6 mg/L biotin, 200 mg/L p-aminobenzoic acid, 1 g/L thiamine-HCl, 1.2 g/L pantothenic acid, 2.3 g/L nicotinic acid, 12 g/L pyridoxine-HCl, 20 mg/L vitamin B12, 100 mg/L riboflavin) and 1 mL/L trace elements (40 mg/L $ZnCl_2$, 200 mg/L $FeCl_3 \cdot 6H_2O$, 10 mg/L $CuCl_2 \cdot 2H_2O$, 10 mg/L $MnCl_2 \cdot 4H_2O$, 10 mg/L

$Na_2B_4O_7 \cdot 10H_2O$, 10 mg/L $(NH_4)_6Mo_7O_{24} \cdot 4H_2O$) per liter medium. Antibiotics were used at ⅓ of the previously used concentrations (11 µg/ml chloramphenicol, 17 µg/ml kanamycin, spectinomycin 33 µg/m). Arabinose was added to a final concentration of 0.2% (v/v) and 300 µL Amberlite XAD16N beads (20-60 mesh, Sigma Aldrich) were added per 10 mL culture.

Cultures were incubated at 25°C for 48 h shaken at 200 rpm. For extraction, the medium was decanted and the XAD beads were incubated with 10 mL of a 1:1 (v/v) mixture of methanol and acetonitrile, and shaken for 30 minutes at 200 rpm. 500 µL aliquot of the extract were centrifuged at 13000 rpm for 20 minutes and 100 µL of this was assayed on the HPLC-MS.

#### LC-MS/MS analysis

HPLC-MS was performed using an Agilent UPLC 1290 Infinity II system with a ACQUITY Premier BEH C18 Column (186009453, Waters) coupled to an AmaZon Speed ESI-IT mass spectrometer (Bruker). Five microliters of sample were injected, and a gradient of 5% to 95% acetonitrile (+0.1% formic acid) in water (+0.1% formic acid) was used over 16 min. Mass spectra were recorded in positive ionization mode (m/z 100–1200, cone voltage 4500 V). Data were processed using DataAnalysis (Bruker).

For relative quantification of XTP production, biological triplicates of split intein XtpS and wild-type XtpS samples were analyzed. Peak areas were calculated from the extracted ion chromatogram (EIC) of cyclic XTP and compared to the wild-type. All data are presented as mean ± SD (n = 3 biological replicates) and visualized in GraphPad Prism 10.

#### SDS page and Western blot

*E. coli* cultures were grown in XPP3 medium (10 mL, baffled flask) for 72 h at 20 °C, 130 rpm. Cell pellets were lysed with NEBExpress E. coli lysis reagent (NEB; 66 µL NEBExpress lysis reagent were added to 33 µl pellet), mixed and incubated for 15 min at room temperature. Lysates were centrifuged (20,000 g, 10 min), supernatants stored at −20 °C. For SDS-PAGE, 26 µL lysate was mixed with 10 µL NuPAGE LDS sample buffer (4x, Thermo Fisher) and 4 µL NuPAGE Sample Reducing Agent (10x, Thermo Fisher), heated at 70 °C for 10 min, and centrifuged (20,000 g, 5 min). 20 µL samples were loaded onto 3–8% Tris-Acetate SDS gels (Thermo Fisher) and run for 35 min at 200 V or 1 h at 150 V in Tris-Acetate SDS running buffer using a Mini Gel Tank (Thermo Fisher).

After electrophoresis, proteins were transferred to 0.2 µm nitrocellulose membranes using the Trans-Blot Turbo Transfer System (“HIGH MW” protocol, 1.3 A, 25 V, 10 min, Bio-Rad).

Membranes were blocked for 1 h at room temperature in 3% (w/v) BSA in TBS-T (10 mM Tris, 150 mM NaCl, 0.05% Tween 20, pH 7.5), then incubated overnight at 4 °C with anti-poly-Histidine HRP conjugate (Sigma, #A7058; used 1:1500) and StrepTag II Antibody HRP conjugate (Sigma, #71591; used 1:3000) in 3% BSA/TBS-T. After five 5-min washes in TBS-T, membranes were developed using the Amersham ECL Prime Western Blot Kit (GE Healthcare) and detected on a ChemiDoc MP Imaging System (Bio-Rad).

##### High-throughput transformation and expression of NRPS library

For high-throughput library screening, transformations and NRPS expression and extraction were performed with protocol modifications as outlined below. Plasmid DNA from library constructs was prepared using midi-prep and diluted to ~25 ng/µL. DNA was distributed into 384-well Echo source plates and transferred (0.5 µL per well) into 96-well PCR plates (Eppendorf twin.tec) using the Echo 525 Acoustic Liquid Handler (Beckman Coulter).

Chemically competent *E. coli* DH10B::mtaA cells (20 µL per well) were added manually with a multi-channel pipette and incubated on ice for 30 min. Heat shock was performed using PCR cyclers (1 min at 42 °C, 3 min at 4 °C), followed by the addition of 150 µL SOC medium per well. The plates were sealed with gas-permeable adhesive seals and incubated at 37 °C, 800 rpm for 1.5 h in an Inforce shaker. After recovery, cells were transferred to sterile 96-deep-well plates containing 1.35 mL LB with antibiotics (standard concentrations), sealed, and grown overnight at 37 °C, 800 rpm.

For production, 4 mL XPP3 medium was dispensed into each well of 24-well deep-well plates, and 150 µL Amberlite XAD16N beads were added. Cultures were inoculated automatically with 40 µL overnight culture per well and incubated at 25 °C, 290 rpm for 48 h. After cultivation, supernatant was removed using a fluid aspiration system, and peptides were extracted from the beads with 4 mL methanol:acetonitrile (1:1, v/v) at 25 °C, 290 rpm for 30 min. Plates were centrifuged (4350 g, 20 min), and 100 µL supernatants were transferred to fresh plates for analysis.

##### SPPS

Solid phase peptide synthesis was performed on a Liberty Prime 2.0 automated microwave peptide synthesizer (CEM) using standard Fmoc chemistry. Fourteen peptides were synthesized as chemical standards. pepIN01, 04, 05, 06, and 07 were synthesized on preloaded Fmoc-(Val or Leu)-2-CT resin (0.1 mmol, 1 eq), while pepIN02, 03, 08, 09, 10, 11,

12, 13, and 14 were synthesized on Fmoc-(Val, Leu or Phe)-Wang resin. Synthesis was undergone with a two-stage reaction of deprotection and coupling. Deprotection was done with 25 % pyrrolidine in DMF for 5 min at 90°C. Coupling was conducted twice at 50°C for 10 minutes with the reagents *N,N'*- Diisopropylcarbodiimide (DIC, 2M), Oxyma (0.25M) and *N*-*N*-Diisopropylethylamine (DIPEA, 0.1 M). After synthesis, the resin was washed 5x with DMF, and 5x DCM. Peptides on 2-CT resin were cleaved with HFIP/DCM (20:80) to keep side chain protection groups, while peptide on Wang resin were cleaved with TFA/$H_2O$/TIS (95:2.5:2.5), both for 30 min at 37°C. The cleavage solution was evaporated at the V10 (Biotage) and washed two times with MeOH.

Peptides pepIN01-07 were cyclized head-to-tail cyclized using 5 eq HATU 12 eq DIPEA in 100 mL DMF for 1h at RT. Subsequently side chain protection groups were removed using TFA/$H_2O$/TIS (95:2.5:2.5) for 1h at RT.

Crude peptides were purified using a Agilent 1290 Infinity II Autoscale Preparative LC System with a ZORBAX Eclipse XDB-C18, PrepHT column (AG-977250-102, Agilent) and a solvent system of acetonitrile and water with 0.1% formic acid.

##### Analysis of MS data

MS data were analyzed using the batch processing function of the DataAnalysis software (Bruker). The following script was used together with an Excel sheet, providing sample names in clolumn A and target m/z values in columns E to T.

```
Dim BPC
Dim EIC
Dim areaThreshold
areaThreshold = 50000
Dim EICwidth
EICwidth = 0.01
Dim relAreaFracThreshold
relAreaFracThreshold = 50

' Define the Excel file path
Dim excelFilePath
excelFilePath = "C:\filepath\filename.xlsx"

' Create an instance of Excel
Dim excelApp
Set excelApp = CreateObject("Excel.Application")

' Open the Excel workbook
Dim workbook
Set workbook = excelApp.Workbooks.Open(excelFilePath)
```

```
' Search column A for the cell that matches the current dataset name
Dim sheet
Set sheet = workbook.Sheets(1)
Dim cell
Dim foundCell
Set foundCell = Nothing

For Each cell In sheet.Range("A:A")
    If cell.Value = analysis.name Then
        Set foundCell = cell
        Exit For
    End If
Next

' Check if the matching cell was found
If Not foundCell Is Nothing Then
    Set BPC = CreateObject("DataAnalysis.BPCChromatogramDefinition")
    Set EIC = CreateObject("DataAnalysis.EICChromatogramDefinition")

    ' Set the colors for the chromatograms
    BPC.Color = RGB(144, 144, 144) ' grey
    EIC.Color = RGB(200, 0, 60)    ' red

    ' Add BPC chromatogram
    Analysis.Chromatograms.Clear
    Analysis.Chromatograms.AddChromatogram BPC

    ' Predefined colors
    Dim colors(4)
    colors(0) = RGB(204, 51, 153)   ' purple
    colors(1) = RGB(17, 115, 159)   ' blue
    colors(2) = RGB(153, 102, 51)   ' brown
    colors(3) = RGB(0, 176, 80)     ' green
    colors(4) = RGB(210, 85, 30)    ' orange

    ' Check for target masses in columns E to T
    Dim targetMass
    Dim col
    Dim colorIndex
    For col = 5 To 20 ' Columns E to T
        targetMass = sheet.Cells(foundCell.Row, col).Value
        If Not IsEmpty(targetMass) Then ' Add EIC chromatograms
            Set EIC = CreateObject("DataAnalysis.EICChromatogramDefinition")
            EIC.Range = targetMass
            colorIndex = (col - 5) Mod 5
            EIC.Color = colors(colorIndex)

            EIC.widthleft = EICwidth
            EIC.widthright = EICwidth
            EIC.BackgroundType = daSpectral
            Analysis.Chromatograms.AddChromatogram EIC
            Analysis.Chromatograms(Analysis.Chromatograms.Count).AddRangeSelection
1.0, 15.0, 0, 0
            Analysis.Chromatograms(Analysis.Chromatograms.Count).IntegrateOnly '
Integrate all chromatograms
```

```
            Dim compoundFound
            compoundFound = False

            For num = 1 To Analysis.Compounds.Count
                If Analysis.Compounds(num).Area > areaThreshold _
                    And InStr(Analysis.Compounds(num).Chromatogram, "EIC") > 0 _
                    And Analysis.Compounds(num).RelativeAreaFraction > 
relAreaFracThreshold Then

                    compoundFound = True
                End If
            Next

            ' Color the cell depending whether compound was found
            If compoundFound Then
                sheet.Cells(foundCell.Row, col).Interior.Color = RGB(0, 255, 0) ' 
Green
            Else
                sheet.Cells(foundCell.Row, col).Interior.Color = RGB(255, 0, 0) ' 
Red
            End If

            Analysis.Compounds.Clear ' clear compound list
        End If
    Next
Else
    MsgBox "No matching dataset name found in column A."
    workbook.Close False
    excelApp.Quit
    Set workbook = Nothing
    Set excelApp = Nothing
    WScript.Quit
End If

' Close Excel & save DataAnalysis file
workbook.Close True ' Save changes
excelApp.Quit
Set workbook = Nothing
Set excelApp = Nothing
analysis.save
form.close
```

#### SUPPLEMENTARY TABLES

**Table S1. | *E. coli* strains used in this work.**

| Strain | Genotype | Reference |
|---|---|---|
| *E. coli* DH10B | F_mcrA (mrr-hsdRMS-mcrBC), 80lacZΔ, M15, ΔlacX74 recA1 endA1 araD 139Δ(ara, leu)7697 galU galK λ rpsL (Strr) nupG / - | [2] |
| *E. coli* DH10B::mtaA | DH10B with mtaA from pCK_mtaA Δ entD | [3] |

**Table S2. | *Photorhabdus and Xenorhabdus* straines with respective BGC and natural products used in this work.**

| Strain | | Gene | Locus tag | GenBank | NP, PubChem CID |
|---|---|---|---|---|---|
| *Xenorhabdus nematophila* | ATCC 19061 | *xtpS* | XNC1_2022 | FN667742 | Xenotetrapeptide 145720651 |
| *Photorhabdus laumondii* TT01 | DSM 15139 | *gxpS* | plu3263 | BX470251 | GameXPeptide A 101582538 |
| *Xenorhabdus miraniensis* | DSM 17902 | *ambS* | Xmir_01407 | NITZ01000005 | Ambactin 139583382 |
| *Xenorhabdus indica* | DSM 17382 | *ambS* | Xind_00972 | NKHP01000004 | Ambactin |
| *Xenorhabdus budapestensis* | DSM 16342 | *bicA* | Xbud_01290 | NIBS01000004 | Bicornitin 162862785 |
| *Xenorhabdus innexi* | DSM 16336 | *fitAB* | Xinn_03284, Xinn_03635 | NIBU01000054, NIBU01000077 | Fitayylide |
| *Xenorhabdus indica* | DSM 17382 | *xldS (truncated iszAB)* | | | Xenolindicin A 139586161 |
| *Xenorhabdus indica* | DSM 17382 | *iszAB* | | | Intraszentin |
| *Mortierella alpina* | ATCC 32222 | *mbpA* | - | MT800759 | Malpibaldin A 101582538 |
| *Xenorhabdus doucetiae* | DSM17909 / FRM16 | *prtAB* | XDD1_1906, XDD1_1907+8 | FO704550 | Protegomycin - PRT-1037 |
| *Xenorhabdus mauleonii* | DSM 17908 | *xabABC* | Xmau_03948, Xmau_03947, Xmau_03946 | NITY01000022 | Xenoamicin A |
| *Xenorhabdus stockiae* | DSM 17904 | *xabABC* | Xsto_01028, Xsto_01027, Xsto_01026 | NJAJ01000007 | Xenoamicin A |
| *Xenorhabdus szentirmaii* | DSM 16338 | *xabABC* | Xsze_03586, Xsze_03587, Xsze_03588 | NIBV01000002 | Xenoamicin A |

**Table S3. | Plasmids used as templates in this work.**

| Plasmid | Description | Backbone | Promoter | Resistance | Reference |
|---|---|---|---|---|---|
| pNA104 | Backbone for initiation units | pACYC_ara | araBAD | Cm | [4] |
| pNA15 | Backbone for elongation units | pCOLA_ara | araBAD | Kan | [4] |
| pNA16 | Backbone for termination untis | pCDF_ara | araBAD | Spec | [4] |
| pLS018 | GxpS 2 T Ser indica GxpS TE (-22a) | | | | [5] |
| pLS119 | GxpS XUT 2 modules Xmau (-21a) | | | | [5] |
| pLS126 | GxpS E XdnAB / fitAB XUT (-19a) | | | | [5] |
| pLS135 | GxpS XUT Moritella Phe (-23a) | | | | [5] |

**Table S4. | Plasmids cloned in this work.**

| Plasmid | Fragment name | Forward primer | Reverse primer | Template | Size (bp) | Resistance |
|---|---|---|---|---|---|---|
| pIN01 | fIN001 - insert gp41-1_N | | | synthetic DNA fragment | 417 | Cm |
| | PCR001 | oIN1F | oIN2R | pNA104 | 5223 | |
| pIN02 | fIN006 - insert gp41-1_C | | | synthetic DNA fragment | 255 | Spec |
| | PCR002 | oIN5F | oIN6R | pNA16 | 3371 | |
| pIN03 | fIN002 - insert gp41-8_N | | | synthetic DNA fragment | 420 | Cm |
| | PCR001 | | | | 5223 | |
| pIN04 | fIN007 - insert gp41-8_C | | | synthetic DNA fragment | 279 | Spec |
| | PCR002 | | | | 3371 | |
| pIN05 | fIN003 - insert NrdH-1_N | | | synthetic DNA fragment | 468 | Cm |
| | PCR001 | | | | 5223 | |
| pIN06 | fIN008 - insert NrdJ-1_C | | | synthetic DNA fragment | 264 | Spec |
| | PCR002 | | | | 3371 | |
| pIN07 | fIN004 - insert IMPDH-1_N | | | synthetic DNA fragment | 456 | Cm |
| | PCR001 | | | | 5223 | |
| pIN08 | fIN009 - insert | | | synthetic DNA fragment | 264 | Spec |

| | | | | | | |
|---|---|---|---|---|---|---|
| | IMPDH-1_C | | | | | |
| | PCR002 | | | | 3371 | |
| pIN09 | fIN005 - insert SspGyrB_N | | | synthetic DNA fragment | 492 | Cm |
| | PCR001 | | | | 5223 | |
| pIN10 | fIN010 - insert SspGyrB_C | | | synthetic DNA fragment | 276 | Spec |
| | PCR002 | | | | 3371 | |
| pXtpS | PCR068 | oIN119F | oIN120R | *X. nematophila* ATCC 19061 | 6513 | Kan |
| | PCR069 | oIN121F | oIN122R | *X. nematophila* ATCC 19061 | 5988 | |
| | PCR067 | oIN117F | oIN118R | pNA15 | 3312 | |
| pIN73 | PCR070 | oIN123F | oIN124R | pXtpS | 4956 | Cm |
| | PCR055 | oIN100F | oIN101R | pIN01 | 5259 | |
| pIN74 | PCR071 | oIN123F | oIN125R | pXtpS | 5022 | Cm |
| | PCR055 | | | | 5259 | |
| pIN76 | PCR073 | oIN123F | oIN127R | pXtpS | 5775 | Cm |
| | PCR055 | | | | 5259 | |
| pIN77 | PCR074 | oIN123F | oIN128R | pXtpS | 6582 | Cm |
| | PCR055 | | | | 5259 | |
| pIN78 | PCR075 | oIN129F | oIN135R | pXtpS | 3279 | Spec |
| | PCR080 | oIN136F | oIN130R | pXtpS | 4326 | |
| | PCR056 | oIN102F | oIN103R | pIN02 | 3401 | |
| pIN79 | PCR076 | oIN131F | oIN135R | pXtpS | 3195 | Spec |
| | PCR080 | | | | 4326 | |
| | PCR056 | | | | 3401 | |
| pIN81 | PCR078 | oIN133F | oIN135R | pXtpS | 2442 | Spec |
| | PCR080 | | | | 4326 | |
| | PCR056 | | | | 3401 | |
| pIN82 | PCR079 | oIN134F | oIN135R | pXtpS | 1653 | Spec |
| | PCR080 | | | | 4326 | |
| | PCR056 | | | | 3401 | |
| pIN85 | PCR081 | oIN140F | oIN141R | pXtpS | 4956 | Cm |
| | PCR003 | oIN24F | oIN25R | pIN01 | 5532 | |
| pIN86 | PCR082 | oIN140F | oIN142R | pXtpS | 4956 | Cm |
| | PCR004 | oIN26F | oIN25R | pIN03 | 5535 | |
| pIN87 | PCR083 | oIN140F | oIN143R | pXtpS | 4956 | Cm |
| | PCR005 | oIN27F | oIN25R | pIN05 | 5583 | |
| pIN88 | PCR084 | oIN140F | oIN144R | pXtpS | 4938 | Cm |
| | PCR003 | | | | 5532 | |
| pIN89 | PCR085 | oIN140F | oIN145R | pXtpS | 4938 | Cm |
| | PCR004 | | | | 5535 | |
| pIN90 | PCR086 | oIN140F | oIN146R | pXtpS | 4938 | Cm |
| | PCR005 | | | | 5583 | |
| pIN91 | PCR087 | oIN140F | oIN147R | pXtpS | 5031 | Cm |
| | PCR003 | | | | 5532 | |
| pIN92 | PCR088 | oIN140F | oIN148R | pXtpS | 5031 | Cm |
| | PCR004 | | | | 5535 | |
| pIN93 | PCR089 | oIN140F | oIN149R | pXtpS | 5031 | Cm |
| | PCR005 | | | | 5583 | |
| pIN94 | PCR090 | oIN140F | oIN150R | pXtpS | 5022 | Cm |
| | PCR003 | | | | 5532 | |

| | | | | | | |
|---|---|---|---|---|---|---|
| pIN95 | PCR091 | oIN140F | oIN151R | pXtpS | 5022 | Cm |
| | PCR004 | | | | 5535 | |
| pIN96 | PCR092 | oIN140F | oIN152R | pXtpS | 5022 | Cm |
| | PCR005 | | | | 5583 | |
| pIN97 | PCR093 | oIN140F | oIN153R | pXtpS | 5784 | Cm |
| | PCR003 | | | | 5532 | |
| pIN98 | PCR094 | oIN140F | oIN154R | pXtpS | 5784 | Cm |
| | PCR004 | | | | 5535 | |
| pIN99 | PCR095 | oIN140F | oIN155R | pXtpS | 5784 | Cm |
| | PCR005 | | | | 5583 | |
| pIN100 | PCR096 | oIN140F | oIN156R | pXtpS | 5775 | Cm |
| | PCR003 | | | | 5532 | |
| pIN101 | PCR097 | oIN140F | oIN157R | pXtpS | 5775 | Cm |
| | PCR004 | | | | 5535 | |
| pIN102 | PCR098 | oIN140F | oIN158R | pXtpS | 5775 | Cm |
| | PCR005 | | | | 5583 | |
| pIN103 | PCR099 | oIN140F | oIN159R | pXtpS | 6582 | Cm |
| | PCR003 | oIN24F | oIN25R | pIN01 | 5532 | |
| pIN104 | PCR100 | oIN140F | oIN160R | pXtpS | 6582 | Cm |
| | PCR004 | oIN26F | oIN25R | pIN03 | 5535 | |
| pIN105 | PCR101 | oIN140F | oIN161R | pXtpS | 6582 | Cm |
| | PCR005 | oIN27F | oIN25R | pIN05 | 5583 | |
| pIN106 | PCR102 | oIN140F | oIN162R | pXtpS | 6573 | Cm |
| | PCR003 | | | | 5532 | |
| pIN107 | PCR103 | oIN140F | oIN163R | pXtpS | 6573 | Cm |
| | PCR004 | | | | 5535 | |
| pIN108 | PCR104 | oIN140F | oIN164R | pXtpS | 6573 | Cm |
| | PCR005 | | | | 5583 | |
| pIN109 | PCR105 | oIN165F | oIN135R | pXtpS | 3279 | Spec |
| | PCR080 | oIN136F | oIN130R | pXtpS | 4326 | |
| | PCR030 | oIN75F | oIN76R | pIN02 | 3518 | |
| pIN110 | PCR106 | oIN166F | oIN135R | pXtpS | 3279 | Spec |
| | PCR080 | | | | 4326 | |
| | PCR031 | oIN75F | oIN77R | pIN04 | 3542 | |
| pIN111 | PCR107 | oIN167F | oIN135R | pXtpS | 3279 | Spec |
| | PCR080 | | | | 4326 | |
| | PCR032 | oIN75F | oIN78R | pIN06 | 3527 | |
| pIN112 | PCR108 | oIN168F | oIN135R | pXtpS | 3204 | Spec |
| | PCR080 | | | | 4326 | |
| | PCR030 | | | | 3518 | |
| pIN113 | PCR109 | oIN169F | oIN135R | pXtpS | 3204 | Spec |
| | PCR080 | | | | 4326 | |
| | PCR031 | | | | 3542 | |
| pIN114 | PCR110 | oIN170F | oIN135R | pXtpS | 3204 | Spec |
| | PCR080 | | | | 4326 | |
| | PCR032 | | | | 3527 | |
| pIN115 | PCR111 | oIN171F | oIN135R | pXtpS | 3195 | Spec |
| | PCR080 | | | | 4326 | |
| | PCR030 | | | | 3518 | |
| pIN116 | PCR112 | oIN172F | oIN135R | pXtpS | 3195 | Spec |
| | PCR080 | | | | 4326 | |
| | PCR031 | | | | 3542 | |
| pIN117 | PCR113 | oIN173F | oIN135R | pXtpS | 3195 | Spec |
| | PCR080 | | | | 4326 | |
| | PCR032 | | | | 3527 | |

| | | | | | | | |
|---|---|---|---|---|---|---|---|
| pIN118 | PCR114 | oIN174F | oIN135R | pXtpS | | 2451 | Spec |
| | PCR080 | | | | | 4326 | |
| | PCR030 | | | | | 3518 | |
| pIN119 | PCR115 | oIN175F | oIN135R | pXtpS | | 2451 | Spec |
| | PCR080 | | | | | 4326 | |
| | PCR031 | | | | | 3542 | |
| pIN120 | PCR116 | oIN176F | oIN135R | pXtpS | | 2451 | Spec |
| | PCR080 | | | | | 4326 | |
| | PCR032 | | | | | 3527 | |
| pIN121 | PCR117 | oIN177F | oIN135R | pXtpS | | 2442 | Spec |
| | PCR080 | | | | | 4326 | |
| | PCR030 | | | | | 3518 | |
| pIN122 | PCR118 | oIN178F | oIN135R | pXtpS | | 2442 | Spec |
| | PCR080 | | | | | 4326 | |
| | PCR031 | | | | | 3542 | |
| pIN123 | PCR119 | oIN179F | oIN135R | pXtpS | | 2442 | Spec |
| | PCR080 | | | | | 4326 | |
| | PCR032 | | | | | 3527 | |
| pIN124 | PCR120 | oIN180F | oIN135R | pXtpS | | 1653 | Spec |
| | PCR080 | | | | | 4326 | |
| | PCR030 | | | | | 3518 | |
| pIN125 | PCR121 | oIN181F | oIN135R | pXtpS | | 1653 | Spec |
| | PCR080 | | | | | 4326 | |
| | PCR031 | | | | | 3542 | |
| pIN126 | PCR122 | oIN182F | oIN135R | pXtpS | | 1653 | Spec |
| | PCR080 | | | | | 4326 | |
| | PCR032 | | | | | 3527 | |
| pIN127 | PCR123 | oIN183F | oIN135R | pXtpS | | 1644 | Spec |
| | PCR080 | | | | | 4326 | |
| | PCR030 | | | | | 3518 | |
| pIN128 | PCR124 | oIN184F | oIN135R | pXtpS | | 1644 | Spec |
| | PCR080 | | | | | 4326 | |
| | PCR031 | | | | | 3542 | |
| pIN129 | PCR125 | oIN185F | oIN135R | pXtpS | | 1644 | Spec |
| | PCR080 | | | | | 4326 | |
| | PCR032 | | | | | 3527 | |
| pIN146 | PCR136 | oIN208F | oIN209R | pIN66 | | 6660 | Kan |
| | PCR137 | oIN210F | oIN211R | pIN70 | | 9141 | |
| | PCR067 | oIN117F | oIN118R | pNA15 | | 3312 | |
| pIN147 | PCR138 | oIN208F | oIN212R | pIN77 | | 6603 | Kan |
| | PCR139 | oIN213F | oIN135R | pIN82 | | 1638 | |
| | PCR140 | oIN136F | oIN211R | pIN82 | | 4356 | |
| | PCR067 | oIN117F | oIN118R | pNA15 | | 3312 | |
| pIN155 | PCR156 | oIN233F | oIN234R | pXtpS | | 3489 | Spec |
| | PCR056 | | | | | 3401 | |
| pIN156 | PCR157 | oIN235F | oIN236R | pXtpS | | 3318 | Kan |
| | PCR067 | | | | | 3312 | |
| pIN160 | PCR160 | oIN140F | oIN249R | pXtpS | | 4956 | Cm |
| | PCR006 | oIN28F | oIN25R | pIN07 | | 5571 | |
| pIN161 | PCR161 | oIN140F | oIN250R | pXtpS | | 4956 | Cm |
| | PCR007 | oIN29F | oIN25R | pIN09 | | 5607 | |
| pIN162 | PCR162 | oIN140F | oIN251R | pXtpS | | 4938 | Cm |
| | PCR006 | | | | | 5571 | |
| pIN163 | PCR163 | oIN140F | oIN252R | pXtpS | | 4938 | Cm |
| | PCR007 | | | | | 5607 | |

| | | | | | | |
|---|---|---|---|---|---|---|
| pIN164 | PCR164 | oIN140F | oIN253R | pXtpS | 5031 | Cm |
| | PCR006 | | | | 5571 | |
| pIN165 | PCR165 | oIN140F | oIN254R | pXtpS | 5031 | Cm |
| | PCR007 | | | | 5607 | |
| pIN166 | PCR166 | oIN140F | oIN255R | pXtpS | 5022 | Cm |
| | PCR006 | | | | 5571 | |
| pIN167 | PCR167 | oIN140F | oIN256R | pXtpS | 5022 | Cm |
| | PCR007 | | | | 5607 | |
| pIN168 | PCR168 | oIN140F | oIN257R | pXtpS | 5784 | Cm |
| | PCR006 | | | | 5571 | |
| pIN169 | PCR169 | oIN140F | oIN258R | pXtpS | 5784 | Cm |
| | PCR007 | | | | 5607 | |
| pIN170 | PCR170 | oIN140F | oIN259R | pXtpS | 5775 | Cm |
| | PCR006 | | | | 5571 | |
| pIN171 | PCR171 | oIN140F | oIN260R | pXtpS | 5775 | Cm |
| | PCR007 | | | | 5607 | |
| pIN172 | PCR172 | oIN140F | oIN261R | pXtpS | 6582 | Cm |
| | PCR006 | | | | 5571 | |
| pIN173 | PCR173 | oIN140F | oIN262R | pXtpS | 6582 | Cm |
| | PCR007 | | | | 5607 | |
| pIN174 | PCR174 | oIN140F | oIN263R | pXtpS | 6573 | Cm |
| | PCR006 | | | | 5571 | |
| pIN175 | PCR175 | oIN140F | oIN264R | pXtpS | 6573 | Cm |
| | PCR007 | | | | 5607 | |
| pIN176 | PCR176 | oIN265F | oIN135R | pXtpS | 3279 | Spec |
| | PCR080 | oIN136F | oIN130R | pXtpS | 4326 | |
| | PCR033 | oIN75F | oIN79R | pIN08 | 3527 | |
| pIN177 | PCR177 | oIN266F | oIN135R | pXtpS | 3279 | Spec |
| | PCR080 | | | | 4326 | |
| | PCR034 | oIN75F | oIN80R | pIN10 | 3539 | |
| pIN178 | PCR178 | oIN267F | oIN135R | pXtpS | 3204 | Spec |
| | PCR080 | | | | 4326 | |
| | PCR033 | | | | 3527 | |
| pIN179 | PCR179 | oIN268F | oIN135R | pXtpS | 3204 | Spec |
| | PCR080 | | | | 4326 | |
| | PCR034 | | | | 3539 | |
| pIN180 | PCR180 | oIN269F | oIN135R | pXtpS | 3195 | Spec |
| | PCR080 | | | | 4326 | |
| | PCR033 | | | | 3527 | |
| pIN181 | PCR181 | oIN270F | oIN135R | pXtpS | 3195 | Spec |
| | PCR080 | | | | 4326 | |
| | PCR034 | | | | 3539 | |
| pIN182 | PCR182 | oIN271F | oIN135R | pXtpS | 2451 | Spec |
| | PCR080 | | | | 4326 | |
| | PCR033 | | | | 3527 | |
| pIN183 | PCR183 | oIN272F | oIN135R | pXtpS | 2451 | Spec |
| | PCR080 | | | | 4326 | |
| | PCR034 | | | | 3539 | |
| pIN184 | PCR184 | oIN273F | oIN135R | pXtpS | 2442 | Spec |
| | PCR080 | | | | 4326 | |
| | PCR033 | | | | 3527 | |
| pIN185 | PCR185 | oIN274F | oIN135R | pXtpS | 2442 | Spec |
| | PCR080 | | | | 4326 | |
| | PCR034 | | | | 3539 | |

| | | | | | | |
|---|---|---|---|---|---|---|
| pIN186 | PCR186 | oIN275F | oIN135R | pXtpS | 1653 | Spec |
| | PCR080 | | | | 4326 | |
| | PCR033 | | | | 3527 | |
| pIN187 | PCR187 | oIN276F | oIN135R | pXtpS | 1653 | Spec |
| | PCR080 | | | | 4326 | |
| | PCR034 | | | | 3539 | |
| pIN188 | PCR188 | oIN277F | oIN135R | pXtpS | 1644 | Spec |
| | PCR080 | | | | 4326 | |
| | PCR033 | | | | 3527 | |
| pIN189 | PCR189 | oIN278F | oIN135R | pXtpS | 1644 | Spec |
| | PCR080 | | | | 4326 | |
| | PCR034 | | | | 3539 | |
| pIN198 | PCR214 | oIN320F | oIN321R | pIN04 | 299 | Kan |
| | PCR215 | oIN322F | oIN323R | pIN05 | 354 | |
| | PCR212 | oIN117F | oIN319R | pNA15 | 428 | |
| | PCR213 | oIN318F | oIN118R | pNA15 | 2911 | |
| pIN199 | PCR216 | oIN324F | oIN325R | pIN02 | 275 | Kan |
| | PCR217 | oIN326F | oIN327R | pIN09 | 378 | |
| | PCR212 | oIN117F | oIN319R | pNA15 | 428 | |
| | PCR213 | oIN318F | oIN118R | pNA15 | 2911 | |
| pIN200 | PCR220 | oIN332F | oIN333R | pXtpS | 3243 | Kan |
| | PCR218 | oIN328F | oIN329R | pIN198 | 3777 | |
| pIN201 | PCR221 | oIN334F | oIN335R | pXtpS | 3243 | Kan |
| | PCR218 | oIN328F | oIN329R | pIN198 | 3777 | |
| pIN202 | PCR222 | oIN336F | oIN337R | pXtpS | 3318 | Kan |
| | PCR219 | oIN330F | oIN331R | pIN199 | 3777 | |
| pIN203 | PCR223 | oIN338F | oIN339R | pXtpS | 3280 | Kan |
| | PCR219 | oIN330F | oIN331R | pIN199 | 3300 | |
| pIN204 | PCR224 | oIN366F | oIN341R | pXtpS | 4392 | Spec |
| | PCR032 | oIN75F | oIN78R | pIN06 | 3527 | |
| pIN205 | PCR225 | oIN342F | oIN341R | pXtpS | 4317 | Spec |
| | PCR032 | oIN75F | oIN78R | pIN06 | 3527 | |
| pIN206 | PCR226 | oIN343F | oIN341R | pXtpS | 3489 | Spec |
| | PCR034 | oIN75F | oIN80R | pIN10 | 3539 | |
| pIN207 | PCR227 | oIN344F | oIN341R | pXtpS | 2709 | Spec |
| | PCR034 | oIN75F | oIN80R | pIN10 | 3539 | |
| pIN208 | PCR228 | oIN345F | oIN346R | pXtpS | 3300 | Kan |
| | PCR067 | oIN117F | oIN118R | pNA15 | 3312 | |
| pIN209 | PCR229 | oIN347F | oIN341R | pXtpS | 2709 | Spec |
| | PCR056 | oIN102F | oIN103R | pIN02 | 3401 | |
| pIN24 | PCR009 | oIN30F | oIN32R | *P. luminescens subsp. laumondii TTO1* | 5001 | Cm |
| | PCR004 | oIN26F | oIN25R | pIN03 | 5535 | |
| pIN241 | PCR280 | oIN432F | oIN433R | *X. budapestensis DSM 16342* | 5631 | Cm |
| | PCR004 | | | | 5535 | |
| pIN243 | PCR282 | oIN436F | oIN437R | *X. indica* DSM 17382 | 6261 | Cm |
| | PCR004 | | | | 5535 | |
| pIN244 | PCR283 | oIN438F | oIN439R | *X. stockiae* DSM 17904 | 6315 | Cm |
| | PCR004 | | | | 5535 | |
| pIN263 | PCR315 | oIN460F | oIN461R | *X. innexi* DSM 16336 | 6147 | Cm |
| | PCR004 | | | | 5535 | |
| pIN264 | PCR316 | oIN462F | oIN463R | *X. szentirmaii strain* DSM 16338 | 6378 | Cm |
| | PCR004 | | | | 5535 | |
| pIN237 | PCR274 | oIN424F | oIN425R | pLS018 | 3174 | Kan |

| | | | | | | |
|---|---|---|---|---|---|---|
| | PCR218 | oIN328F | oIN329R | pIN198 | 3777 | |
| pIN238 | PCR275 | oIN426F | oIN427R | pLS119 | 6617 | Kan |
| | PCR218 | | | | 3777 | |
| pIN239 | PCR276 | oIN428F | oIN429R | pLS126 | 4536 | Kan |
| | PCR218 | | | | 3777 | |
| pIN240 | PCR277 | oIN430F | oIN431R | pLS135 | 3223 | Kan |
| | PCR218 | | | | 3777 | |
| pIN246 | PCR285 | oIN442F | oIN443R | *X. miraniensis DSM 17902* | 6462 | Kan |
| | PCR218 | | | | 3777 | |
| pIN247 | PCR286 | oIN444F | oIN445R | *X. indica* DSM 17382 | 6573 | Kan |
| | PCR218 | | | | 3777 | |
| pIN269 | PCR321 | oIN472F | oIN473R | *X. innexi* DSM 16336 | 7428 | Kan |
| | PCR218 | | | | 3777 | |
| pIN285 | PCR347 | oIN524F | oIN525R | *P. luminescens subsp. laumondii* TTO1 | 6396 | Kan |
| | PCR218 | | | | 3777 | |
| pIN45 | PCR037 | oIN84F | oIN82R | *P. luminescens subsp. laumondii* TTO1 | 10764 | Spec |
| | PCR032 | oIN75F | oIN78R | pIN06 | 3527 | |
| pIN235 | PCR272 | oIN420F-2 | oIN421R | *X. doucetiae* DSM17909 / FRM16 | 8660 | Spec |
| | PCR032 | | | | 3527 | |
| pIN275 | PCR327 | oIN484F | oIN485R | *X. miraniensis* DSM 17902 | 3327 | Spec |
| | PCR328 | oIN486F | oIN487R | pIN204 | 4619 | |
| pIN286 | PCR348 | oIN526F | oIN527R | *P. luminescens subsp. laumondii* TTO1 | 4368 | Spec |
| | PCR032 | | | | 3527 | |

**Table S5. | Oligonucleotides used in this work.**

| Name | Sequence 5' → 3' | Length (bp) |
|---|---|---|
| oIN1F | TGACAATTAATCATCGGCTCGTATAATG | 28 |
| oIN2R | CATGGAATTCCTCCTGTTAGCCC | 23 |
| oIN5F | TGACAATTAATCATCGGCTCGTATAATG | 28 |
| oIN6R | CATGGAATTCCTCCTGTTAGCCC | 23 |
| oIN24F | AGCGGCTATTGTCTGGATCTG | 21 |
| oIN25R | GCTGCCGCTACCTTTCTCAAAC | 22 |
| oIN26F | CTGAACCGCTGTCTGAGCCTG | 21 |
| oIN27F | AACCCTTGTTGTCTGGTTGGTAGC | 24 |
| oIN28F | GGTGGAGGATGTTTTGTTCCGG | 22 |
| oIN29F | GCTGGGGGTTGTTTTTCTGGAG | 22 |
| oIN30F | CATCCGCAGTTTGAGAAAGGTAGCGGCAGCAAAGATAGCATGGCTAAAAAGGAAATTATCTTTG | 64 |
| oIN32R | CATGGTATCCAGGCTCAGACAGCGGTTCAGAATTTGGCGAGCAAAAGCATCCTCTC | 56 |
| oIN75F | GGTAGCGGCAGCCATCATC | 19 |
| oIN76R | CGAGCTTGAATTATGGGTCAGAATATC | 27 |
| oIN77R | CACTGCAGAATTATGGGTCAGAATAC | 26 |
| oIN78R | AATTTCGCTATTATGCACCAGAATATCATTG | 31 |
| oIN79R | GCAGATGCTATTATGAACAACGGTG | 25 |
| oIN80R | CTTCGCTGAGTTATGGACAAACAC | 24 |

| | | |
|---|---|---|
| oIN82R | ATGATGATGATGATGATGGCTGCCGCTACCCAGCGCCTCCGCTTCACAATTC | 52 |
| oIN84F | AATGATATTCTGGTGCATAATAGCGAAATTTATGTCGCGCCACAGGGAG | 49 |
| oIN100F | TGACAATTAATCATCGGCTCGTATAATG | 28 |
| oIN101R | GCTGCCGCTACCTTTCTCAAACTG | 24 |
| oIN102F | GGTAGCGGCAGCCATCATCATC | 22 |
| oIN103R | CATGGAATTCCTCCTGTTAGCCC | 23 |
| oIN117F | TGACAATTAATCATCGGCTCGTATAATGTG | 30 |
| oIN118R | CATGGAATTCCTCCTGTTAGCCC | 23 |
| oIN119F | TTTTTTTGGGCTAACAGGAGGAATTCCATGAAAGATAGCATGGCTAAAAAGGGAATTATCTTTGAC | 66 |
| oIN120R | AATATCAATAACCGCAACAGGTTGTTGAGG | 30 |
| oIN121F | CCTCAACAACCTGTTGCGGTTATTGATATT | 30 |
| oIN122R | CACATTATACGAGCCGATGATTAATTGTCACAGCGCCTCCACTTCGCAATTCATTG | 56 |
| oIN123F | CATCCGCAGTTTGAGAAAGGTAGCGGCAGCAAAGATAGCATGGCTAAAAAGGGAATTATCTTTGAC | 66 |
| oIN124R | CACATTATACGAGCCGATGATTAATTGTCAAATCTGGCGGGCGAAAGCCTC | 51 |
| oIN125R | CACATTATACGAGCCGATGATTAATTGTCAAATACCCAGTAATTCACGCCAGATAGCC | 58 |
| oIN127R | CACATTATACGAGCCGATGATTAATTGTCACCGCTGCCAGGCAGCATAATC | 51 |
| oIN128R | CACATTATACGAGCCGATGATTAATTGTCACCAGGTTTTTAACAACAATGTGCGTTCC | 58 |
| oIN129F | TTTTTTTGGGCTAACAGGAGGAATTCCATGTATGTTGCGCCACAAGGAGAAATGG | 55 |
| oIN130R | ATGATGATGATGATGATGGCTGCCGCTACCCAGCGCCTCCACTTCGCAATTC | 52 |
| oIN131F | TTTTTTTGGGCTAACAGGAGGAATTCCATGGATAATTTCTTTTCCTTGGGCGGACATTC | 59 |
| oIN133F | TTTTTTTGGGCTAACAGGAGGAATTCCATGGGCGTGCAGGTGCAGAGTG | 49 |
| oIN134F | TTTTTTTGGGCTAACAGGAGGAATTCCATGAATGCCACTGAAACCGTGTATCCTG | 55 |
| oIN135R | AATATTCAGTAATTCGCACCAGATAGCCGC | 30 |
| oIN136F | GCGGCTATCTGGTGCGAATTACTGAATATT | 30 |
| oIN140F | CATCCGCAGTTTGAGAAAGGTAGCGGCAGCAAAGATAGCATGGCTAAAAAGGGAATTATCTTTGACG | 67 |
| oIN141R | CTGGGTTTTCAGATCCAGACAATAGCCGCTAATCTGGCGGGCGAAAGCCTCTTCC | 55 |
| oIN142R | CATGGTATCCAGGCTCAGACAGCGGTTCAGAATCTGGCGGGCGAAAGCCTCTTCC | 55 |
| oIN143R | TTCGCTGCTACCAACCAGACAACAAGGGTTAATCTGGCGGGCGAAAGCCTCTTCC | 55 |
| oIN144R | CTGGGTTTTCAGATCCAGACAATAGCCGCTCTCTTCCCCCGGCGCGGGCAATGCC | 55 |
| oIN145R | CATGGTATCCAGGCTCAGACAGCGGTTCAGCTCTTCCCCCGGCGCGGGCAATGCC | 55 |
| oIN146R | TTCGCTGCTACCAACCAGACAACAAGGGTTCTCTTCCCCCGGCGCGGGCAATGCC | 55 |
| oIN147R | CTGGGTTTTCAGATCCAGACAATAGCCGCTGATCTGTTCAATACCCAGTAATTCACGCCAGATAGCCGCTATTGCGATTTCCATT | 85 |
| oIN148R | CATGGTATCCAGGCTCAGACAGCGGTTCAGGATCTGTTCAATACCCAGTAATTCACGCCAGATAGCCGCTATTGCGATTTCCATT | 85 |

| | | |
|---|---|---|
| oIN149R | TTCGCTGCTACCAACCAGACAACAAGGGTTGATCTGTTCAATACCCAG TAATTCACGCCAGATAGCCGCTATTGCGATTTCCATT | 85 |
| oIN150R | CTGGGTTTTCAGATCCAGACAATAGCCGCTAATACCCAGTAATTCACG CCAGATAGCCGCTATTGCGATTTCCATT | 76 |
| oIN151R | CATGGTATCCAGGCTCAGACAGCGGTTCAGAATACCCAGTAATTCACG CCAGATAGCCGCTATTGCGATTTCCATT | 76 |
| oIN152R | TTCGCTGCTACCAACCAGACAACAAGGGTTAATACCCAGTAATTCACG CCAGATAGCCGCTATTGCGATTTCCATT | 76 |
| oIN153R | CTGGGTTTTCAGATCCAGACAATAGCCGCTAAATACCTGCCGCTGCCA GGCAGC | 54 |
| oIN154R | CATGGTATCCAGGCTCAGACAGCGGTTCAGAAATACCTGCCGCTGCCA GGCAGC | 54 |
| oIN155R | TTCGCTGCTACCAACCAGACAACAAGGGTTAAATACCTGCCGCTGCCA GGCAGC | 54 |
| oIN156R | CTGGGTTTTCAGATCCAGACAATAGCCGCTCCGCTGCCAGGCAGCATA ATCAGG | 54 |
| oIN157R | CATGGTATCCAGGCTCAGACAGCGGTTCAGCCGCTGCCAGGCAGCAT AATCAGG | 54 |
| oIN158R | TTCGCTGCTACCAACCAGACAACAAGGGTTCCGCTGCCAGGCAGCATA ATCAGG | 54 |
| oIN159R | CTGGGTTTTCAGATCCAGACAATAGCCGCTCCAGGTTTTTAACAACAAT GTGCGTTCCGTGGACG | 65 |
| oIN160R | CATGGTATCCAGGCTCAGACAGCGGTTCAGCCAGGTTTTTAACAACAA TGTGCGTTCCGTGGACG | 65 |
| oIN161R | TTCGCTGCTACCAACCAGACAACAAGGGTTCCAGGTTTTTAACAACAAT GTGCGTTCCGTGGACG | 65 |
| oIN162R | CTGGGTTTTCAGATCCAGACAATAGCCGCTTAACAACAATGTGCGTTC CGTGGACGACAAAATA | 64 |
| oIN163R | CATGGTATCCAGGCTCAGACAGCGGTTCAGTAACAACAATGTGCGTTC CGTGGACGACAAAATAT | 65 |
| oIN164R | TTCGCTGCTACCAACCAGACAACAAGGGTTTAACAACAATGTGCGTTC CGTGGACGACAAAATAT | 65 |
| oIN165F | AATGATATTCTGACCCATAATTCAAGCTCGTATGTTGCGCCACAAGGAG AAATGGAAATCG | 61 |
| oIN166F | AATGGTATTCTGACCCATAATTCTGCAGTGTATGTTGCGCCACAAGGA GAAATGGAAATCG | 61 |
| oIN167F | AATGATATTCTGGTGCATAATAGCGAAATTTATGTTGCGCCACAAGGAG AAATGGAAATCG | 61 |
| oIN168F | AATGATATTCTGACCCATAATTCAAGCTCGGGCCGGTATGATAATTTCT TTTCCTTGGG | 59 |
| oIN169F | AATGGTATTCTGACCCATAATTCTGCAGTGGGCCGGTATGATAATTTCT TTTCCTTGGG | 59 |
| oIN170F | AATGATATTCTGGTGCATAATAGCGAAATTGGCCGGTATGATAATTTCT TTTCCTTGGG | 59 |
| oIN171F | AATGATATTCTGACCCATAATTCAAGCTCGGATAATTTCTTTTCCTTGG GCGGACATTCTC | 61 |
| oIN172F | AATGGTATTCTGACCCATAATTCTGCAGTGGATAATTTCTTTTCCTTGG GCGGACATTCTC | 61 |
| oIN173F | AATGATATTCTGGTGCATAATAGCGAAATTGATAATTTCTTTTCCTTGGG CGGACATTCTC | 61 |
| oIN174F | AATGATATTCTGACCCATAATTCAAGCTCGTCTGATGAAGGCGTGCAG GTGCAG | 54 |
| oIN175F | AATGGTATTCTGACCCATAATTCTGCAGTGTCTGATGAAGGCGTGCAG GTGCAG | 54 |
| oIN176F | AATGATATTCTGGTGCATAATAGCGAAATTTCTGATGAAGGCGTGCAG GTGCAG | 54 |

| | | |
|---|---|---|
| oIN177F | AATGATATTCTGACCCATAATTCAAGCTCGGGCGTGCAGGTGCAGAGT GATTA | 53 |
| oIN178F | AATGGTATTCTGACCCATAATTCTGCAGTGGGCGTGCAGGTGCAGAGT GATTA | 53 |
| oIN179F | AATGATATTCTGGTGCATAATAGCGAAATTGGCGTGCAGGTGCAGAGT GATTA | 53 |
| oIN180F | AATGATATTCTGACCCATAATTCAAGCTCGAATGCCACTGAAACCGTGT ATCCTGAATC | 59 |
| oIN181F | AATGGTATTCTGACCCATAATTCTGCAGTGAATGCCACTGAAACCGTGT ATCCTGAATC | 59 |
| oIN182F | AATGATATTCTGGTGCATAATAGCGAAATTAATGCCACTGAAACCGTGT ATCCTGAATC | 59 |
| oIN183F | AATGATATTCTGACCCATAATTCAAGCTCGGAAACCGTGTATCCTGAAT CGTTATGTATTCATC | 64 |
| oIN184F | AATGGTATTCTGACCCATAATTCTGCAGTGGAAACCGTGTATCCTGAAT CGTTATGTATTCATC | 64 |
| oIN185F | AATGATATTCTGGTGCATAATAGCGAAATTGAAACCGTGTATCCTGAAT CGTTATGTATTCATC | 64 |
| oIN208F | TTTTTTTGGGCTAACAGGAGGAATTCCATGTGGAGCCATCCGCAGTTT GAGAAAGG | 56 |
| oIN209R | TTCTTCGGTCGCATTCCAATTTTCCAGTAATAACTCCCGCTCAGA | 45 |
| oIN210F | TTACTGGAAAATTGGAATGCGACCGAAGAACCGTATCCTACT | 42 |
| oIN211R | CACATTATACGAGCCGATGATTAATTGTCAATGATGATGATGATGATGG CTGCCGCT | 57 |
| oIN212R | GGTTTCAGTGGCATTCCAGGTTTTTAACAACAATGTGCGTTCCG | 44 |
| oIN213F | TTGTTAAAAACCTGGAATGCCACTGAAACCGTGTATCCTGAATCG | 45 |
| oIN233F | TTTTTTTGGGCTAACAGGAGGAATTCCATGAGTCAGGCAGAACATACC CGTTTCTTTACCGATATGTTGGC | 71 |
| oIN234R | ATGATGATGATGATGATGGCTGCCGCTACCCAGCGCCTCCACTTCGCA ATTCATTGCCAATGC | 63 |
| oIN235F | TTTTTTTGGGCTAACAGGAGGAATTCCATGTCTGATGAAGGCGTGCAG GTGCAGAGTGATTACTGGC | 67 |
| oIN236R | CACATTATACGAGCCGATGATTAATTGTCAGACACCCTGCCGAGCCTG TGCCACGAGATGG | 61 |
| oIN249R | CAGTGTACCCGGAACAAAACATCCTCCACCAATCTGGCGGGCGAAAG CCTCTTCC | 55 |
| oIN250R | TAATGTATCTCCAGAAAAACAACCCCCAGCAATCTGGCGGGCGAAAGC CTCTTCC | 55 |
| oIN251R | CAGTGTACCCGGAACAAAACATCCTCCACCCTCTTCCCCCGGCGCGG GCAATGCC | 55 |
| oIN252R | TAATGTATCTCCAGAAAAACAACCCCCAGCCTCTTCCCCCGGCGCGGG CAATGCC | 55 |
| oIN253R | CAGTGTACCCGGAACAAAACATCCTCCACCGATCTGTTCAATACCCAG TAATTCACGCCAGATAGCCGCTATTGCGATTTCCATT | 85 |
| oIN254R | TAATGTATCTCCAGAAAAACAACCCCCAGCGATCTGTTCAATACCCAGT AATTCACGCCAGATAGCCGCTATTGCGATTTCCATT | 85 |
| oIN255R | CAGTGTACCCGGAACAAAACATCCTCCACCAATACCCAGTAATTCACG CCAGATAGCCGCTATTGCGATTTCCATT | 76 |
| oIN256R | TAATGTATCTCCAGAAAAACAACCCCCAGCAATACCCAGTAATTCACGC CAGATAGCCGCTATTGCGATTTCCATT | 76 |
| oIN257R | CAGTGTACCCGGAACAAAACATCCTCCACCAAATACCTGCCGCTGCCA GGCAGC | 54 |
| oIN258R | TAATGTATCTCCAGAAAAACAACCCCCAGCAAATACCTGCCGCTGCCA GGCAGC | 54 |
| oIN259R | CAGTGTACCCGGAACAAAACATCCTCCACCCCGCTGCCAGGCAGCAT AATCAGG | 54 |

| | | |
|---|---|---|
| oIN260R | TAATGTATCTCCAGAAAAACAACCCCCAGCCCGCTGCCAGGCAGCATAATCAGG | 54 |
| oIN261R | CAGTGTACCCGGAACAAAACATCCTCCACCCCAGGTTTTTAACAACAATGTGCGTTCCGTGGACG | 65 |
| oIN262R | TAATGTATCTCCAGAAAAACAACCCCCAGCCCAGGTTTTTAACAACAATGTGCGTTCCGTGGACG | 65 |
| oIN263R | CAGTGTACCCGGAACAAAACATCCTCCACCTAACAACAATGTGCGTTCCGTGGACGACAAAATA | 64 |
| oIN264R | TAATGTATCTCCAGAAAAACAACCCCCAGCTAACAACAATGTGCGTTCCGTGGACGACAAAATA | 64 |
| oIN265F | CGTGGCACCGTTGTTCATAATAGCATCTGCTATGTTGCGCCACAAGGAGAAATGGAAATCG | 61 |
| oIN266F | AGCGGAGTGTTTGTCCATAACTCAGCGAAGTATGTTGCGCCACAAGGAGAAATGGAAATCG | 61 |
| oIN267F | CGTGGCACCGTTGTTCATAATAGCATCTGCGGCCGGTATGATAATTTCTTTTCCTTGGG | 59 |
| oIN268F | AGCGGAGTGTTTGTCCATAACTCAGCGAAGGGCCGGTATGATAATTTCTTTTCCTTGGG | 59 |
| oIN269F | CGTGGCACCGTTGTTCATAATAGCATCTGCGATAATTTCTTTTCCTTGGGCGGACATTCTC | 61 |
| oIN270F | AGCGGAGTGTTTGTCCATAACTCAGCGAAGGATAATTTCTTTTCCTTGGGCGGACATTCTC | 61 |
| oIN271F | CGTGGCACCGTTGTTCATAATAGCATCTGCTCTGATGAAGGCGTGCAGGTGCAG | 54 |
| oIN272F | AGCGGAGTGTTTGTCCATAACTCAGCGAAGTCTGATGAAGGCGTGCAGGTGCAG | 54 |
| oIN273F | CGTGGCACCGTTGTTCATAATAGCATCTGCGGCGTGCAGGTGCAGAGTGATTA | 53 |
| oIN274F | AGCGGAGTGTTTGTCCATAACTCAGCGAAGGGCGTGCAGGTGCAGAGTGATTA | 53 |
| oIN275F | CGTGGCACCGTTGTTCATAATAGCATCTGCAATGCCACTGAAACCGTGTATCCTGAATC | 59 |
| oIN276F | AGCGGAGTGTTTGTCCATAACTCAGCGAAGAATGCCACTGAAACCGTGTATCCTGAATC | 59 |
| oIN277F | CGTGGCACCGTTGTTCATAATAGCATCTGCGAAACCGTGTATCCTGAATCGTTATGTATTCATC | 64 |
| oIN278F | AGCGGAGTGTTTGTCCATAACTCAGCGAAGGAAACCGTGTATCCTGAATCGTTATGTATTCATC | 64 |
| oIN318F | GCTGACGACCGGGATTCCGCAAGTGGC | 27 |
| oIN319R | GCCACTTGCGGAATCCCGGTCGTCAGC | 27 |
| oIN320F | TTTTTTTGGGCTAACAGGAGGAATTCCATGTG | 32 |
| oIN321R | TTCGCTGCTACCAACCAGACAACAAGGGTTAGAGACCGTAGCCACCAAGTTGGTG | 55 |
| oIN322F | AACCCTTGTTGTCTGGTTGGTAGCAGC | 27 |
| oIN323R | CACATTATACGAGCCGATGATTAATTGTCAAATTGCAACCACCAGTTCATCATCATCGG | 59 |
| oIN324F | TTTTTTTGGGCTAACAGGAGGAATTCCATGATG | 33 |
| oIN325R | TAATGTATCTCCAGAAAAACAACCCCCAGCAGAGACCGTAGCCACCAAGTTGGTG | 55 |
| oIN326F | GCTGGGGGTTGTTTTTCTGGAGATACATTAG | 31 |
| oIN327R | CACATTATACGAGCCGATGATTAATTGTCACGCATGACCAGAATCTTCCGTAGTCG | 56 |
| oIN328F | AACCCTTGTTGTCTGGTTGGTAGCAGC | 27 |
| oIN329R | CACTGCAGAATTATGGGTCAGAATACCATTGG | 32 |
| oIN330F | GCTGGGGGTTGTTTTTCTGGAGATACATTAG | 31 |
| oIN331R | CGAGCTTGAATTATGGGTCAGAATATCATTGGC | 33 |

| | | |
|---|---|---|
| oIN332F | AATGGTATTCTGACCCATAATTCTGCAGTGTATGTTGCGCCACAAGGAGAAATGGAAATCG | 61 |
| oIN333R | TTCGCTGCTACCAACCAGACAACAAGGGTTGACCTGCCGGGCAAACGCCTCATTG | 55 |
| oIN334F | AATGGTATTCTGACCCATAATTCTGCAGTGGGCCGGTATGATAATTTCTTTTCCTTGGGC | 60 |
| oIN335R | TTCGCTGCTACCAACCAGACAACAAGGGTTAATTCGCTCAATATTCAGTAATTCGCACCAGATAGC | 66 |
| oIN336F | AATGATATTCTGACCCATAATTCAAGCTCGTCTGATGAAGGCGTGCAGGTGCAGAGTGATTAC | 63 |
| oIN337R | TAATGTATCTCCAGAAAAACAACCCCCAGCGACACCCTGCCGAGCCTGTGCCACGAG | 57 |
| oIN338F | AATGATATTCTGACCCATAATTCAAGCTCGAATGCCACTGAAACCGTGTATCCTGAATCGTTATG | 65 |
| oIN339R | TAATGTATCTCCAGAAAAACAACCCCCAGCCCAGGTTTTTAACAACAGTGTGCGTTCAGTTTCCGGCA | 68 |
| oIN341R | ATGATGATGATGATGATGGCTGCCGCTACCCAGCGCCTCCACTTCGCAATTCATTGCC | 58 |
| oIN342F | AATAGCGAAATTGGCCGACATGACAGTTTCTTTGC | 35 |
| oIN343F | AGCGGAGTGTTTGTCCATAACTCAGCGAAGAGTCAGGCAGAACATACCCGTTTCTTTACC | 60 |
| oIN344F | AGCGGAGTGTTTGTCCATAACTCAGCGAAGAATGCCACTGAAACCGTATATCCTGAATCGTTATG | 65 |
| oIN345F | TTTTTTTGGGCTAACAGGAGGAATTCCATGAATGCCACTGAAACCGTGTATCCTGAATCGTTATG | 65 |
| oIN346R | CACATTATACGAGCCGATGATTAATTGTCACCAGGTTTTTAACAACAGTGTGCGTTCAGTTTCCGGCA | 68 |
| oIN347F | TTTTTTTGGGCTAACAGGAGGAATTCCATGAATGCCACTGAAACCGTATATCCTGAATCGTTATG | 65 |
| oIN366F | AATGATATTCTGGTGCATAATAGCGAAATTTATGCGGCTCCGCAGGGGGAAACCGAAATAGCATTAGCGGCTATCTGGTGCG | 82 |
| oIN420F-2 | TATTCTGGTGCATAATAGCGAAATTTTTATCGCACCGCGCACGGAGCTGG | 50 |
| oIN421R | GATGATGATGATGGCTGCCGCTACCTTAATGATTATCCTTAATTGACAAGGTATTCATTATTTCTGTACTGAT | 73 |
| oIN424F | TATTCTGACCCATAATTCTGCAGTGGAAGCGCCCATTGGCAAATT | |
| oIN425R | GCTACCAACCAGACAACAAGGGTTATAGCCACGTGTAACAACCGCTGA | |
| oIN426F | TATTCTGACCCATAATTCTGCAGTGCAAGCGCCGGAAAGCCCAAT | |
| oIN427R | TGCTACCAACCAGACAACAAGGGTTATAGGCACGGGTTATCATTGAGG | |
| oIN428F | TATTCTGACCCATAATTCTGCAGTGTACACCGCTCCCCGCAAT | |
| oIN429R | TGCTACCAACCAGACAACAAGGGTTATTGTCGCGGTTGCCCCATA | |
| oIN430F | TATTCTGACCCATAATTCTGCAGTGTATGTGTCACCGCAAGGAAGG | |
| oIN431R | TGCTACCAACCAGACAACAAGGGTTGTCCTTGTTTACGAAAGAATCGCT | |
| oIN432F | CATCCGCAGTTTGAGAAAGGTAGCGGCAGCAAAGATAACATTGCTACAGTGGCAAATAGTTTTAACTGTGAAGCG | 75 |
| oIN433R | CATGGTATCCAGGCTCAGACAGCGGTTCAGGGCTCGCCGGGCAAAGGCCTGATTG | 55 |
| oIN436F | CATCCGCAGTTTGAGAAAGGTAGCGGCAGCACACGTAACCATACATCCTCATTAGAGAATCATTTG | 66 |
| oIN437R | CATGGTATCCAGGCTCAGACAGCGGTTCAGAGCATGGGTTGCTACGGCTGATTGG | 55 |
| oIN438F | CATCCGCAGTTTGAGAAAGGTAGCGGCAGCTCGTGCAATCGTATCAATAACGAAGCCC | 58 |
| oIN439R | CATGGTATCCAGGCTCAGACAGCGGTTCAGACTGCGGGTTATCACTGTCGAAACATCAGG | 60 |

| | | |
|---|---|---|
| oIN442F | AATGGTATTCTGACCCATAATTCTGCAGTGTATGAAGCACCGCAAGGA GAGATGGAAATAACG | 63 |
| oIN443R | TTCGCTGCTACCAACCAGACAACAAGGGTTCACCTGATGGGCAAACGC TTTTTCGTCC | 58 |
| oIN444F | AATGGTATTCTGACCCATAATTCTGCAGTGTATGAAGCGCCACAAGGA GAAATGGAAATTGC | 62 |
| oIN445R | TTCGCTGCTACCAACCAGACAACAAGGGTTAATCTGGCGGGCAAAGG CTTTCTCGTCTGG | 60 |
| oIN460F | CATCCGCAGTTTGAGAAAGGTAGCGGCAGCAGATCATTTGAGGATTCA CTGAATTCGGATTTATATTATCTTCACC | 76 |
| oIN461R | CATGGTATCCAGGCTCAGACAGCGGTTCAGACTTTCTTTATTGCTGAA GACCGGTTCAGGC | 61 |
| oIN462F | CATCCGCAGTTTGAGAAAGGTAGCGGCAGCCACCGTATTAACAACGAA GCTCCCTCC | 57 |
| oIN463R | CATGGTATCCAGGCTCAGACAGCGGTTCAGGCGGTGCATCACGACGG CGGATGC | 54 |
| oIN472F | AATGGTATTCTGACCCATAATTCTGCAGTGTATATTGCCCCTCGCAATG AGCTGGAAACTCAAC | 64 |
| oIN473R | TTCGCTGCTACCAACCAGACAACAAGGGTTGCTGTCTTTGTTTCCCCAT ATCGGCGC | 57 |
| oIN484F | AATGATATTCTGGTGCATAATAGCGAAATTTACGAAGCGCCGAAAGGA GAACGGG | 55 |
| oIN485R | CATTTCGGTTTCCCCCTGCGGAGCCGCATAAGCCTGACGGACAAAGG CCTCTTCACC | 57 |
| oIN486F | TATGCGGCTCCGCAGGGGGAAACCGAAATG | 30 |
| oIN487R | AATTTCGCTATTATGCACCAGAATATCATTG | 31 |
| oIN524F | AATGGTATTCTGACCCATAATTCTGCAGTGTATGTCGCGCCACAGGGA GACATGG | 55 |
| oIN525R | TTCGCTGCTACCAACCAGACAACAAGGGTTAATCTGGCGGGCAAAAGC ATCCTCC | 55 |
| oIN526F | AATGATATTCTGGTGCATAATAGCGAAATTTATGCTGCACCGCAAGGA GAAACCGAAACC | 60 |
| oIN527R | ATGATGATGATGATGATGGCTGCCGCTACCCAGCGCCTCCGCTTCACA ATTCATTGC | 57 |
| oIN530F | AACGAGAAGGAGGAATTAAAATCGAAAAAGGCTG | 34 |
| oIN531R | CATGGAATTCCTCCTGTTAGCCCAAAAAAACG | 32 |
| oIN538F | AATGAGAACGAAACCCTGAAGAAAAAGAACCTGC | 34 |
| oIN539R | TGAGATAGCTGCAGTCAGCTCGTTATCAAGG | 31 |
| oIN542F | TGACAATTAATCATCGGCTCGTATAATGTGTGG | 33 |
| oIN543R | CTGTTCGTGAGACGCAACTTCGTTTTCG | 28 |
| oIN597F | TTGGGCTAACAGGAGGAATTCCATGTGGAGCCATCCGCAGTTTGAGAA AGGTAG | 54 |
| oIN598R | TCGATTTTAATTCCTCCTTCTCGTTAATCTGGCGGGCGAAAGCCTCTTC C | 50 |
| oIN599F | TTGGGCTAACAGGAGGAATTCCATGTGGAGCCATCCGCAGTTTGAGAA AGGTAG | 54 |
| oIN600R | TCGATTTTAATTCCTCCTTCTCGTTAATTTGGCGAGCAAAAGCATCCTC TCCCGGT | 56 |
| oIN601F | TTGGGCTAACAGGAGGAATTCCATGTGGAGCCATCCGCAGTTTGAGAA AGGTAGC | 55 |
| oIN602R | TCGATTTTAATTCCTCCTTCTCGTTGGCTCGCCGGGCAAAGGCCTGA | 47 |
| oIN603F | TTGGGCTAACAGGAGGAATTCCATGTGGAGCCATCCGCAGTTTGAGAA AGGTAG | 54 |
| oIN604R | TCGATTTTAATTCCTCCTTCTCGTTAGCATGGGTTGCTACGGCTGATTG GTCA | 53 |

| | | |
|---|---|---|
| oIN605F | TAACGAGCTGACTGCAGCTATCTCAGAAGCGCCCATTGGCAAATTGGAAATAACACTG | 58 |
| oIN606R | TTTTCTTCAGGGTTTCGTTCTCATTATAGCCACGTGTAACAACCGCTGACTGG | 53 |
| oIN607F | TAACGAGCTGACTGCAGCTATCTCACAAGCGCCGGAAAGCCCAATGGAGTG | 51 |
| oIN608R | TTTTCTTCAGGGTTTCGTTCTCATTATAGGCACGGGTTATCATTGAGGATGAATCAGG | 58 |
| oIN610R | TATACGAGCCGATGATTAATTGTCAATGATGATGATGATGATGGCTGCCGCTACCC | 56 |
| oIN611F | AAACGAAGTTGCGTCTCACGAACAGTACGAAGCGCCGAAAGGAGAACGGGAAAC | 54 |
| oIN612R | TATACGAGCCGATGATTAATTGTCAATGATGATGATGATGATGGCTGCCGCTACCC | 56 |
| oIN613F | AAACGAAGTTGCGTCTCACGAACAGTATGTTGCGCCACAAGGAGAAATGGAAATCGCAA | 59 |
| oIN614R | TATACGAGCCGATGATTAATTGTCAATGATGATGATGATGATGGCTGCCGCTACC | 55 |
| oIN615F | AAACGAAGTTGCGTCTCACGAACAGTATGTCGCGCCACAGGGAGACATGGAA | 52 |
| oIN616R | TATACGAGCCGATGATTAATTGTCAATGATGATGATGATGATGGCTGCCGCTACCCA | 57 |
| oIN620F | AAACGAAGTTGCGTCTCACGAACAGTATGCGGCTCCGCAGGGGGAAACCGAAATAGCATTAGCGGCTATCTGGTGCG | 77 |

**Table S6. | Synthetic DNA fragments used in this work.**

| Fragment Name | Sequence 5' → 3' | Length (bp) |
|---|---|---|
| fIN001 - insert gp41-1_N | TTTTTTTGGGCTAACAGGAGGAATTCCATGTGGAGCCATCCGCAGTTTGAGAAAGGTAGCGGCAGCCGAGACCGTAGCCTTAGGCCAATTACCAACTTGGTGGCTACGGTCTCTAGCGGCTATTGTCTGGATCTGAAAACCCAGGTTCAGACACCGCAGGGTATGAAAGAAATTTCAAATATTCAGGTGGGTGATCTGGTTCTGAGCAATACCGGTTATAATGAAGTGCTGAATGTGTTTCCGAAAAGCAAAAAGAAAAGCTATAAAATCACCCTGGAAGATGGCAAAGAAATCATTTGTAGCGAAGAACACCTGTTTCCGACCCAGACCGGTGAAATGAATATTAGCGGTGGTCTGAAAGAAGGTATGTGCCTGTATGTTAAAGAATGACAATTAATCATCGGCTCGTATAATGTG | 417 |
| fIN002 - insert gp41-8_N | TTTTTTTGGGCTAACAGGAGGAATTCCATGTGGAGCCATCCGCAGTTTGAGAAAGGTAGCGGCAGCCGAGACCGTAGCCTTAGGCCAATTACCAACTTGGTGGCTACGGTCTCTCTGAACCGCTGTCTGAGCCTGGATACCATGGTTGTTACCAATGGTAAAGCCATTGAAATTCGTGATGTGAAAGTTGGTGATTGGCTGGAAAGCGAATGTGGTCCGGTTCAGGTTACCGAAGTTCTGCCGATTATCAAACAGCCGGTTTTTGAAATTGTGCTGAAAAGCGGCAAAAAAATCCGTGTTAGCGCCAATCATAAATTCCCGACCAAAGATGGTCTGAAAACCATTAATAGCGGTCTGAAAGTGGGCGATTTTCTGCGTAGCCGTGCAAAATGACAATTAATCATCGGCTCGTATAATGTG | 420 |
| fIN003 - insert NrdH-1_N | TTTTTTTGGGCTAACAGGAGGAATTCCATGTGGAGCCATCCGCAGTTTGAGAAAGGTAGCGGCAGCCGAGACCGTAGCCTTAGGCCAATTACCAACTTGGTGGCTACGGTCTCTAACCCTTGTTGTCTGGTTGGTAGCAGCGAAATCATTACCCGTAATTATGGTAAAACCACCATCAAAGAAGTGGTCGAGATCTTCGATAACGACAAAAACATTCAGGTGCTGGCCTTTAATACCCATACCGATAATATTGAATGGGCACCGATTAAAGCAGCACAGCTGACCCGTCCGAATGCAGAACTGGTTGAACTGGAAATTGATACCCTGCATGGTGTTAAAACCATTCGTTGTACACCGGATCATCCTGTGTATACCAAAAATCGTGG | 468 |

| | | |
|---|---|---|
| | TTATGTTCGTGCAGATGAACTGACCGATGATGATGAACTGGTGGTTGCAATTTGACAATTAATCATCGGCTCGTATAATGTG | |
| fIN004 - insert IMPDH-1_N | TTTTTTTGGGCTAACAGGAGGAATTCCATGTGGAGCCATCCGCAGTTTGAGAAAGGTAGCGGCAGCCGAGACCGTAGCCTTAGGCCAATTACCAACTTGGTGGCTACGGTCTCTGGTGGAGGATGTTTTGTTCCGGGTACACTGGTGAATACCGAAAATGGTCTGAAAAAAATCGAAGAAATCAAAGTGGGCGACAAAGTGTTTAGCCATACCGGTAAACTGCAAGAAGTTGTTGATACCCTGATCTTTGATCGTGATGAAGAGATTATTAGCATCAACGGTATCGACTGCACCAAAAACCATGAGTTTTATGTGATCGACAAAGAAAATGCCAATCGCGTGAACGAAGATAACATTCACCTGTTTGCACGTTGGGTTCATGCCGAAGAACTGGATATGAAAAAACATCTGCTGATCGAGCTGGAATGACAATTAATCATCGGCTCGTATAATGTG | 456 |
| fIN005 - insert SspGyrB_N | TTTTTTTGGGCTAACAGGAGGAATTCCATGTGGAGCCATCCGCAGTTTGAGAAAGGTAGCGGCAGCCGAGACCGTAGCCTTAGGCCAATTACCAACTTGGTGGCTACGGTCTCTGCTGGGGGTTGTTTTTCTGGAGATACATTAGTCGCTTTAACTGATGGTCGTAGCGTTAGCTTTGAGCAATTGGTTGAAGAAGAAAAACAAGGAAAACAAAACTTTTGTTATACCATCCGCCATGATGGTTCTATAGGGGTTGAAAAAATCATCAATGCCCGCAAAACAAAAACTAATGCGAAGGTAATCAAGGTTACGTTGGACAATGGTGAGTCTATTATTTGCACCCCGGATCATAAATTCATGTTGCGGGATGGGAGCTACAAATGTGCGATGGATTTAACTCTCGATGATTCGTTAATGCCGTTACACCGAAAAATTTCGACTACGGAAGATTCTGGTCATGCGTGACAATTAATCATCGGCTCGTATAATGTG | 492 |
| fIN006 - insert gp41-1_C | TTTTTTTGGGCTAACAGGAGGAATTCCATGATGCTGAAAAAAATCCTGAAAATCGAGGAACTGGATGAACGCGAACTGATTGATATTGAAGTTAGCGGTAACCACCTGTTCTATGCCAATGATATTCTGACCCATAATTCAAGCTCGCGAGACCGTAGCCTTAGGCCAATTACACTGAAATTTATCTGCACTACAGGTAAACTGCCGGTGCCCTGGCCAACACCAACTTGGTGGCTACGGTCTCTGGTAGCGGCAGCCATCATCATCATCATCATTGACAATTAATCATCGGCTCGTATAATGTG | 305 |
| fIN007 - insert gp41-8_C | TTTTTTTGGGCTAACAGGAGGAATTCCATGTGTGAAATCTTTGAAAACGAGATCGACTGGGATGAAATTGCCAGCATTGAATATGTTGGTGTGGAAGAAACCATCGATATTAACGTGACCAATGATCGTCTGTTTTTTGCCAATGGTATTCTGACCCATAATTCTGCAGTGCGAGACCGTAGCCTTAGGCCAATTACACTGAAATTTATCTGCACTACAGGTAAACTGCCGGTGCCCTGGCCAACACCAACTTGGTGGCTACGGTCTCTGGTAGCGGCAGCCATCATCATCATCATCATTGACAATTAATCATCGGCTCGTATAATGTG | 329 |
| fIN008 - insert NrdJ-1_C | TTTTTTTGGGCTAACAGGAGGAATTCCATGGAAGCCAAAACCTATATCGGCAAACTGAAAAGCCGTAAAATTGTGAGCAACGAGGATACCTATGATATTCAGACCAGCACCCATAACTTTTTCGCCAATGATATTCTGGTGCATAATAGCGAAATTCGAGACCGTAGCCTTAGGCCAATTACACTGAAATTTATCTGCACTACAGGTAAACTGCCGGTGCCCTGGCCAACACCAACTTGGTGGCTACGGTCTCTGGTAGCGGCAGCCATCATCATCATCATCATTGACAATTAATCATCGGCTCGTATAATGTG | 314 |
| fIN009 - insert IMPDH-1_C | TTTTTTTGGGCTAACAGGAGGAATTCCATGAAATTCAAACTGAAAGAGATCACCAGCATCGAAACCAAACACTATAAAGGCAAAGTTCATGATCTGACCGTGAATCAGGATCATAGCTATAACGTTCGTGGCACCGTTGTTCATAATAGCATCTGCCGAGACCGTAGCCTTAGGCCAATTACACTGAAATTTATCTGCACTACAGGTAAACTGCCGGTGCCCTGGCCAACACCAACTTGGTGGCTACGGTCTCTGGTAGCGGCAGCCATCATCATCATCATCATTGACAATTAATCATCGGCTCGTATAATGTG | 314 |
| fIN010 - insert SspGyrB_C | TTTTTTTGGGCTAACAGGAGGAATTCCATGGAAGCAGTATTAAATTACAATCACAGAATTGTAAATATTGAAGCTGTGTCAGAAACAATCGATGTTTATGATATTGAGGTTCCCCACACCCACAATTTTGCTTTGGCAAGCGGAGTGTTTGTCCATAACTCAGCGAAGCGAGACCGTAGCCTTAGGCCAATTACACTGAAATTTATCTGCACTACAGGTAAACTGCCGGTGCCCTGGCCAAC | 326 |

| | |
|---|---|
| | ACCAACTTGGTGGCTACGGTCTCTGGTAGCGGCAGCCATCATCATCAT CATCATTGACAATTAATCATCGGCTCGTATAATGTG |

**Table S7. | Library peptides detected in this work.** Colored squares indicate whether for this NRPS an expected cyclic NRP (yellow), linear NRP (green), both (blue), or no expected NRP (red) was found. Peptides that were confirmed by synthesis are marked in grey. Lowercase italic letters denote D-amino acids; for threonine and leucine, these correspond to D-*allo*-threonine and D-*allo*-isoleucine, respectively.

| | NRPS | Pep. | sequence | Sum formula | Exact mass | Δppm |
|---|---|---|---|---|---|---|
| | **AHQ** | (15) | cyclo[*v*-L-*v*-V] | C21H38N4O4 | 410.2893 | 0.7 |
| | | (16) | *v*-L-*v*-V | C21H40N4O5 | 428.2999 | 0.8 |
| | **AHR** | (17) | *v*-L-*v*-L | C22H42N4O5 | 442.3155 | 0.7 |
| | **AHS** | (18) | cyclo[*v*-L-*v*-K] | C22H41N5O4 | 439.3159 | -0.1 |
| | **AHT** | (19) | cyclo[*v*-L-V-*v*-V] | C26H47N5O5 | 509.3577 | -0.6 |
| | | (20) | *v*-L-V-*v*-V | C26H49N5O6 | 527.3683 | 0.1 |
| | **AHU** | (21) | *v*-L-*v*-Y-F | C34H49N5O7 | 639.3632 | 1.7 |
| | | (22) | *v*-L-*v*-Y-W | C36H50N6O7 | 678.3741 | 2.0 |
| | **AHV** | (23) | cyclo[*v*-L-V-*f*-*l*-L] | C37H60N6O6 | 684.4574 | -0.4 |
| | | (24) | *v*-L-V-*f*-*l*-L | C37H62N6O7 | 702.4680 | 0.1 |
| | **AIQ** | (25) | cyclo[*v*-L-*s*-V] | C19H34N4O5 | 398.2529 | 0.3 |
| | | (26) | *v*-L-*s*-V | C19H36N4O6 | 416.2635 | 0.6 |
| | **AIR** | (27) | *v*-L-*s*-L | C20H38N4O6 | 430.2791 | -1.8 |
| | **AIS** | **(1)** | cyclo[*v*-L-*s*-K] | C20H37N5O5 | 427.2795 | -0.8 |
| | **AIT** | (28) | cyclo[*v*-L-S-*v*-V] | C24H43N5O6 | 497.3213 | 0.4 |
| | **AIU** | (29) | *v*-L-*s*-Y-F | C32H45N5O8 | 627.3268 | 0.9 |
| | | (30) | *v*-L-*s*-Y-W | C34H46N6O8 | 666.3377 | 0.1 |
| | **AIV** | (31) | *v*-L-S-*f*-*l*-L | C35H58N6O8 | 690.4316 | 0.3 |
| | **AJQ** | (32) | cyclo[*v*-*l*-*y*-V] | C25H38N4O5 | 474.2842 | 0.6 |
| | | (33) | *v*-*l*-*y*-V | C25H40N4O6 | 492.2948 | 0.1 |
| | **AJR** | | | | | |
| | **AJS** | | | | | |
| | **AJT** | (34) | cyclo[*v*-*l*-Y-*v*-V] | C30H47N5O6 | 573.3526 | 0.5 |
| | **AJU** | (35) | *v*-*l*-*y*-Y-F | C38H49N5O8 | 703.3581 | 1.0 |
| | | (36) | *v*-*l*-*y*-Y-W | C40H50N6O8 | 742.3690 | 1.2 |
| | **AJV** | | | | | |
| | **AKQ** | (37) | cyclo[*v*-L-*f*-V] | C25H38N4O4 | 458.2893 | -0.2 |
| | | (38) | *v*-L-*f*-V | C25H40N4O5 | 476.2999 | 0.5 |
| | **AKR** | (39) | *v*-L-*f*-L | C26H42N4O5 | 490.3155 | 0.8 |
| | **AKS** | (40) | cyclo[*v*-L-*f*-K] | C26H41N5O4 | 487.3159 | 0.3 |
| | **AKT** | (41) | cyclo[*v*-L-F-*v*-V] | C30H47N5O5 | 557.3577 | -0.4 |
| | **AKU** | (42) | *v*-L-*f*-Y-F | C38H49N5O7 | 687.3632 | 1.6 |
| | | (43) | *v*-L-*f*-Y-W | C40H50N6O7 | 726.3741 | 1.2 |
| | **AKV** | **(3)** | cyclo[*v*-L-F-*f*-*l*-L] | C41H60N6O6 | 732.4574 | 1.0 |
| | | (44) | *v*-L-F-*f*-*l*-L | C41H62N6O7 | 750.4680 | -0.2 |
| | **ALQ** | **(2)** | cyclo[*v*-L-*f*-*l*-V] | C31H49N5O5 | 571.3734 | 1.1 |
| | | (45) | *v*-L-*f*-*l*-V | C31H51N5O6 | 589.3839 | 1.9 |
| | **ALR** | (46) | cyclo[*v*-L-*f*-*l*-L] | C32H51N5O5 | 585.3890 | -1.2 |
| | | (47) | *v*-L-*f*-*l*-L | C32H53N5O6 | 603.3996 | 0.6 |
| | **ALS** | (48) | cyclo[*v*-L-*f*-*l*-K] | C32H52N6O5 | 600.3999 | 0.5 |
| | **ALT** | (49) | *v*-L-*f*-L-*v*-V | C36H60N6O7 | 688.4523 | 1.3 |
| | **ALU** | (50) | *v*-L-*f*-*l*-Y-F | C44H60N6O8 | 800.4473 | -1.6 |

| | | | | | | |
|---|---|---|---|---|---|---|
| | | (51) | *v*-L-*f*-*l*-Y-W | C46H61N7O8 | 839.4582 | -1.4 |
| | **ALV** | (52) | *v*-L-*f*-L-*f*-*l*-L | C47H73N7O8 | 863.5521 | -0.9 |
| | **AMQ** | (53) | cyclo[*v*-L-*f*-*l*-V] | C31H49N5O5 | 571.3734 | -1.1 |
| | | (54) | cyclo[*v*-L-*w*-*l*-V] | C33H50N6O5 | 610.3843 | -0.7 |
| | **AMR** | (55) | cyclo[*v*-L-*f*-*l*-L] | C32H51N5O5 | 585.3890 | -0.3 |
| | | (56) | cyclo[*v*-L-*w*-*l*-L] | C34H52N6O5 | 624.3999 | 0.2 |
| | **AMS** | (57) | cyclo[*v*-L-*w*-*l*-K] | C34H53N7O5 | 639.4108 | 1.7 |
| | **AMT** | (58) | *v*-L-*f*-L-*v*-V | C36H60N6O7 | 688.4523 | -0.4 |
| | | (59) | *v*-L-*w*-L-*v*-V | C38H61N7O7 | 727.4632 | -0.8 |
| | **AMU** | | | | | |
| | **AMV** | (60) | *v*-L-*f*-L-*f*-*l*-L | C47H73N7O8 | 863.5521 | -2.0 |
| | | (61) | *v*-L-*w*-L-*f*-*l*-L | C49H74N8O8 | 902.5630 | -1.3 |
| | **ANQ** | (62) | cyclo[*v*-L-*w*-*l*-V] | C33H50N6O5 | 610.3843 | -0.4 |
| | | (63) | cyclo[*v*-L-*w*-*f*-V] | C36H48N6O5 | 644.3686 | 0.0 |
| | **ANR** | (64) | cyclo[*v*-L-*w*-*l*-L] | C34H52N6O5 | 624.3999 | 1.1 |
| | | (65) | cyclo[*v*-L-*w*-*f*-L] | C37H50N6O5 | 658.3843 | 1.3 |
| | **ANS** | (66) | cyclo[*v*-L-*w*-*l*-K] | C34H53N7O5 | 639.4108 | 0.8 |
| | | (67) | cyclo[*v*-L-*w*-*f*-K] | C37H51N7O5 | 673.3952 | 0.7 |
| | **ANT** | (68) | *v*-L-*w*-L-*v*-V | C38H61N7O7 | 727.4632 | 0.2 |
| | | (69) | *v*-L-*w*-F-*v*-V | C41H59N7O7 | 761.4476 | 0.2 |
| | **ANU** | | | | | |
| | **ANV** | (70) | *v*-L-*w*-L-*f*-*l*-L | C49H74N8O8 | 902.5630 | -0.7 |
| | | (71) | *v*-L-*w*-F-*f*-*l*-L | C52H72N8O8 | 936.5473 | -1.7 |
| | **AOQ** | (72) | cyclo[*v*-L-*l*-*i*-V] | C28H51N5O5 | 537.3890 | -0.6 |
| | | (73) | *v*-L-*l*-*i*-V | C28H53N5O6 | 555.3996 | 1.4 |
| | **AOR** | (74) | cyclo[*v*-L-*l*-*i*-L] | C29H53N5O5 | 551.4047 | 0.1 |
| | | (75) | *v*-L-*l*-*i*-L | C29H55N5O6 | 569.4152 | 0.7 |
| | **AOS** | | | | | |
| | **AOT** | (76) | *v*-L-*l*-I-*v*-V | C33H62N6O7 | 654.4680 | 0.1 |
| | **AOU** | (77) | *v*-L-*l*-*i*-Y-F | C41H62N6O8 | 766.4629 | -0.7 |
| | | (78) | *v*-L-*l*-*i*-Y-W | C43H63N7O8 | 805.4738 | -1.0 |
| | **AOV** | | | | | |
| | **APQ** | (**4**) | cyclo[*v*-L-bA-*y*-V] | C28H43N5O6 | 545.3213 | 1.3 |
| | **APR** | | | | | |
| | **APS** | | | | | |
| | **APT** | (79) | *v*-L-bA-Y-*v*-V | C33H54N6O8 | 662.4003 | -0.5 |
| | **APU** | (80) | *v*-L-bA-*y*-Y-F | C41H54N6O9 | 774.3952 | -0.1 |
| | | (81) | *v*-L-bA-*y*-Y-W | C43H55N7O9 | 813.4061 | -0.1 |
| | **APV** | (82) | *v*-L-bA-Y-*f*-*l*-L | C44H67N7O9 | 837.5000 | -0.1 |
| | **BHQ** | (83) | cyclo[*r*-L-*v*-V] | C22H41N7O4 | 467.3220 | -3.9 |
| | | (84) | *r*-L-*v*-V | C22H43N7O5 | 485.3326 | -0.3 |
| | **BHR** | (85) | cyclo[*r*-L-*v*-L] | C23H43N7O4 | 481.3377 | -0.1 |
| | | (86) | *r*-L-*v*-L | C23H45N7O5 | 499.3482 | -0.4 |
| | **BHS** | | | | | |
| | **BHT** | (87) | cyclo[*r*-L-V-*v*-V] | C27H50N8O5 | 566.3904 | 0.0 |
| | | (88) | *r*-L-V-*v*-V | C27H52N8O6 | 584.4010 | -0.1 |
| | **BHU** | | | | | |
| | **BHV** | (89) | cyclo[*r*-L-V-*f*-*l*-L] | C38H63N9O6 | 741.4901 | -0.4 |
| | **BIQ** | (90) | cyclo[*r*-L-*s*-V] | C20H37N7O5 | 455.2856 | 0.2 |
| | **BIR** | (91) | cyclo[*r*-L-*s*-L] | C21H39N7O5 | 469.3013 | 1.0 |
| | | (92) | *r*-L-*s*-L | C21H41N7O6 | 487.3118 | 1.0 |
| | **BIS** | | | | | |
| | **BIT** | (93) | cyclo[*r*-L-S-*v*-V] | C25H46N8O6 | 554.3540 | 2.0 |
| | **BIU** | | | | | |
| | **BIV** | (**7**) | cyclo[*r*-L-S-*f*-*l*-L] | C36H59N9O7 | 729.4537 | 0.0 |

| | | | | | | |
|---|---|---|---|---|---|---|
| | | (94) | *r*-L-S-*f*-*l*-L | C36H61N9O8 | 747.4643 | 0.3 |
| | **BJQ** | | | | | |
| | **BJR** | | | | | |
| | **BJS** | | | | | |
| | **BJT** | (95) | cyclo[*r*-*l*-Y-*v*-V] | C31H50N8O6 | 630.3853 | -0.1 |
| | | (96) | *r*-*l*-Y-*v*-V | C31H52N8O7 | 648.3959 | -0.5 |
| | **BJU** | (97) | *r*-*l*-*y*-Y-F | C39H52N8O8 | 760.3908 | -0.1 |
| | | (98) | *r*-*l*-*y*-Y-W | C41H53N9O8 | 799.4017 | 0.6 |
| | **BJV** | | | | | |
| | **BKQ** | **(5)** | cyclo[*r*-L-*f*-V] | C26H41N7O4 | 515.3220 | -1.2 |
| | | (99) | *r*-L-*f*-V | C26H43N7O5 | 533.3326 | -0.3 |
| | **BKR** | (100) | *r*-L-*f*-L | C27H45N7O5 | 547.3482 | 0.4 |
| | **BKS** | | | | | |
| | **BKT** | (101) | cyclo[*r*-L-F-*v*-V] | C31H50N8O5 | 614.3904 | 1.5 |
| | **BKU** | (102) | *r*-L-*f*-Y-F | C39H52N8O7 | 744.3959 | -0.6 |
| | | (103) | *r*-L-*f*-Y-W | C41H53N9O7 | 783.4068 | -0.9 |
| | **BKV** | | | | | |
| | **BLQ** | (104) | cyclo[*r*-L-*f*-*l*-V] | C32H52N8O5 | 628.4061 | -0.4 |
| | **BLR** | (105) | cyclo[*r*-L-*f*-*l*-L] | C33H54N8O5 | 642.4217 | -0.2 |
| | **BLS** | | | | | |
| | **BLT** | (106) | *r*-L-*f*-L-*v*-V | C37H63N9O7 | 745.4850 | -0.4 |
| | **BLU** | (107) | *r*-L-*f*-*l*-Y-F | C45H63N9O8 | 857.4800 | -1.0 |
| | | (108) | *r*-L-*f*-*l*-Y-W | C47H64N10O8 | 896.4909 | -1.2 |
| | **BLV** | (109) | *r*-L-*f*-L-*f*-*l*-L | C48H76N10O8 | 920.5848 | -1.0 |
| | **BMQ** | (110) | cyclo[*r*-L-*f*-*l*-V] | C32H52N8O5 | 628.4061 | -0.1 |
| | | (111) | cyclo[*r*-L-*w*-*l*-V] | C34H53N9O5 | 667.4170 | -2.0 |
| | **BMR** | (112) | cyclo[*r*-L-*f*-*l*-L] | C33H54N8O5 | 642.4217 | -1.2 |
| | | (113) | cyclo[*r*-L-*w*-*l*-L] | C35H55N9O5 | 681.4326 | -2.4 |
| | **BMS** | | | | | |
| | **BMT** | (114) | *r*-L-*w*-L-*v*-V | C39H64N10O7 | 784.4959 | 0.7 |
| | **BMU** | | | | | |
| | **BMV** | | | | | |
| | **BNQ** | (115) | cyclo[*r*-L-*w*-*l*-V] | C34H53N9O5 | 667.4170 | -0.8 |
| | | (116) | cyclo[*r*-L-*w*-*f*-V] | C37H51N9O5 | 701.4013 | -0.4 |
| | **BNR** | **(6)** | cyclo[*r*-L-*w*-*l*-L] | C35H55N9O5 | 681.4326 | -1.6 |
| | | (117) | cyclo[*r*-L-*w*-*f*-L] | C38H53N9O5 | 715.4170 | -2.9 |
| | **BNS** | | | | | |
| | **BNT** | (118) | *r*-L-*w*-L-*v*-V | C39H64N10O7 | 784.4959 | 0.3 |
| | | (119) | *r*-L-*w*-F-*v*-V | C42H62N10O7 | 818.4803 | 0.2 |
| | **BNU** | | | | | |
| | **BNV** | (120) | *r*-L-*w*-L-*f*-*l*-L | C50H77N11O8 | 959.5957 | -0.9 |
| | **BOQ** | (121) | cyclo[*r*-L-*l*-*i*-V] | C29H54N8O5 | 594.4217 | 0.2 |
| | **BOR** | (122) | cyclo[*r*-L-*l*-*i*-L] | C30H56N8O5 | 608.4374 | -1.9 |
| | | (123) | *r*-L-*l*-*i*-L | C30H58N8O6 | 626.4479 | 1.1 |
| | **BOS** | | | | | |
| | **BOT** | (124) | *r*-L-*l*-I-*v*-V | C34H65N9O7 | 711.5007 | 0.8 |
| | **BOU** | (125) | *r*-L-*l*-*i*-Y-F | C42H65N9O8 | 823.4956 | -0.3 |
| | | (126) | *r*-L-*l*-*i*-Y-W | C44H66N10O8 | 862.5065 | 0.0 |
| | **BOV** | | | | | |
| | **BPQ** | | | | | |
| | **BPR** | | | | | |
| | **BPS** | | | | | |
| | **BPT** | | | | | |
| | **BPU** | **(8)** | *r*-L-bA-*y*-Y-F | C42H57N9O9 | 831.4279 | 0.0 |
| | | (127) | *r*-L-bA-*y*-Y-W | C44H58N10O9 | 870.4388 | 0.1 |

| | | | | | |
|---|---|---|---|---|---|
| **BPV** | (128) | *r*-L-bA-Y-*f*-*l*-L | C45H70N10O9 | 894.5327 | -0.6 |
| **CHQ** | | | | | |
| **CHR** | (**9**) | C2-I-T-*v*-L | C23H42N4O7 | 486.3054 | 2.5 |
| **CHS** | | | | | |
| **CHT** | | | | | |
| **CHU** | | | | | |
| **CHV** | (129) | C2-I-T-V-*f*-*l*-L | C38H62N6O9 | 746.4578 | 2.7 |
| **CIQ** | | | | | |
| **CIR** | | | | | |
| **CIS** | | | | | |
| **CIT** | | | | | |
| **CIU** | | | | | |
| **CIV** | (**10**) | C2-I-T-S-*f*-*l*-L | C36H58N6O10 | 734.4214 | 2.2 |
| **CJQ** | | | | | |
| **CJR** | | | | | |
| **CJS** | | | | | |
| **CJT** | | | | | |
| **CJU** | | | | | |
| **CJV** | | | | | |
| **CKQ** | | | | | |
| **CKR** | | | | | |
| **CKS** | | | | | |
| **CKT** | (130) | C2-I-T-F-*v*-V | C31H49N5O8 | 619.3581 | 1.9 |
| **CKU** | | | | | |
| **CKV** | | | | | |
| **CLQ** | (131) | C2-I-T-*f*-*l*-V | C32H51N5O8 | 633.3738 | 2.9 |
| **CLR** | (132) | C2-I-T-*f*-*l*-L | C33H53N5O8 | 647.3894 | 2.9 |
| **CLS** | (133) | C2-I-T-*f*-*l*-K | C33H54N6O8 | 662.4003 | 3.6 |
| **CLT** | (134) | C2-I-T-*f*-L-*v*-V | C37H60N6O9 | 732.4422 | 2.4 |
| **CLU** | | | | | |
| **CLV** | | | | | |
| **CMQ** | (135) | C2-I-T-*f*-*l*-V | C32H51N5O8 | 633.3738 | 2.1 |
| | (136) | C2-I-T-*w*-*l*-V | C34H52N6O8 | 672.3847 | 2.3 |
| **CMR** | (137) | C2-I-T-*w*-*l*-L | C35H54N6O8 | 686.4003 | 2.2 |
| **CMS** | (138) | C2-I-T-*f*-*l*-K | C33H54N6O8 | 662.4003 | 1.2 |
| | (139) | C2-I-T-*w*-*l*-K | C35H55N7O8 | 701.4112 | 2.8 |
| **CMT** | | | | | |
| **CMU** | | | | | |
| **CMV** | | | | | |
| **CNQ** | (140) | C2-I-T-*w*-*l*-V | C34H52N6O8 | 672.3847 | 1.7 |
| | (141) | C2-I-T-*w*-*f*-V | C37H50N6O8 | 706.3690 | 2.1 |
| **CNR** | | | | | |
| **CNS** | | | | | |
| **CNT** | | | | | |
| **CNU** | | | | | |
| **CNV** | | | | | |
| **COQ** | (142) | C2-I-T-*l*-*i*-V | C29H53N5O8 | 599.3894 | 3.3 |
| **COR** | (143) | C2-I-T-*l*-*i*-L | C30H55N5O8 | 613.4051 | 3.2 |
| **COS** | | | | | |
| **COT** | | | | | |
| **COU** | (144) | C2-I-T-*l*-*i*-Y-W | C44H63N7O10 | 849.4636 | 3.6 |
| **COV** | | | | | |
| **CPQ** | | | | | |
| **CPR** | (145) | C2-I-T-bA-*y*-L | C30H47N5O9 | 621.3374 | 4.3 |
| **CPS** | | | | | |

| | | | | | | |
|---|---|---|---|---|---|---|
| | **CPT** | (146) | C2-I-T-bA-Y-*v*-V | C34H54N6O10 | 706.3901 | 2.4 |
| | **CPU** | (147) | C2-I-cyclo[T-bA-*y*-Y-F] | C42H52N6O10 | 800.3745 | 2.6 |
| | | (148) | C2-I-T-bA-*y*-Y-F | C42H54N6O11 | 818.3851 | 2.5 |
| | | (149) | C2-I-cyclo[T-bA-*y*-Y-W] | C44H53N7O10 | 839.3854 | 3.3 |
| | | (150) | C2-I-T-bA-*y*-Y-W | C44H55N7O11 | 857.3960 | 2.3 |
| | **CPV** | (151) | C2-I-T-bA-Y-*f*-*l*-L | C45H67N7O11 | 881.4899 | 2.6 |
| | **DHQ** | (152) | C4-P-G-*v*-V | C21H36N4O6 | 440.2635 | 2.6 |
| | **DHR** | (153) | C4-P-G-*v*-L | C22H38N4O6 | 454.2791 | 2.9 |
| | **DHS** | | | | | |
| | **DHT** | (154) | C4-P-G-V-*v*-V | C26H45N5O7 | 539.3319 | 2.6 |
| | **DHU** | (**11**) | C4-P-G-*v*-Y-F | C34H45N5O8 | 651.3268 | 2.0 |
| | | (155) | C4-P-G-*v*-Y-W | C36H46N6O8 | 690.3377 | 2.7 |
| | **DHV** | (156) | C4-P-G-V-*f*-*l*-L | C37H58N6O8 | 714.4316 | 2.1 |
| | **DIQ** | | | | | |
| | **DIR** | (157) | C4-P-G-*s*-L | C20H34N4O7 | 442.2427 | 2.5 |
| | **DIS** | | | | | |
| | **DIT** | (158) | C4-P-G-S-*v*-V | C24H41N5O8 | 527.2955 | 3.2 |
| | **DIU** | | | | | |
| | **DIV** | (159) | C4-P-G-S-*f*-*l*-L | C35H54N6O9 | 702.3952 | 1.9 |
| | **DJQ** | | | | | |
| | **DJR** | | | | | |
| | **DJS** | | | | | |
| | **DJT** | (160) | C4-P-G-Y-*v*-V | C30H45N5O8 | 603.3268 | 2.6 |
| | **DJU** | | | | | |
| | **DJV** | | | | | |
| | **DKQ** | | | | | |
| | **DKR** | | | | | |
| | **DKS** | | | | | |
| | **DKT** | | | | | |
| | **DKU** | | | | | |
| | **DKV** | | | | | |
| | **DLQ** | (161) | C4-P-G-*f*-*l*-V | C31H47N5O7 | 601.3475 | 0.7 |
| | **DLR** | (162) | C4-P-G-*f*-*l*-L | C32H49N5O7 | 615.3632 | -0.2 |
| | **DLS** | (163) | C4-P-G-*f*-*l*-K | C32H50N6O7 | 630.3741 | 2.0 |
| | **DLT** | (164) | C4-P-G-*f*-L-*v*-V | C36H56N6O8 | 700.4160 | 1.5 |
| | **DLU** | (165) | C4-P-G-*f*-*l*-Y-F | C44H56N6O9 | 812.4109 | 1.3 |
| | | (166) | C4-P-G-*f*-*l*-Y-W | C46H57N7O9 | 851.4218 | 0.8 |
| | **DLV** | | | | | |
| | **DMQ** | (167) | C4-P-G-*f*-*l*-V | C31H47N5O7 | 601.3475 | 1.5 |
| | | (168) | C4-P-G-*w*-*l*-V | C33H48N6O7 | 640.3584 | 2.2 |
| | **DMR** | (169) | C4-P-G-*w*-*l*-L | C34H50N6O7 | 654.3741 | 1.5 |
| | **DMS** | (170) | C4-P-G-*f*-*l*-K | C32H50N6O7 | 630.3741 | 2.7 |
| | | (171) | C4-P-G-*w*-*l*-K | C34H51N7O7 | 669.3850 | 2.7 |
| | **DMT** | | | | | |
| | **DMU** | | | | | |
| | **DMV** | (**12**) | C4-P-G-*f*-L-*f*-*l*-L | C47H69N7O9 | 875.5157 | 1.5 |
| | | (172) | C4-P-G-*w*-L-*f*-*l*-L | C49H70N8O9 | 914.5266 | 2.1 |
| | **DNQ** | (173) | C4-P-G-*w*-*l*-V | C33H48N6O7 | 640.3584 | 1.6 |
| | | (174) | C4-P-G-*w*-*f*-V | C36H46N6O7 | 674.3428 | 1.2 |
| | **DNR** | (175) | C4-P-G-*w*-*l*-L | C34H50N6O7 | 654.3741 | 1.6 |
| | **DNS** | | | | | |
| | **DNT** | (176) | C4-P-G-*w*-L-*v*-V | C38H57N7O8 | 739.4269 | 1.8 |
| | | (177) | C4-P-G-*w*-F-*v*-V | C41H55N7O8 | 773.4112 | 2.6 |
| | **DNU** | | | | | |
| | **DNV** | (178) | C4-P-G-*w*-L-*f*-*l*-L | C49H70N8O9 | 914.5266 | 2.0 |

| | | | | | |
|---|---|---|---|---|---|
| DOQ | (179) | C4-P-G-*l*-*i*-V | C28H49N5O7 | 567.3632 | 3.3 |
| DOR | (180) | C4-P-G-*l*-*i*-L | C29H51N5O7 | 581.3788 | 1.4 |
| DOS | | | | | |
| DOT | (181) | C4-P-G-*l*-I-*v*-V | C33H58N6O8 | 666.4316 | 2.7 |
| DOU | | | | | |
| DOV | | | | | |
| DPQ | | | | | |
| DPR | | | | | |
| DPS | | | | | |
| DPT | (182) | C4-P-G-bA-Y-*v*-V | C33H50N6O9 | 674.3639 | 1.6 |
| DPU | (183) | C4-P-G-bA-*y*-Y-F | C41H50N6O10 | 786.3588 | 1.0 |
| | (184) | C4-P-G-bA-*y*-Y-W | C43H51N7O10 | 825.3697 | 2.3 |
| DPV | (185) | C4-P-G-bA-Y-*f*-*l*-L | C44H63N7O10 | 849.4636 | 2.1 |
| EHQ | (186) | C6-T-A-*v*-V | C23H42N4O7 | 486.3054 | 1.3 |
| | (187) | C5-T-A-*v*-V | C22H40N4O7 | 472.2897 | 1.2 |
| EHR | (188) | C6-T-A-*v*-L | C24H44N4O7 | 500.3210 | 1.6 |
| | (189) | C5-T-A-*v*-L | C23H42N4O7 | 486.3054 | 1.3 |
| EHS | | | | | |
| EHT | (190) | C6-T-A-V-*v*-V | C28H51N5O8 | 585.3738 | 2.5 |
| | (191) | C5-T-A-V-*v*-V | C27H49N5O8 | 571.3581 | 1.6 |
| EHU | | | | | |
| EHV | (192) | C6-T-A-V-*f*-*l*-L | C39H64N6O9 | 760.4735 | -0.8 |
| | (193) | C5-T-A-V-*f*-*l*-L | C38H62N6O9 | 746.4578 | -0.4 |
| EIQ | | | | | |
| EIR | | | | | |
| EIS | | | | | |
| EIT | (194) | C6-T-A-S-*v*-V | C26H47N5O9 | 573.3374 | 1.1 |
| | (195) | C5-T-A-S-*v*-V | C25H45N5O9 | 559.3217 | 0.5 |
| EIU | | | | | |
| EIV | (196) | C6-T-A-S-*f*-*l*-L | C37H60N6O10 | 748.4371 | 0.0 |
| | (197) | C5-T-A-S-*f*-*l*-L | C36H58N6O10 | 734.4214 | -0.4 |
| EJQ | | | | | |
| EJR | | | | | |
| EJS | | | | | |
| EJT | (**13**) | C6-T-*a*-Y-*v*-V | C32H51N5O9 | 649.3687 | -0.4 |
| | (198) | C5-T-*a*-Y-*v*-V | C31H49N5O9 | 635.3530 | -0.5 |
| EJU | (199) | C6-T-*a*-*y*-Y-F | C40H51N5O10 | 761.3636 | 0.1 |
| | (200) | C5-T-*a*-*y*-Y-F | C39H49N5O10 | 747.3479 | 0.2 |
| EJV | | | | | |
| EKQ | | | | | |
| EKR | | | | | |
| EKS | (201) | C6-T-A-*f*-K | C28H45N5O7 | 563.3319 | 1.2 |
| | (202) | C5-T-A-*f*-K | C27H43N5O7 | 549.3162 | 1.7 |
| EKT | | | | | |
| EKU | | | | | |
| EKV | | | | | |
| ELQ | (203) | C6-T-A-*f*-*l*-V | C33H53N5O8 | 647.3894 | 1.7 |
| | (204) | C5-T-A-*f*-*l*-V | C32H51N5O8 | 633.3738 | 1.5 |
| ELR | (205) | C6-T-A-*f*-*l*-L | C34H55N5O8 | 661.4051 | 2.3 |
| | (206) | C5-T-A-*f*-*l*-L | C33H53N5O8 | 647.3894 | 1.2 |
| ELS | (207) | C6-T-A-*f*-*l*-K | C34H56N6O8 | 676.4160 | 2.4 |
| | (208) | C5-T-A-*f*-*l*-K | C33H54N6O8 | 662.4003 | 2.9 |
| ELT | | | | | |
| ELU | (209) | C5-T-A-*f*-*l*-Y-F | C45H60N6O10 | 844.4371 | 1.0 |
| ELV | (210) | C6-T-A-*f*-L-*f*-*l*-L | C49H75N7O10 | 921.5575 | 2.6 |

| | | | | | | |
|---|---|---|---|---|---|---|
| | | (211) | C5-T-A-*f*-L-*f*-*l*-L | C48H73N7O10 | 907.5419 | 2.7 |
| | **EMQ** | (212) | C6-T-A-*f*-*l*-V | C33H53N5O8 | 647.3894 | 0.8 |
| | | (213) | C5-T-A-*f*-*l*-V | C32H51N5O8 | 633.3738 | 0.9 |
| | **EMR** | (214) | C6-T-A-*w*-*l*-L | C36H56N6O8 | 700.4160 | 0.5 |
| | **EMS** | (215) | C6-T-A-*f*-*l*-K | C34H56N6O8 | 676.4160 | 1.2 |
| | | (216) | C5-T-A-*f*-*l*-K | C33H54N6O8 | 662.4003 | 1.0 |
| | | (217) | C6-T-A-*w*-*l*-K | C36H57N7O8 | 715.4269 | 1.9 |
| | **EMT** | | | | | |
| | **EMU** | | | | | |
| | **EMV** | | | | | |
| | **ENQ** | (218) | C6-T-A-*w*-*l*-V | C35H54N6O8 | 686.4003 | 0.7 |
| | | (219) | C5-T-A-*w*-*l*-V | C34H52N6O8 | 672.3847 | 1.3 |
| | **ENR** | (220) | C6-T-A-*w*-*l*-L | C36H56N6O8 | 700.4160 | 0.6 |
| | | (221) | C5-T-A-*w*-*l*-L | C35H54N6O8 | 686.4003 | 0.7 |
| | **ENS** | | | | | |
| | **ENT** | | | | | |
| | **ENU** | | | | | |
| | **ENV** | | | | | |
| | **EOQ** | (222) | C6-T-A-*l*-*i*-V | C30H55N5O8 | 613.4051 | 1.2 |
| | **EOR** | (223) | C5-T-A-*l*-*i*-L | C30H55N5O8 | 613.4051 | -0.3 |
| | **EOS** | | | | | |
| | **EOT** | | | | | |
| | **EOU** | | | | | |
| | **EOV** | | | | | |
| | **EPQ** | | | | | |
| | **EPR** | | | | | |
| | **EPS** | | | | | |
| | **EPT** | (224) | C6-T-A-bA-Y-*v*-V | C35H56N6O10 | 720.4058 | 1.1 |
| | | (225) | C5-T-A-bA-Y-*v*-V | C34H54N6O10 | 706.3901 | 1.7 |
| | **EPU** | (226) | C6-cyclo[T-A-bA-*y*-Y-F] | C43H54N6O10 | 814.3901 | 1.5 |
| | | (227) | C6-T-A-bA-*y*-Y-F | C43H56N6O11 | 832.4007 | 1.4 |
| | | (228) | C5-T-A-bA-*y*-Y-F | C42H54N6O11 | 818.3851 | 1.5 |
| | | (229) | C6-cyclo[T-A-bA-*y*-Y-W] | C45H55N7O10 | 853.4010 | 1.7 |
| | | (230) | C6-T-A-bA-*y*-Y-W | C45H57N7O11 | 871.4116 | 1.1 |
| | **EPV** | | | | | |
| | **FHQ** | (231) | C14-*q*-N-*v*-V | C33H60N6O8 | 668.4473 | -0.1 |
| | **FHR** | (232) | C14-*q*-N-*v*-L | C34H62N6O8 | 682.4629 | 0.4 |
| | **FHS** | | | | | |
| | **FHT** | (233) | C14-*q*-N-V-*v*-V | C38H69N7O9 | 767.5157 | 0.9 |
| | **FHU** | (234) | C14-*q*-N-*v*-Y-F | C46H69N7O10 | 879.5106 | 0.4 |
| | | (235) | C14-*q*-N-*v*-Y-W | C48H70N8O10 | 918.5215 | -1.2 |
| | **FHV** | | | | | |
| | **FIQ** | (236) | C14-*q*-N-*s*-V | C31H56N6O9 | 656.4109 | -0.7 |
| | **FIR** | (237) | C14-*q*-N-*s*-L | C32H58N6O9 | 670.4265 | -0.3 |
| | **FIS** | (238) | C14-*q*-N-*s*-K | C32H59N7O9 | 685.4374 | 0.6 |
| | **FIT** | (239) | C14-*q*-N-S-*v*-V | C36H65N7O10 | 755.4793 | 0.1 |
| | **FIU** | | | | | |
| | **FIV** | (240) | C14-*q*-N-S-*f*-*l*-L | C47H78N8O11 | 930.5790 | 0.7 |
| | **FJQ** | (241) | C14-*q*-*n*-*y*-V | C37H60N6O9 | 732.4422 | -0.1 |
| | **FJR** | (242) | C14-*q*-*n*-*y*-L | C38H62N6O9 | 746.4578 | -0.7 |
| | **FJS** | | | | | |
| | **FJT** | (243) | C14-*q*-*n*-Y-*v*-V | C42H69N7O10 | 831.5106 | 0.1 |
| | **FJU** | (244) | C14-*q*-*n*-*y*-Y-F | C50H69N7O11 | 943.5055 | 1.0 |
| | | (245) | C14-*q*-*n*-*y*-Y-W | C52H70N8O11 | 982.5164 | 1.0 |
| | **FJV** | (246) | C14-*q*-*n*-Y-*f*-*l*-L | C53H82N8O11 | 1006.6103 | -0.2 |

| | | | | | | |
|---|---|---|---|---|---|---|
| | **FKQ** | (247) | C14-*q*-N-*f*-V | C37H60N6O8 | 716.4473 | -0.5 |
| | **FKR** | (248) | C14-*q*-N-*f*-L | C38H62N6O8 | 730.4629 | -0.4 |
| | **FKS** | | | | | |
| | **FKT** | | | | | |
| | **FKU** | | | | | |
| | **FKV** | | | | | |
| | **FLQ** | (249) | C14-*q*-N-*f*-*l*-V | C43H71N7O9 | 829.5313 | 0.6 |
| | **FLR** | (250) | C14-*q*-N-*f*-*l*-L | C44H73N7O9 | 843.5470 | -0.2 |
| | **FLS** | (251) | C14-*q*-N-*f*-*l*-K | C44H74N8O9 | 858.5579 | 0.5 |
| | **FLT** | | | | | |
| | **FLU** | (252) | C14-*q*-N-*f*-*l*-Y-F | C56H80N8O11 | 1040.5947 | 0.6 |
| | **FLV** | | | | | |
| | **FMQ** | | | | | |
| | **FMR** | | | | | |
| | **FMS** | | | | | |
| | **FMT** | | | | | |
| | **FMU** | | | | | |
| | **FMV** | | | | | |
| | **FNQ** | (253) | C14-*q*-N-*w*-*l*-V | C45H72N8O9 | 868.5422 | 0.4 |
| | **FNR** | | | | | |
| | **FNS** | | | | | |
| | **FNT** | | | | | |
| | **FNU** | | | | | |
| | **FNV** | | | | | |
| | **FOQ** | (254) | C14-*q*-N-*l*-*i*-V | C40H73N7O9 | 795.5470 | -0.8 |
| | **FOR** | (255) | C14-*q*-N-*l*-*i*-L | C41H75N7O9 | 809.5626 | 0.1 |
| | **FOS** | | | | | |
| | **FOT** | (**14**) | C14-*q*-N-*l*-I-*v*-V | C45H82N8O10 | 894.6154 | -0.6 |
| | **FOU** | | | | | |
| | **FOV** | | | | | |
| | **FPQ** | | | | | |
| | **FPR** | | | | | |
| | **FPS** | | | | | |
| | **FPT** | (256) | C14-*q*-N-bA-Y-*v*-V | C45H74N8O11 | 902.5477 | -0.1 |
| | **FPU** | (257) | C14-*q*-N-bA-*y*-Y-F | C53H74N8O12 | 1014.5426 | 0.9 |
| | | (258) | C14-*q*-N-bA-*y*-Y-W | C55H75N9O12 | 1053.5535 | 1.1 |
| | **FPV** | | | | | |

### SUPPLEMENTARY FIGURES

#### Figure S1. | Chemical structures and chromatograms of NRP compared with synthetic standards using HR-LC-MS/MS.

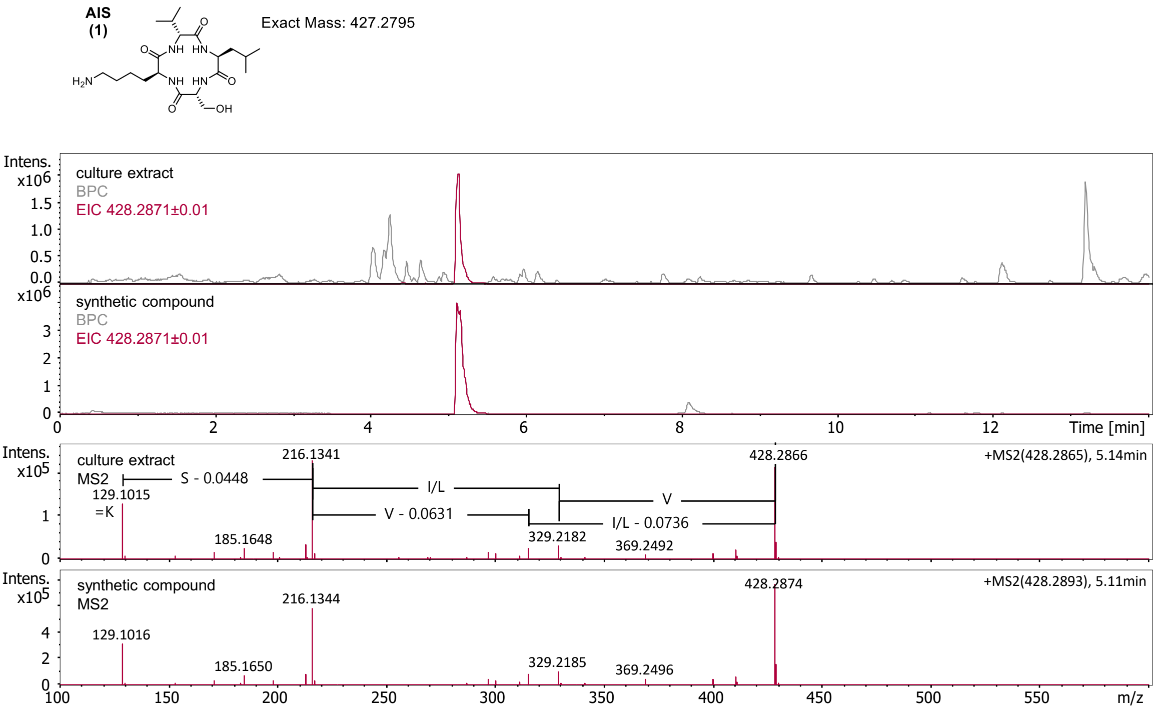

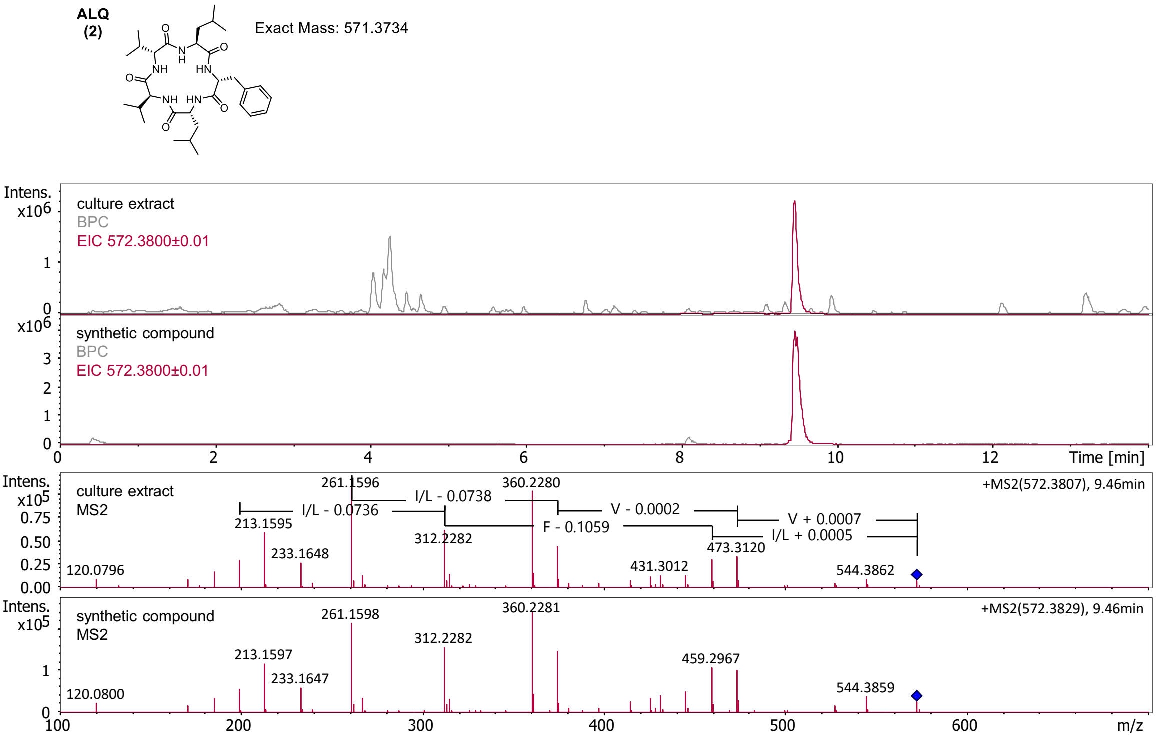

**AKV**
**(3)**

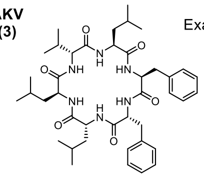

Exact Mass: 732.4574

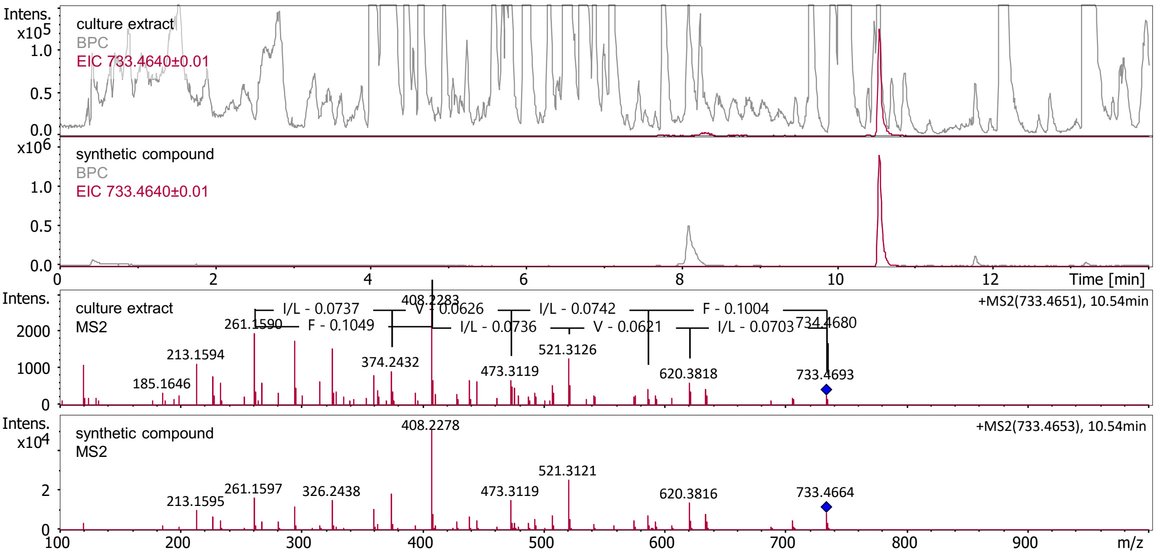

**APQ**
**(4)**

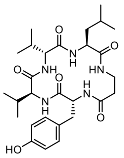

Exact Mass: 545.3213

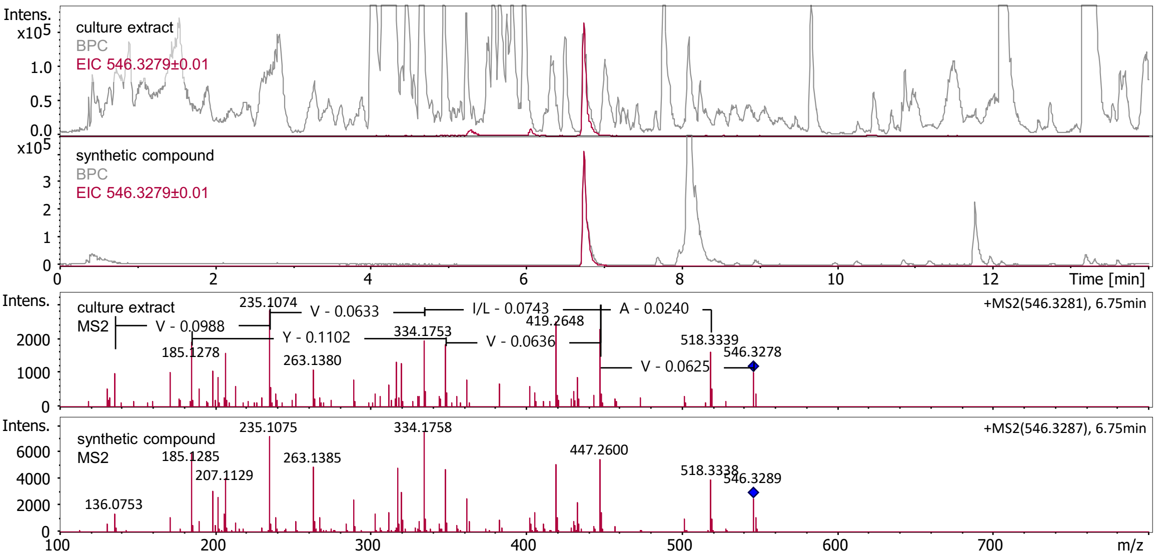

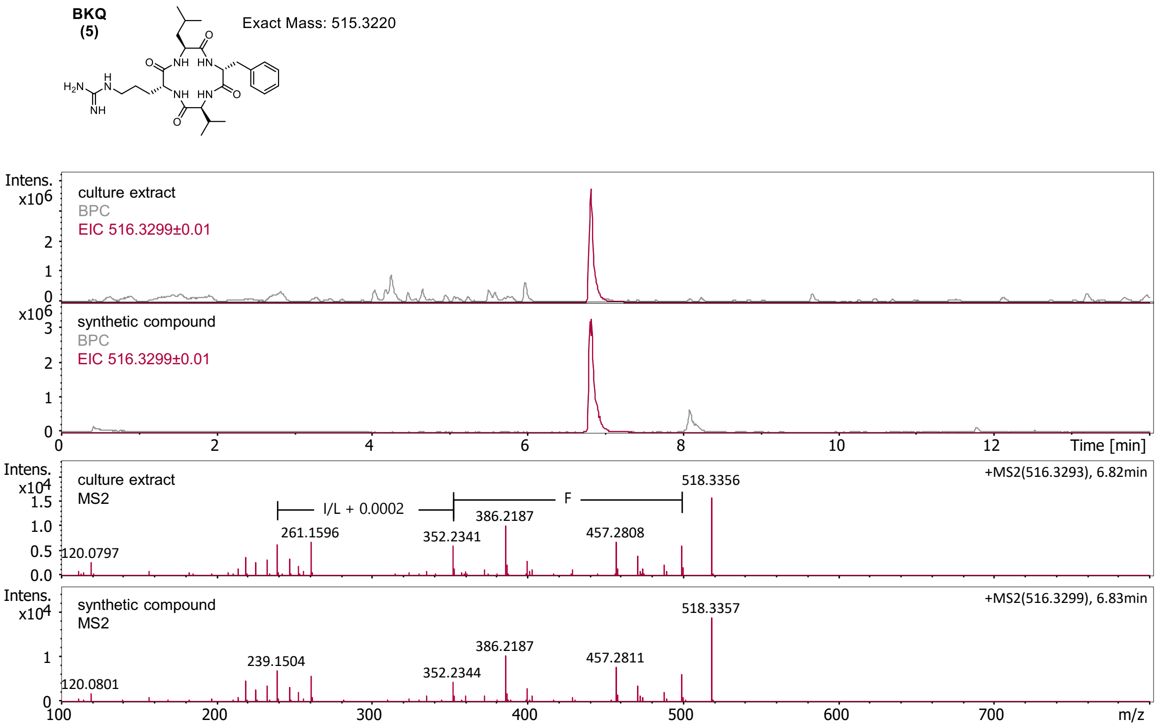
BKQ
(5)
Exact Mass: 515.3220
culture extract
BPC
EIC 516.3299±0.01
synthetic compound
BPC
EIC 516.3299±0.01
Time [min]
culture extract
MS2
+MS2(516.3293), 6.82min
I/L + 0.0002
F
518.3356
386.2187
261.1596
352.2341
457.2808
120.0797
synthetic compound
MS2
+MS2(516.3299), 6.83min
518.3357
386.2187
239.1504
352.2344
457.2811
120.0801
m/z

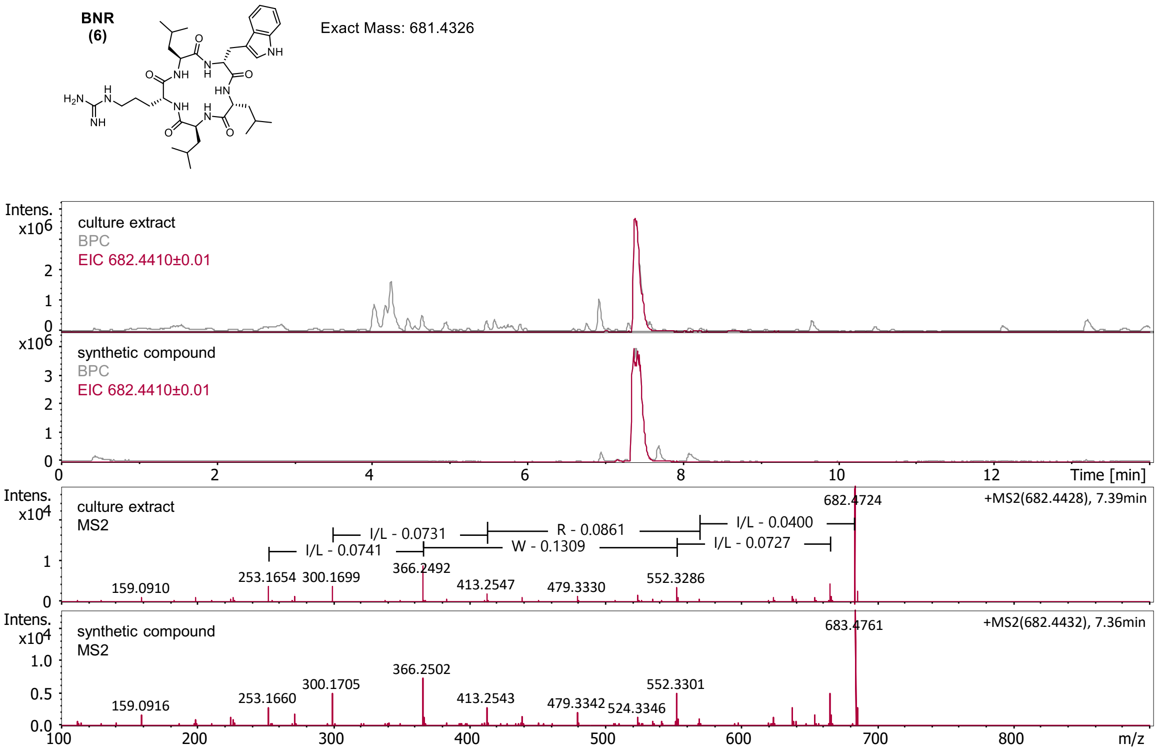
BNR
(6)
Exact Mass: 681.4326
culture extract
BPC
EIC 682.4410±0.01
synthetic compound
BPC
EIC 682.4410±0.01
Time [min]
culture extract
MS2
+MS2(682.4428), 7.39min
682.4724
I/L - 0.0731
R - 0.0861
I/L - 0.0400
I/L - 0.0741
W - 0.1309
I/L - 0.0727
366.2492
253.1654
300.1699
159.0910
413.2547
479.3330
552.3286
synthetic compound
MS2
+MS2(682.4432), 7.36min
683.4761
366.2502
300.1705
552.3301
253.1660
159.0916
413.2543
479.3342
524.3346
m/z

**BIV**
**(7)**

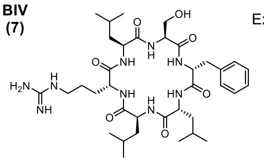

Exact Mass: 729.4537

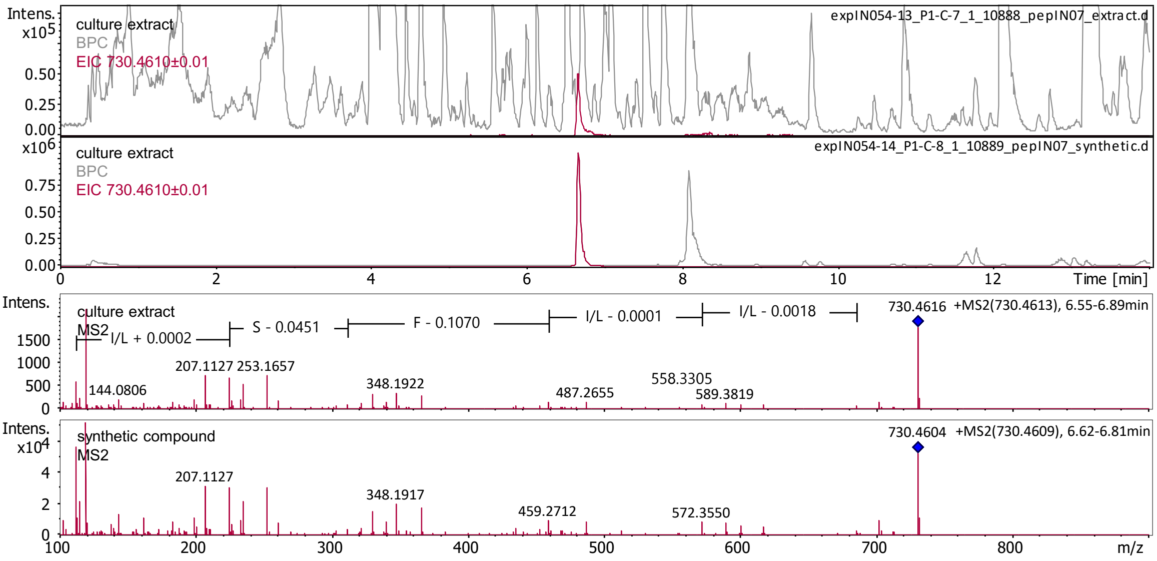

**BPU**
**(8)**

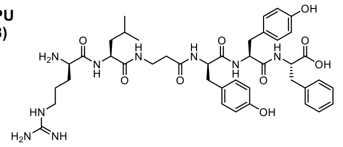

Exact Mass: 831.4279

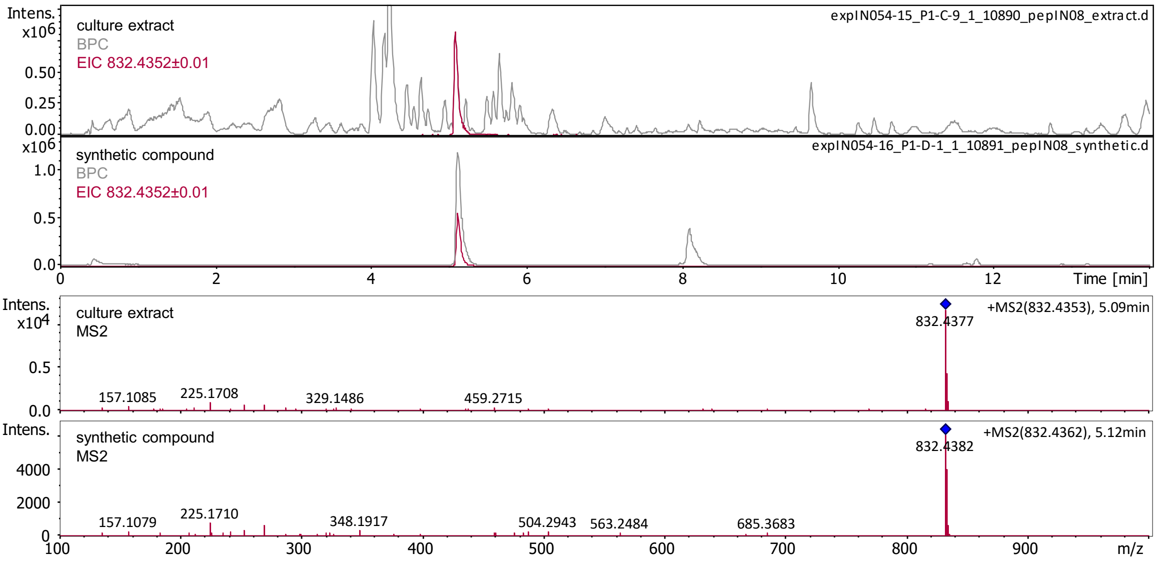

**CHR**
**(9)**

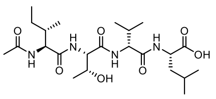

Exact Mass: 486.3053

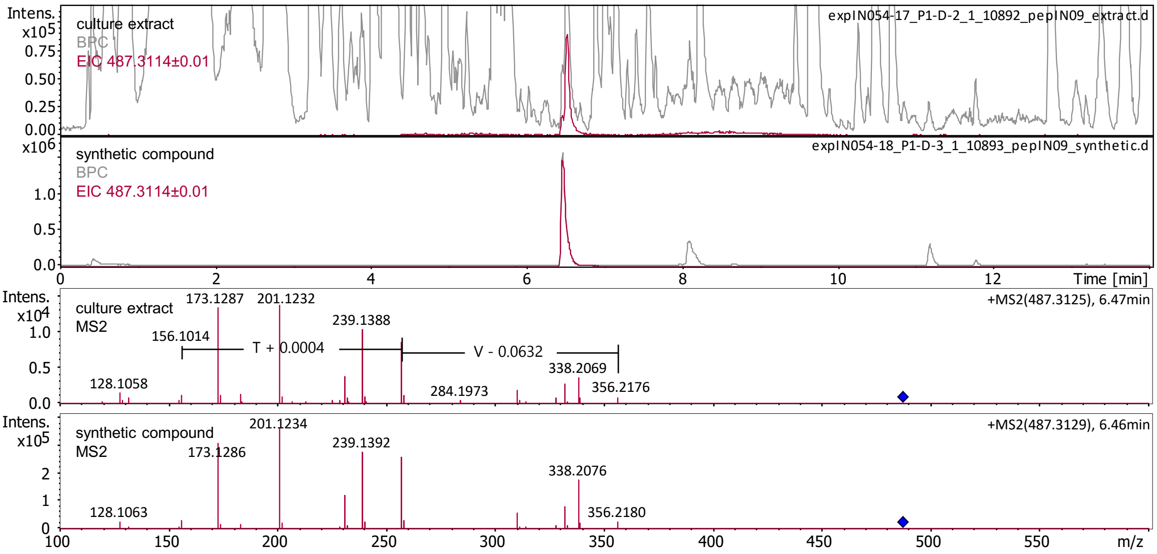

**CIV**
**(10)**

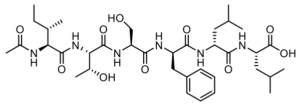

Exact Mass: 734.4214

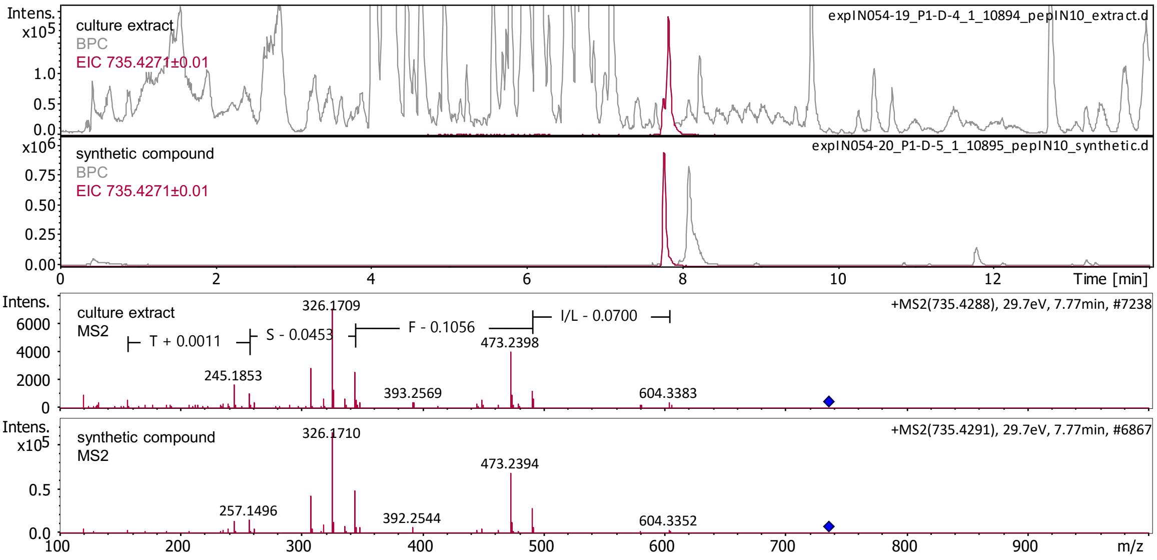

**DHU**
**(11)**

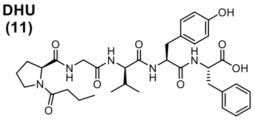

Exact Mass: 651.3268

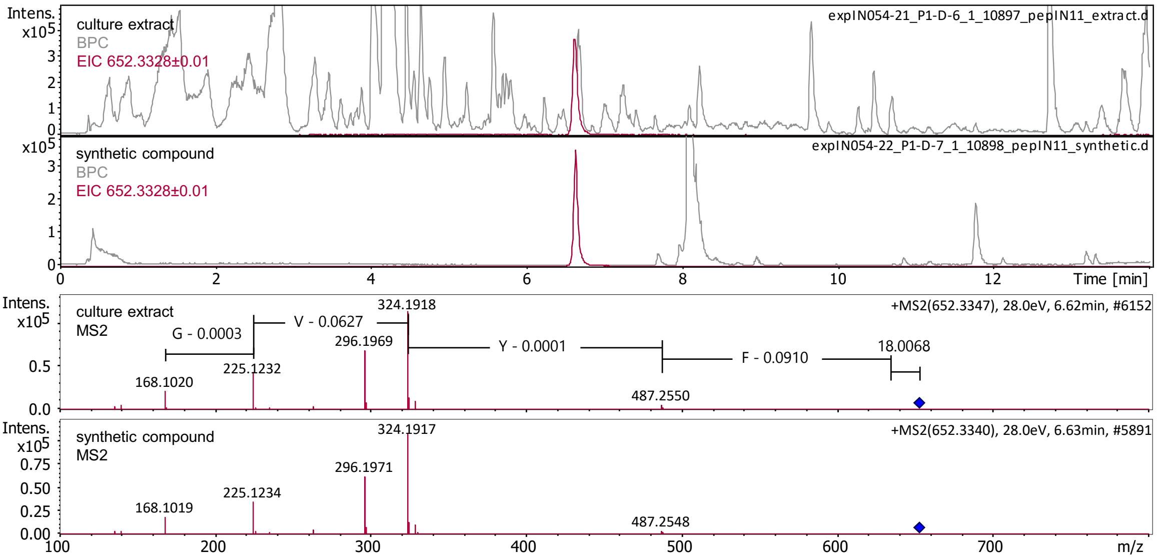

**DMV**
**(12)**

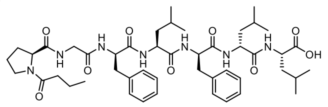

Exact Mass: 875.5157

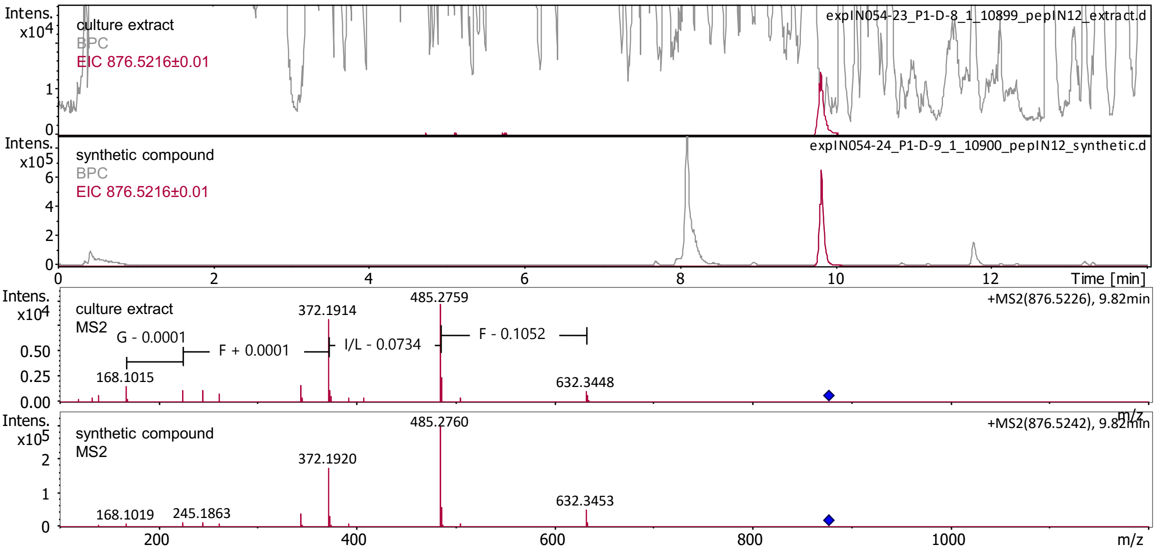

**EJT**
**(13)**

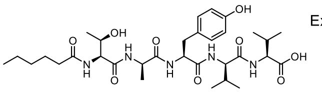

Exact Mass: 649.3687

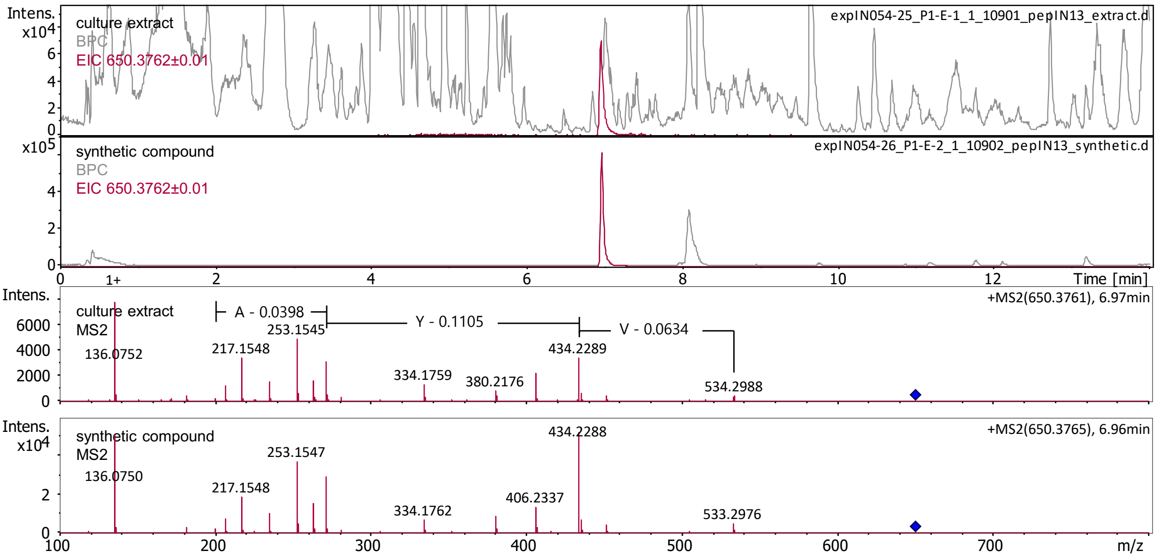

**FOT**
**(14)**

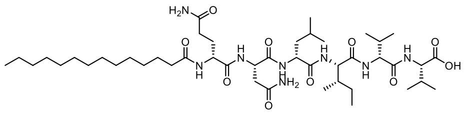

Exact Mass: 894.6154

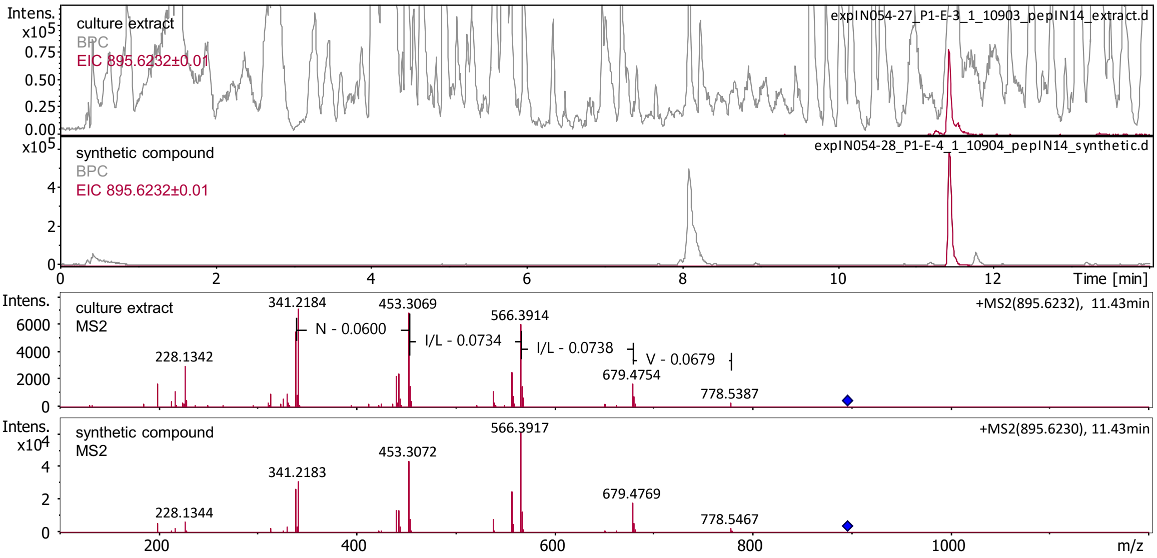

**Figure S2. | HR-LC-MS chromatograms of library peptides.** BPCs are shown in gray. EICs of expected compounds are displayed. In case multiple derivatives are produced,the first EIC of the expected *m/z* value is colorcoded in purple, blue (second), brown (third), green (fourth), and orange (fifth). The corresponding NRPS combinations and EIC values are indicated in the top right corner of each chromatogram.

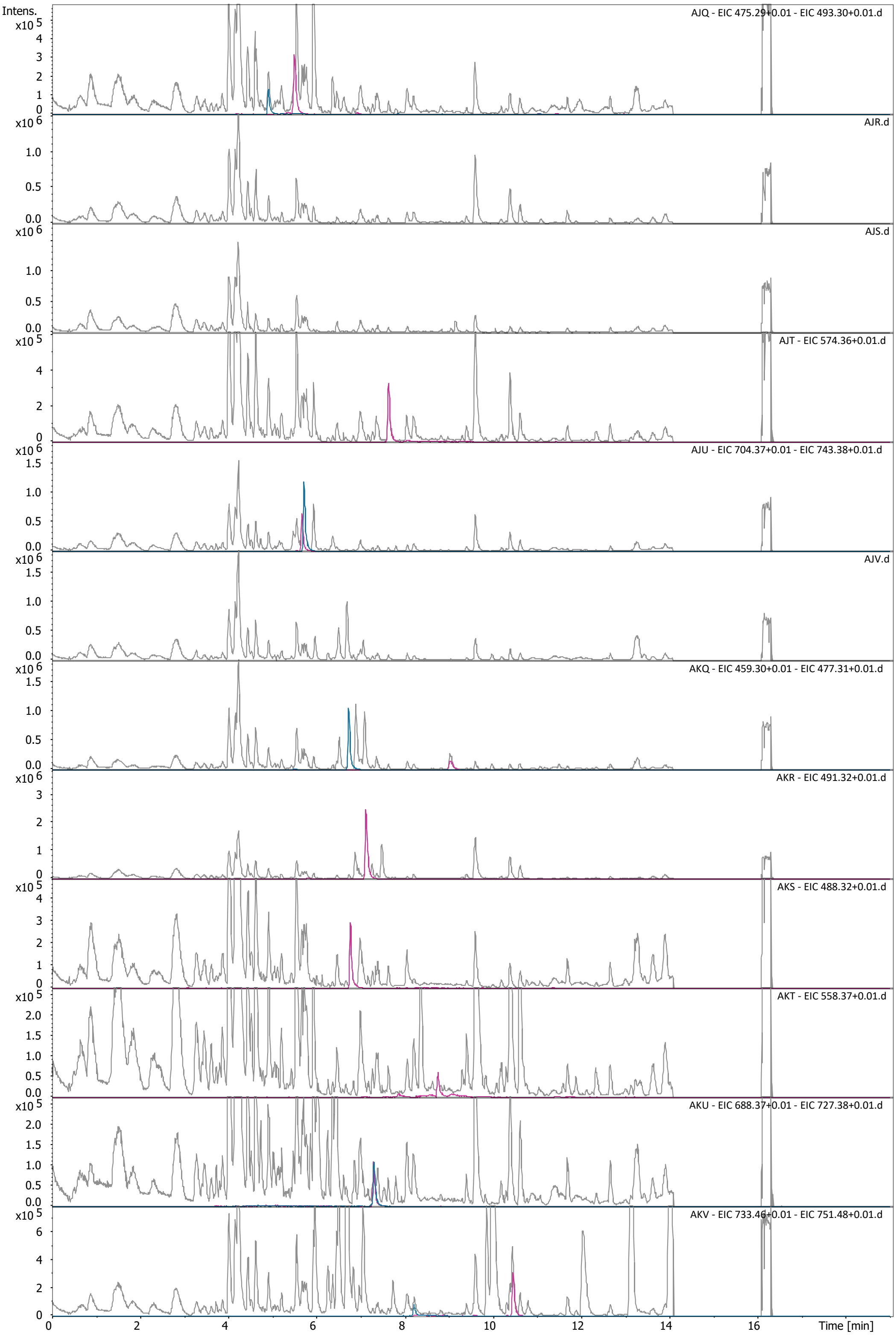

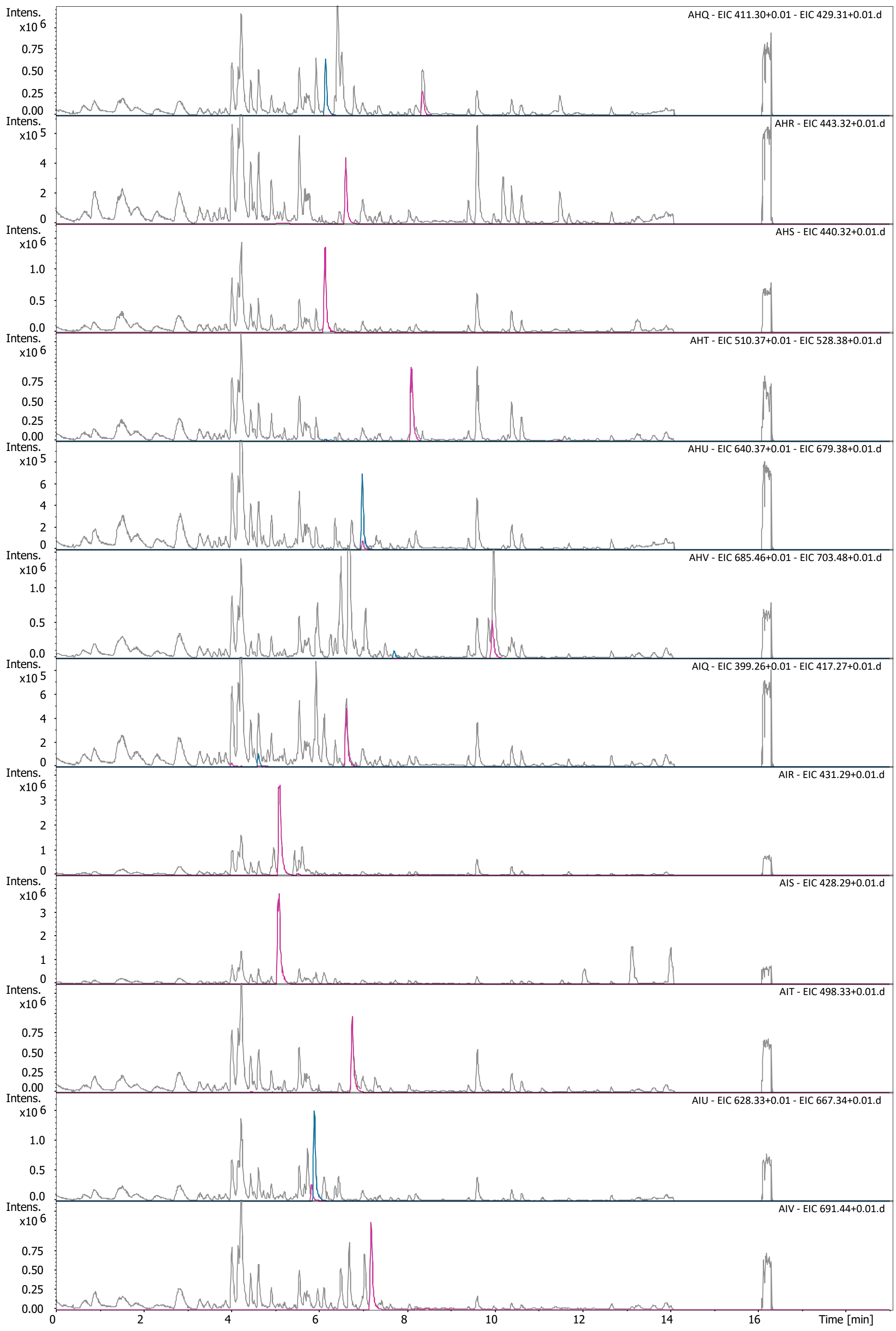
AHQ - EIC 411.30+0.01 - EIC 429.31+0.01.d
AHR - EIC 443.32+0.01.d
AHS - EIC 440.32+0.01.d
AHT - EIC 510.37+0.01 - EIC 528.38+0.01.d
AHU - EIC 640.37+0.01 - EIC 679.38+0.01.d
AHV - EIC 685.46+0.01 - EIC 703.48+0.01.d
AIQ - EIC 399.26+0.01 - EIC 417.27+0.01.d
AIR - EIC 431.29+0.01.d
AIS - EIC 428.29+0.01.d
AIT - EIC 498.33+0.01.d
AIU - EIC 628.33+0.01 - EIC 667.34+0.01.d
AIV - EIC 691.44+0.01.d
Intens.
Time [min]

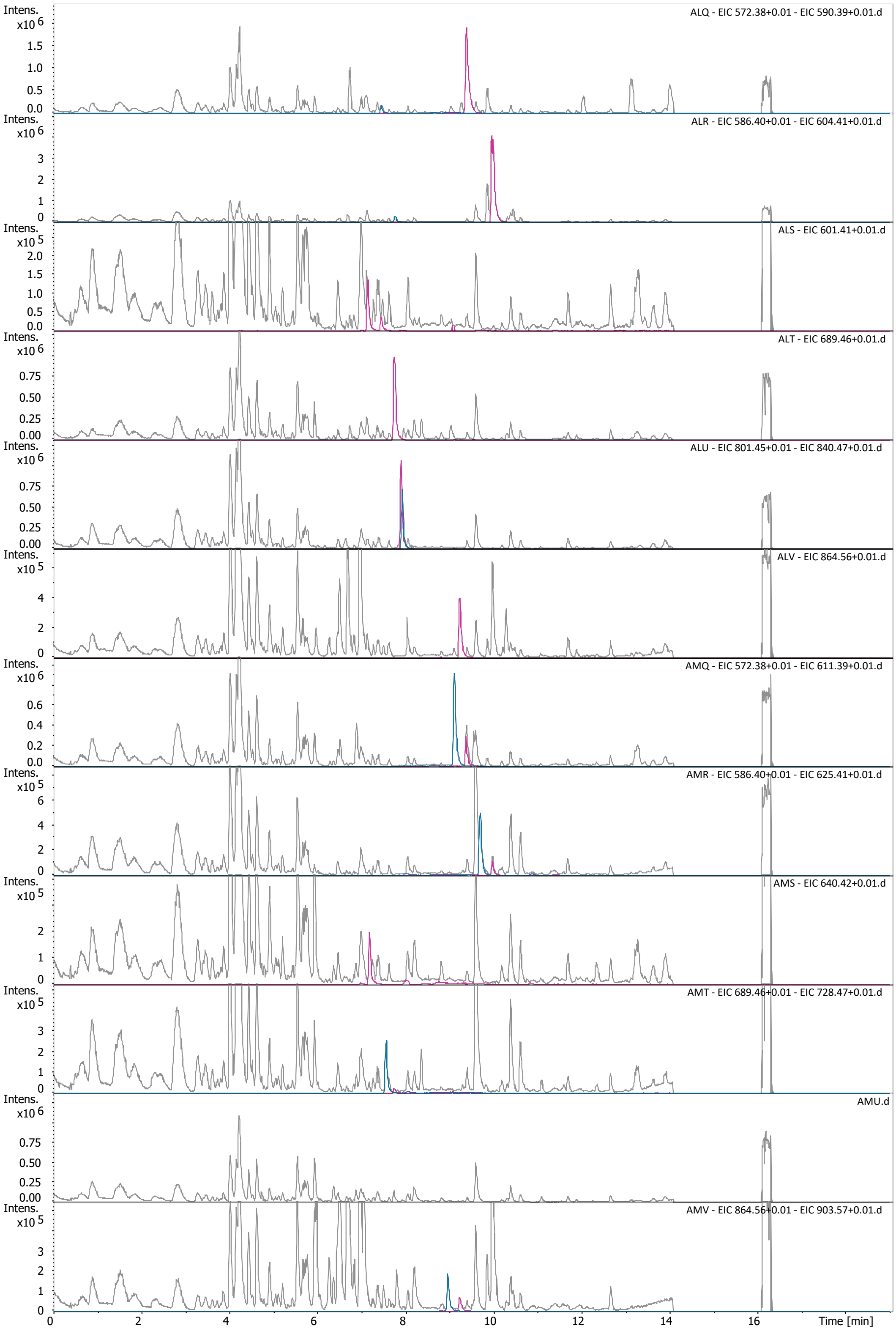
ALQ - EIC 572.38+0.01 - EIC 590.39+0.01.d
ALR - EIC 586.40+0.01 - EIC 604.41+0.01.d
ALS - EIC 601.41+0.01.d
ALT - EIC 689.46+0.01.d
ALU - EIC 801.45+0.01 - EIC 840.47+0.01.d
ALV - EIC 864.56+0.01.d
AMQ - EIC 572.38+0.01 - EIC 611.39+0.01.d
AMR - EIC 586.40+0.01 - EIC 625.41+0.01.d
AMS - EIC 640.42+0.01.d
AMT - EIC 689.46+0.01 - EIC 728.47+0.01.d
AMU.d
AMV - EIC 864.56+0.01 - EIC 903.57+0.01.d
Intens.
Time [min]

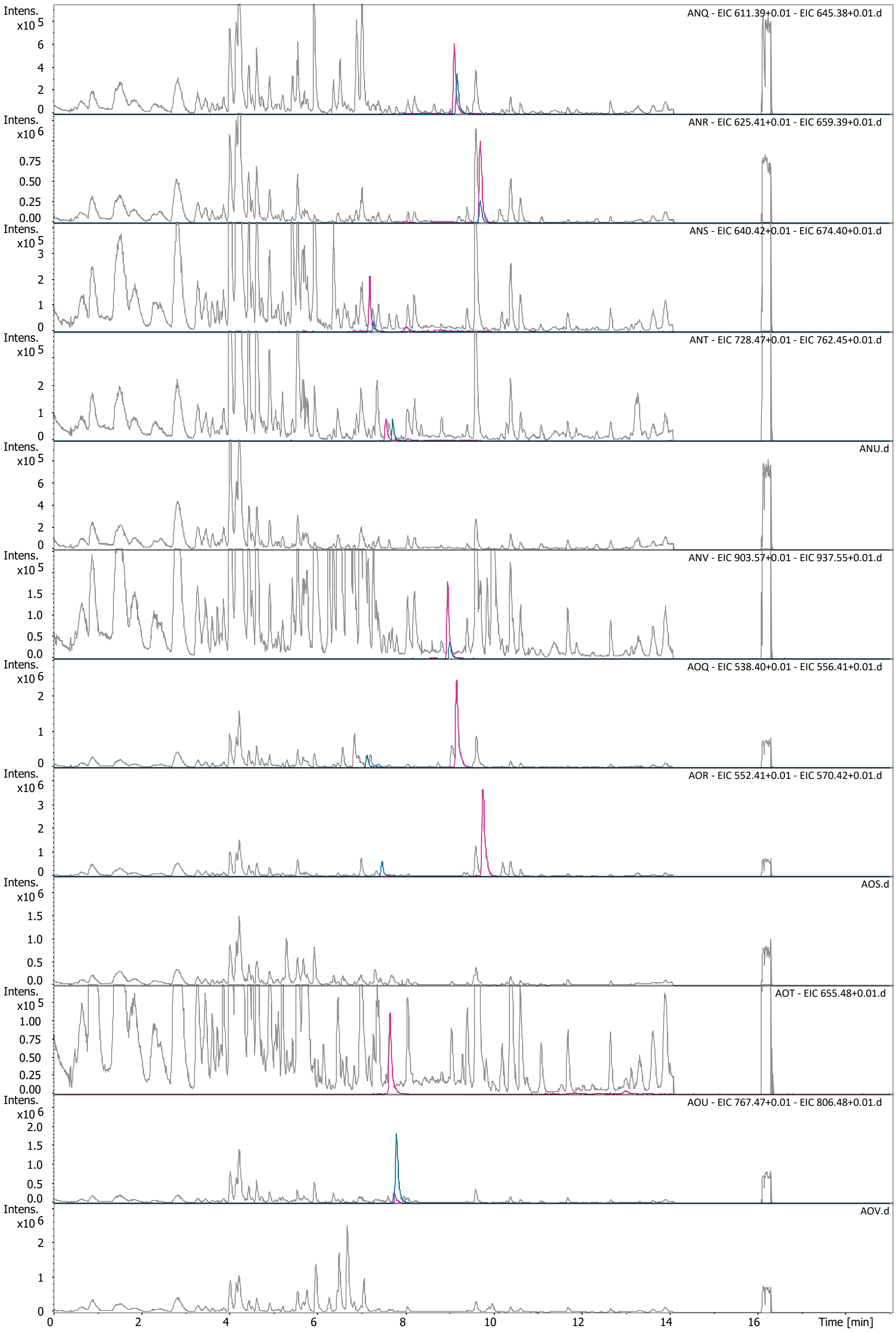
Intens.
x10 5
ANQ - EIC 611.39+0.01 - EIC 645.38+0.01.d
ANR - EIC 625.41+0.01 - EIC 659.39+0.01.d
ANS - EIC 640.42+0.01 - EIC 674.40+0.01.d
ANT - EIC 728.47+0.01 - EIC 762.45+0.01.d
ANU.d
ANV - EIC 903.57+0.01 - EIC 937.55+0.01.d
AOQ - EIC 538.40+0.01 - EIC 556.41+0.01.d
AOR - EIC 552.41+0.01 - EIC 570.42+0.01.d
AOS.d
AOT - EIC 655.48+0.01.d
AOU - EIC 767.47+0.01 - EIC 806.48+0.01.d
AOV.d
Time [min]

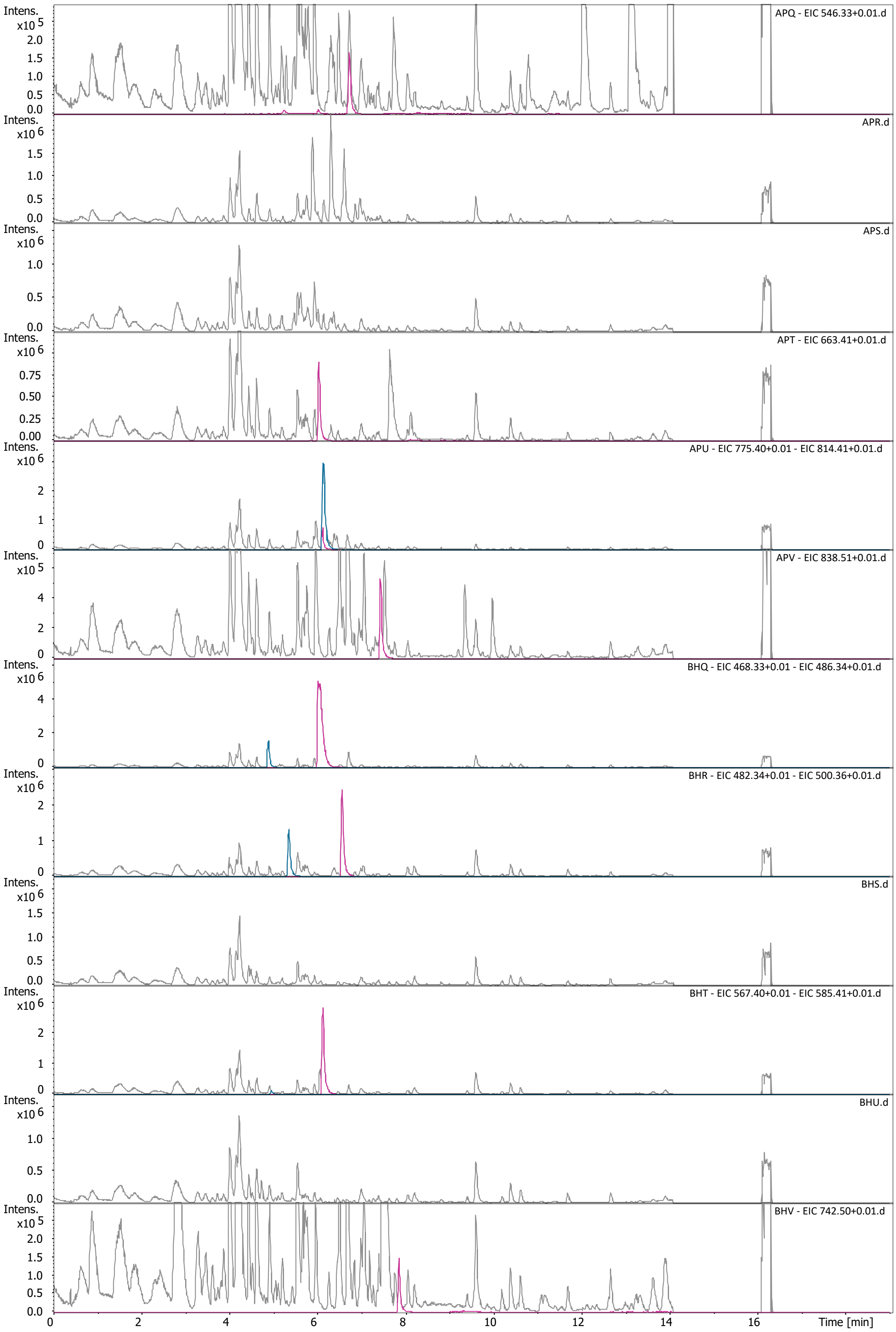

APQ - EIC 546.33+0.01.d
APR.d
APS.d
APT - EIC 663.41+0.01.d
APU - EIC 775.40+0.01 - EIC 814.41+0.01.d
APV - EIC 838.51+0.01.d
BHQ - EIC 468.33+0.01 - EIC 486.34+0.01.d
BHR - EIC 482.34+0.01 - EIC 500.36+0.01.d
BHS.d
BHT - EIC 567.40+0.01 - EIC 585.41+0.01.d
BHU.d
BHV - EIC 742.50+0.01.d
Intens.
Time [min]

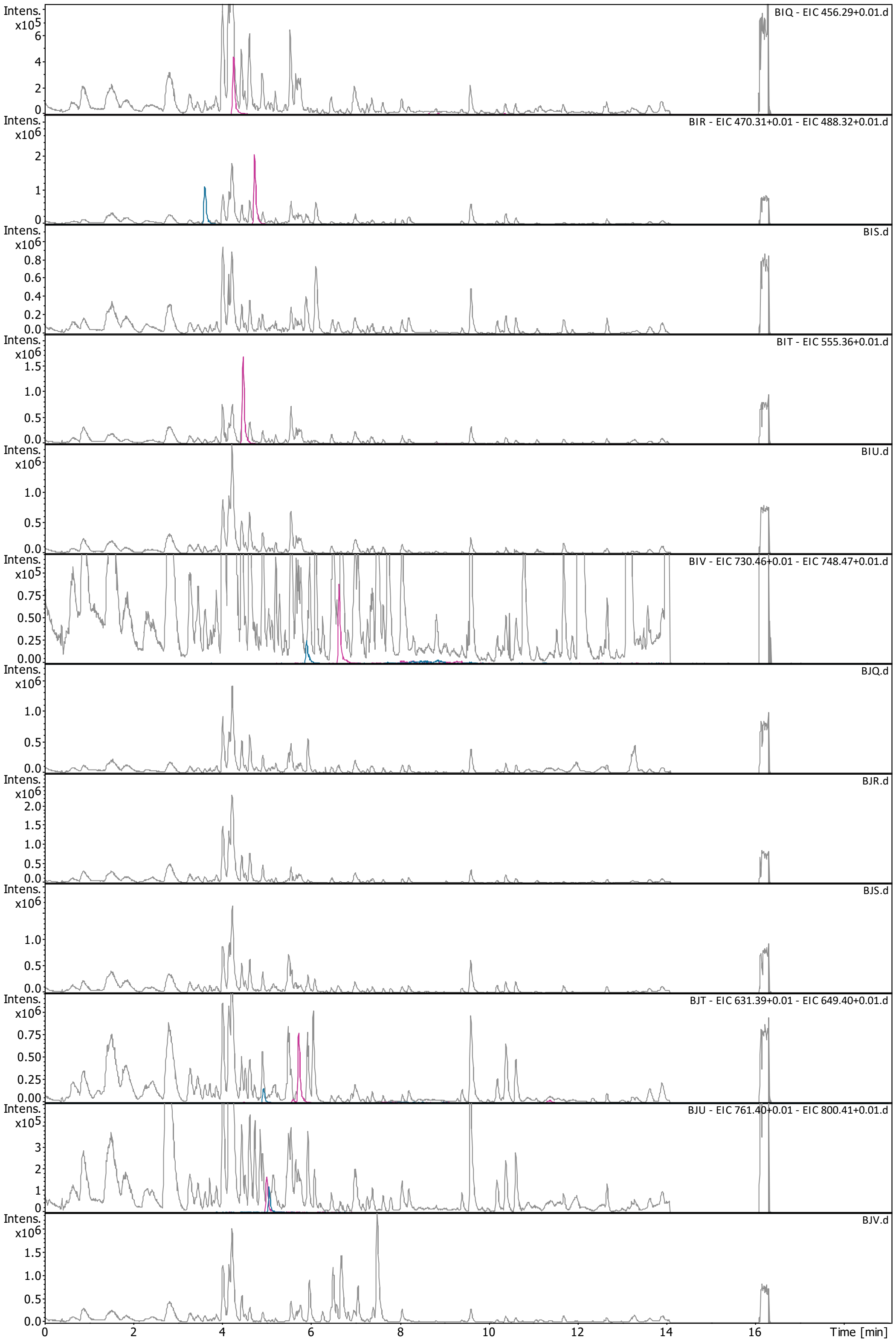
BIQ - EIC 456.29+0.01.d
BIR - EIC 470.31+0.01 - EIC 488.32+0.01.d
BIS.d
BIT - EIC 555.36+0.01.d
BIU.d
BIV - EIC 730.46+0.01 - EIC 748.47+0.01.d
BJQ.d
BJR.d
BJS.d
BJT - EIC 631.39+0.01 - EIC 649.40+0.01.d
BJU - EIC 761.40+0.01 - EIC 800.41+0.01.d
BJV.d
Intens.
Time [min]

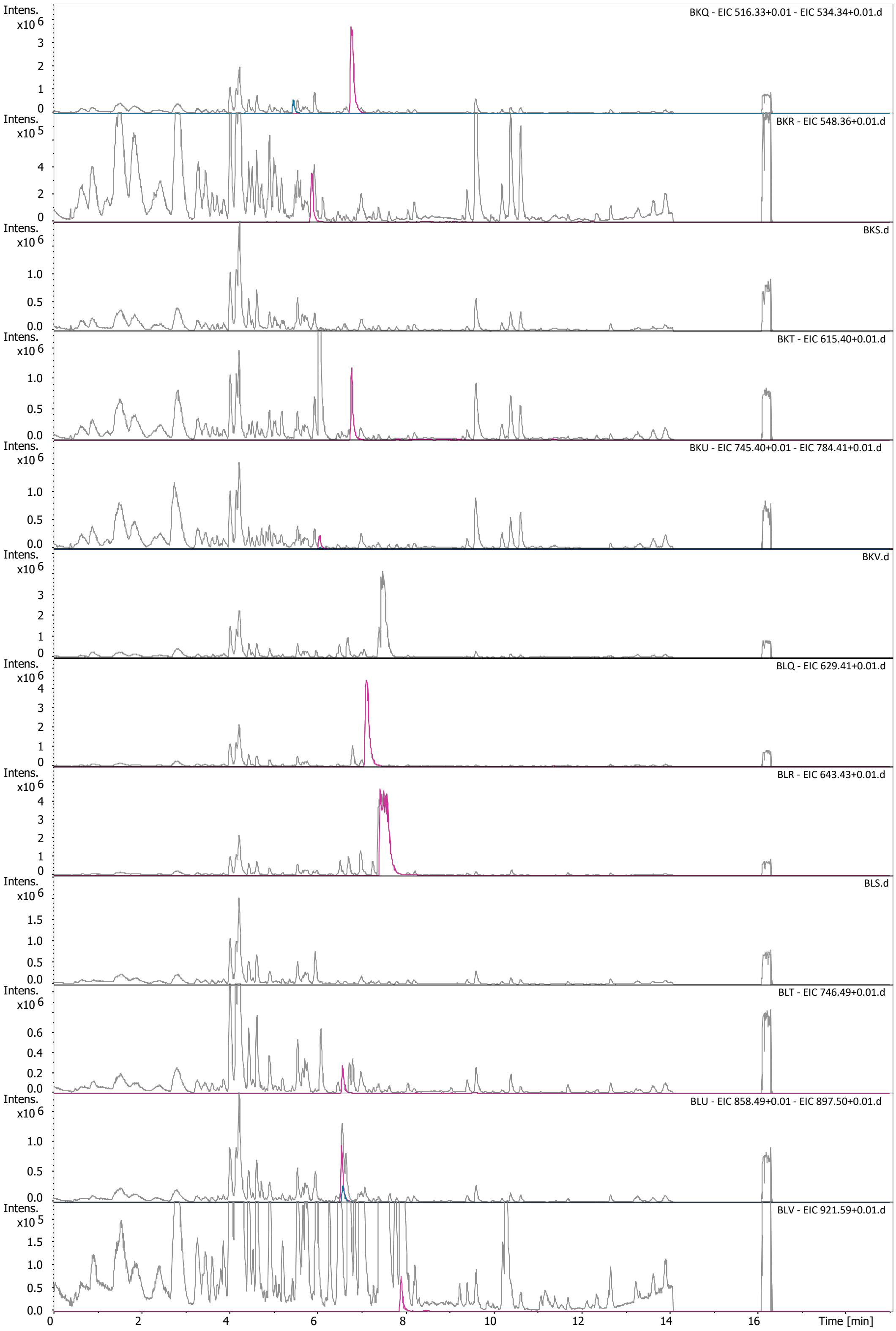
BKQ - EIC 516.33+0.01 - EIC 534.34+0.01.d
BKR - EIC 548.36+0.01.d
BKS.d
BKT - EIC 615.40+0.01.d
BKU - EIC 745.40+0.01 - EIC 784.41+0.01.d
BKV.d
BLQ - EIC 629.41+0.01.d
BLR - EIC 643.43+0.01.d
BLS.d
BLT - EIC 746.49+0.01.d
BLU - EIC 858.49+0.01 - EIC 897.50+0.01.d
BLV - EIC 921.59+0.01.d
Intens.
Time [min]

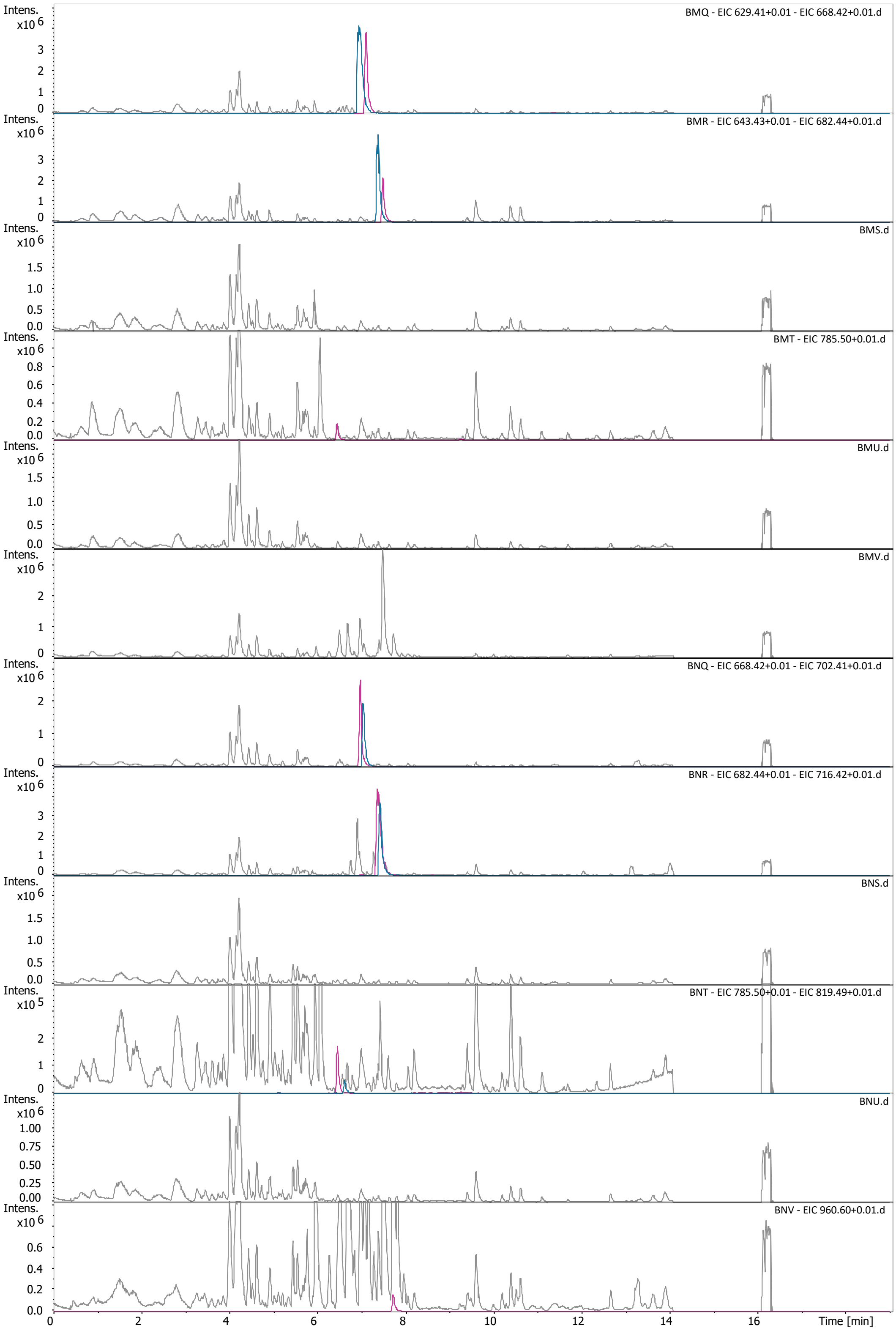

Intens. x10^6
BMQ - EIC 629.41+0.01 - EIC 668.42+0.01.d
BMR - EIC 643.43+0.01 - EIC 682.44+0.01.d
BMS.d
BMT - EIC 785.50+0.01.d
BMU.d
BMV.d
BNQ - EIC 668.42+0.01 - EIC 702.41+0.01.d
BNR - EIC 682.44+0.01 - EIC 716.42+0.01.d
BNS.d
Intens. x10^5
BNT - EIC 785.50+0.01 - EIC 819.49+0.01.d
BNU.d
BNV - EIC 960.60+0.01.d
Time [min]

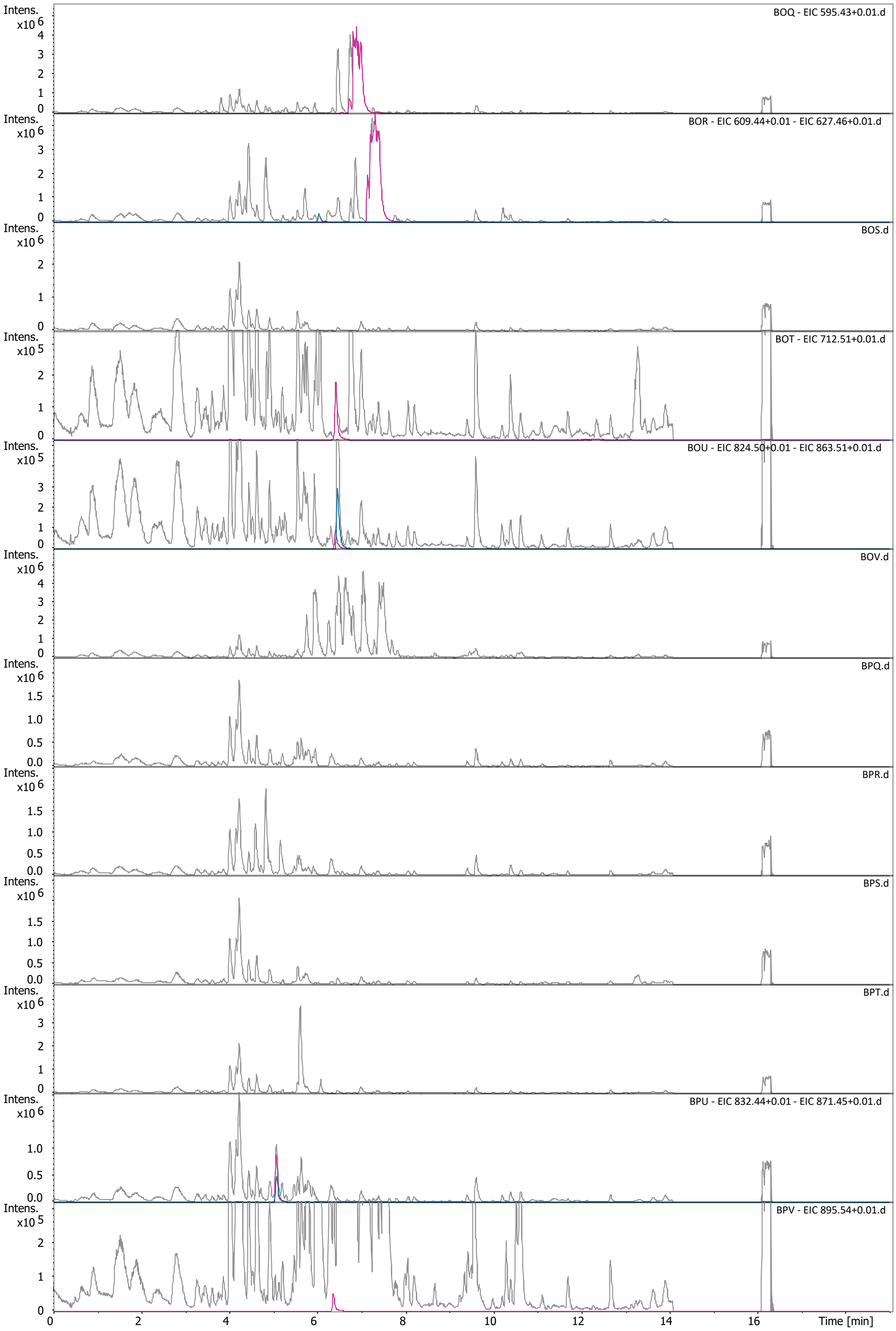

BOQ - EIC 595.43+0.01.d
BOR - EIC 609.44+0.01 - EIC 627.46+0.01.d
BOS.d
BOT - EIC 712.51+0.01.d
BOU - EIC 824.50+0.01 - EIC 863.51+0.01.d
BOV.d
BPQ.d
BPR.d
BPS.d
BPT.d
BPU - EIC 832.44+0.01 - EIC 871.45+0.01.d
BPV - EIC 895.54+0.01.d
Intens.
x10 6
x10 5
Time [min]

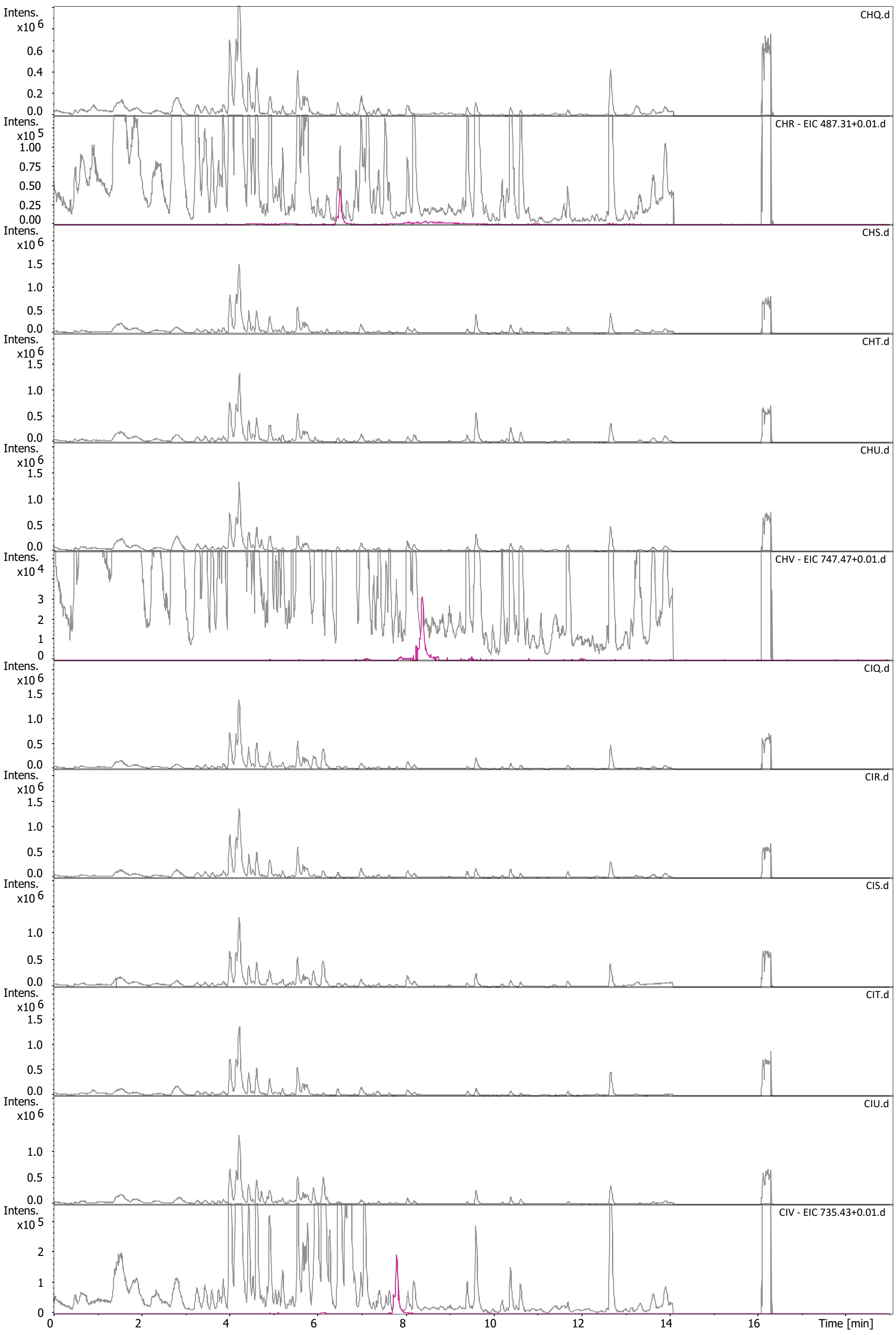

CHQ.d
CHR - EIC 487.31+0.01.d
CHS.d
CHT.d
CHU.d
CHV - EIC 747.47+0.01.d
CIQ.d
CIR.d
CIS.d
CIT.d
CIU.d
CIV - EIC 735.43+0.01.d
Intens.
Time [min]

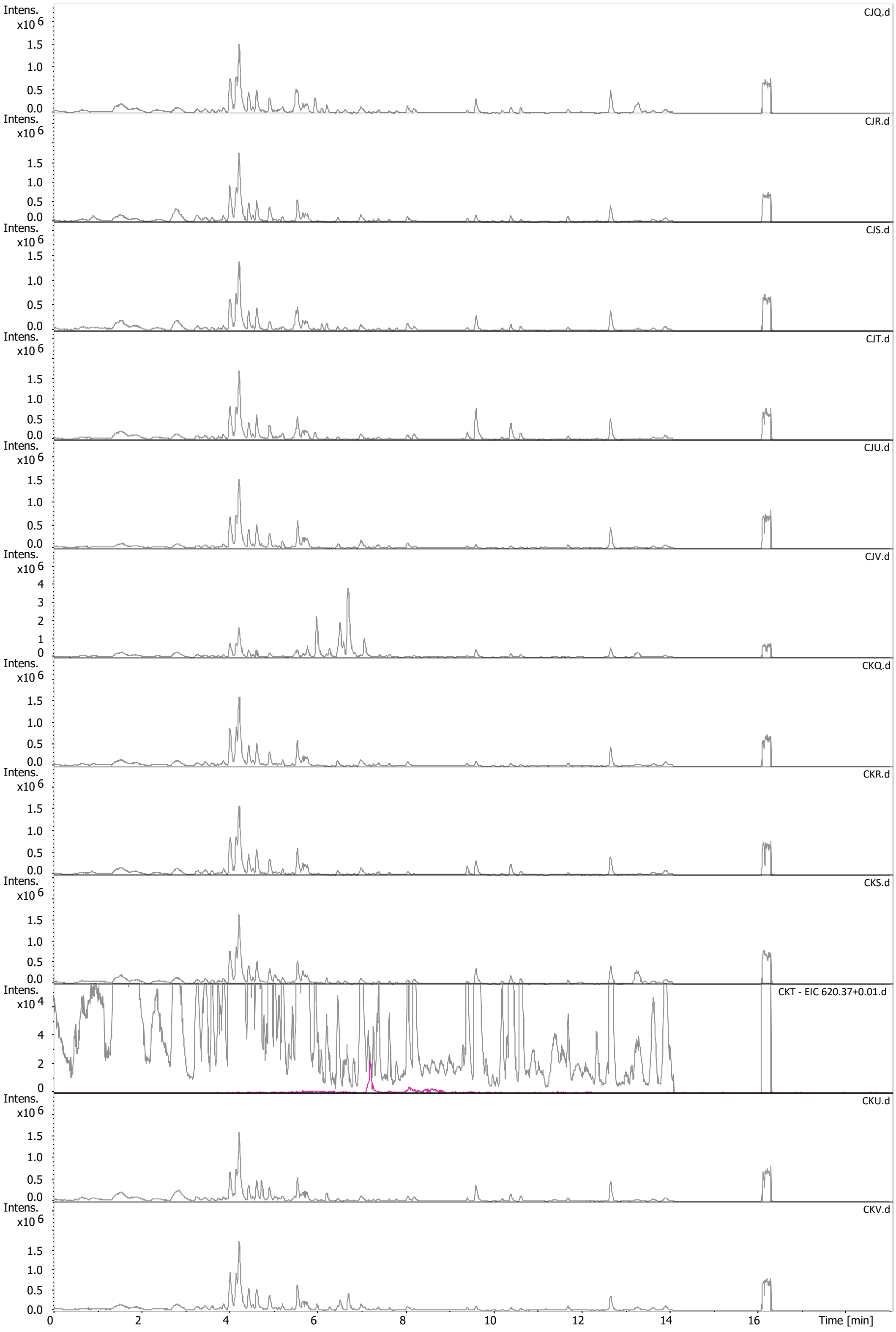

CJQ.d
CJR.d
CJS.d
CJT.d
CJU.d
CJV.d
CKQ.d
CKR.d
CKS.d
CKT - EIC 620.37+0.01.d
CKU.d
CKV.d
Intens.
x10 6
x10 4
Time [min]

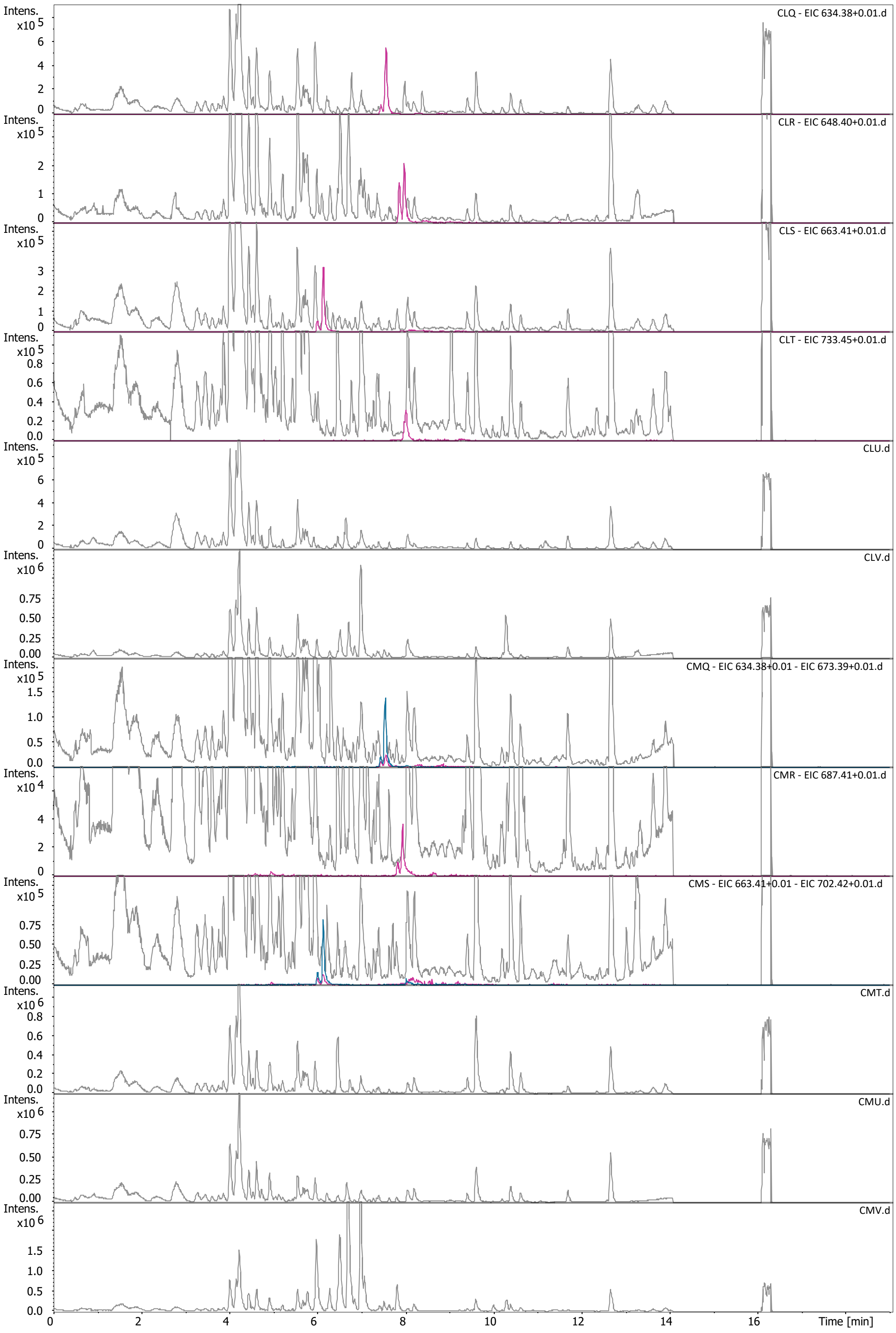
CLQ - EIC 634.38+0.01.d
CLR - EIC 648.40+0.01.d
CLS - EIC 663.41+0.01.d
CLT - EIC 733.45+0.01.d
CLU.d
CLV.d
CMQ - EIC 634.38+0.01 - EIC 673.39+0.01.d
CMR - EIC 687.41+0.01.d
CMS - EIC 663.41+0.01 - EIC 702.42+0.01.d
CMT.d
CMU.d
CMV.d
Intens.
Time [min]

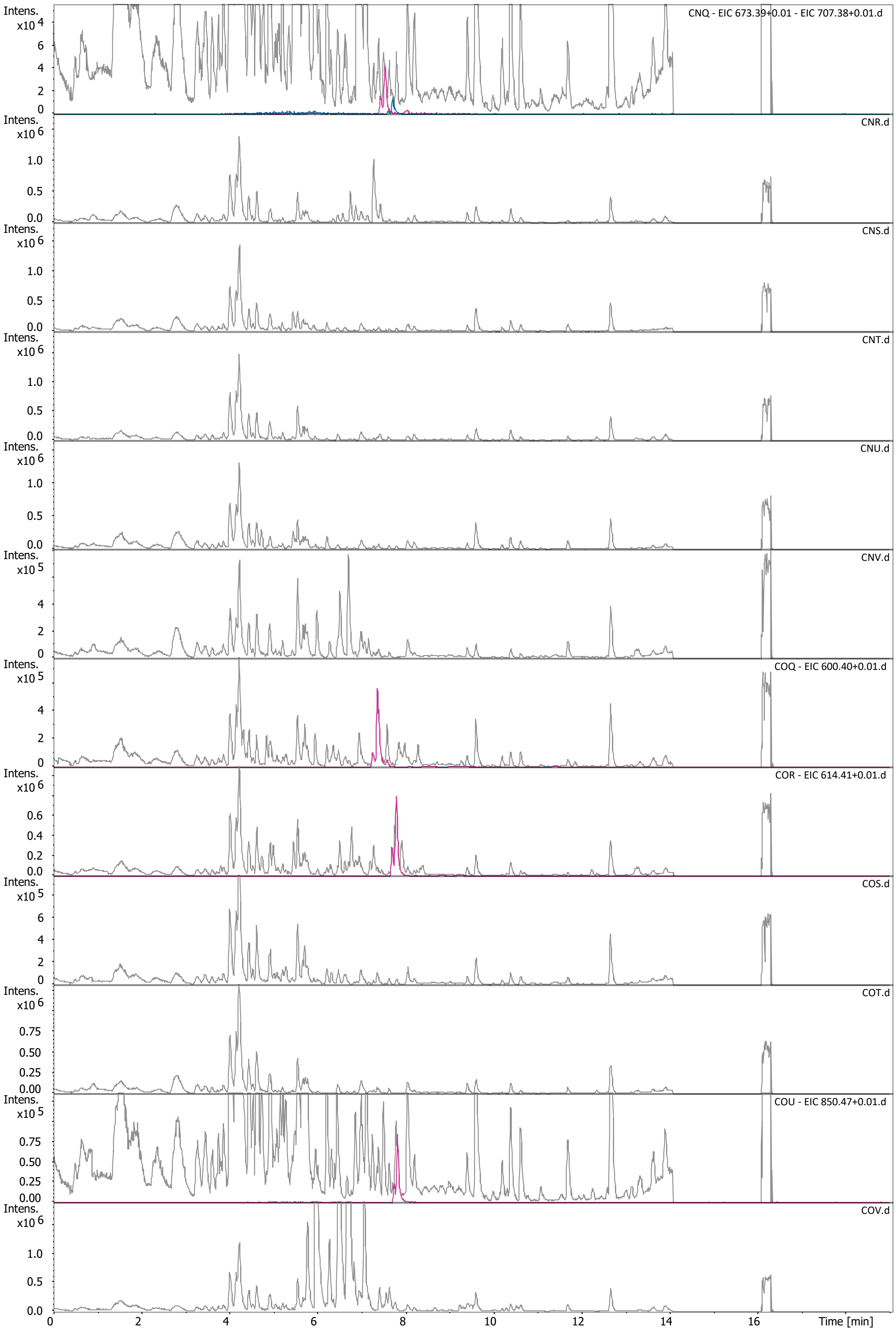
CNQ - EIC 673.39+0.01 - EIC 707.38+0.01.d
CNR.d
CNS.d
CNT.d
CNU.d
CNV.d
COQ - EIC 600.40+0.01.d
COR - EIC 614.41+0.01.d
COS.d
COT.d
COU - EIC 850.47+0.01.d
COV.d
Intens.
Time [min]

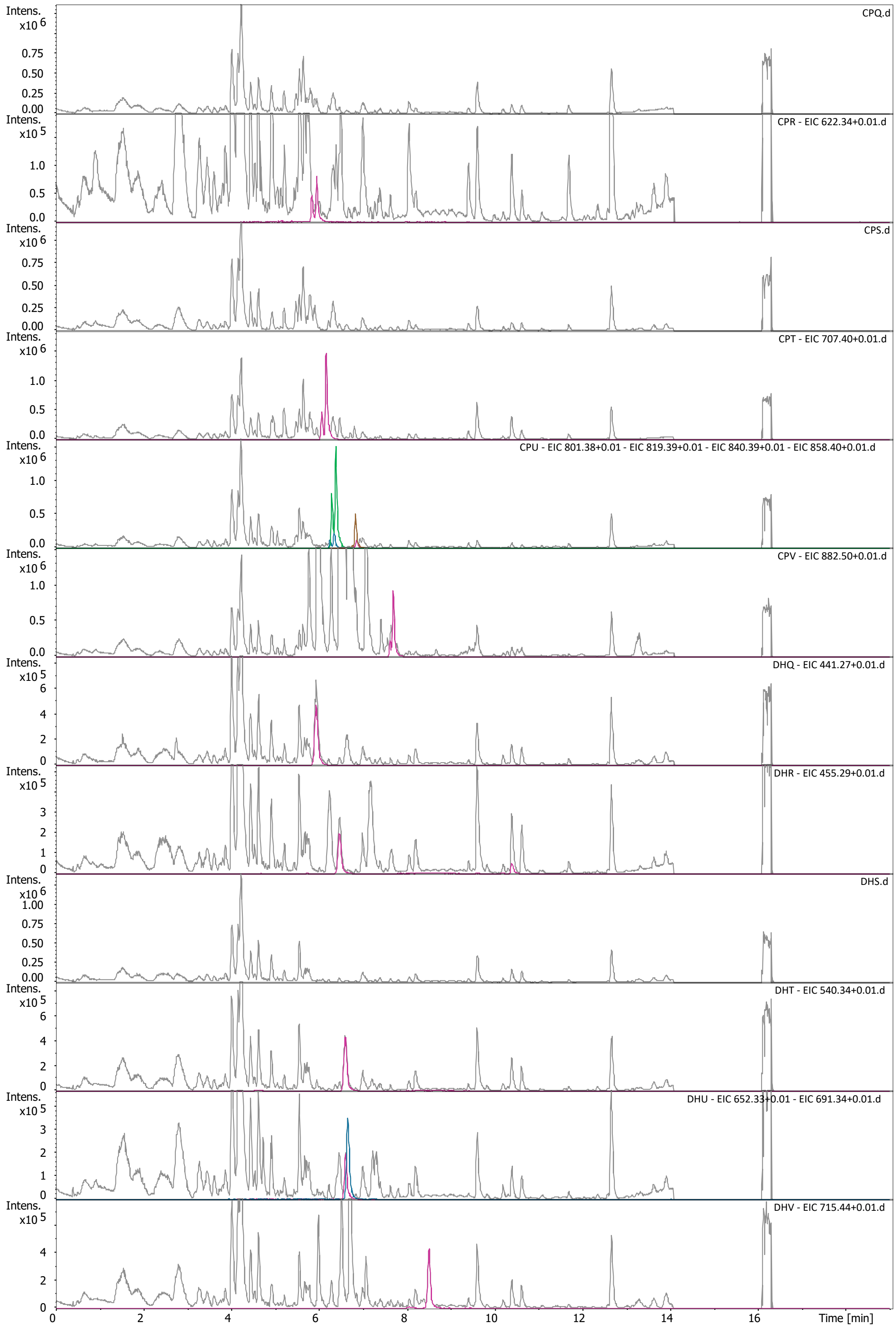
CPQ.d
CPR - EIC 622.34+0.01.d
CPS.d
CPT - EIC 707.40+0.01.d
CPU - EIC 801.38+0.01 - EIC 819.39+0.01 - EIC 840.39+0.01 - EIC 858.40+0.01.d
CPV - EIC 882.50+0.01.d
DHQ - EIC 441.27+0.01.d
DHR - EIC 455.29+0.01.d
DHS.d
DHT - EIC 540.34+0.01.d
DHU - EIC 652.33+0.01 - EIC 691.34+0.01.d
DHV - EIC 715.44+0.01.d
Intens.
Time [min]

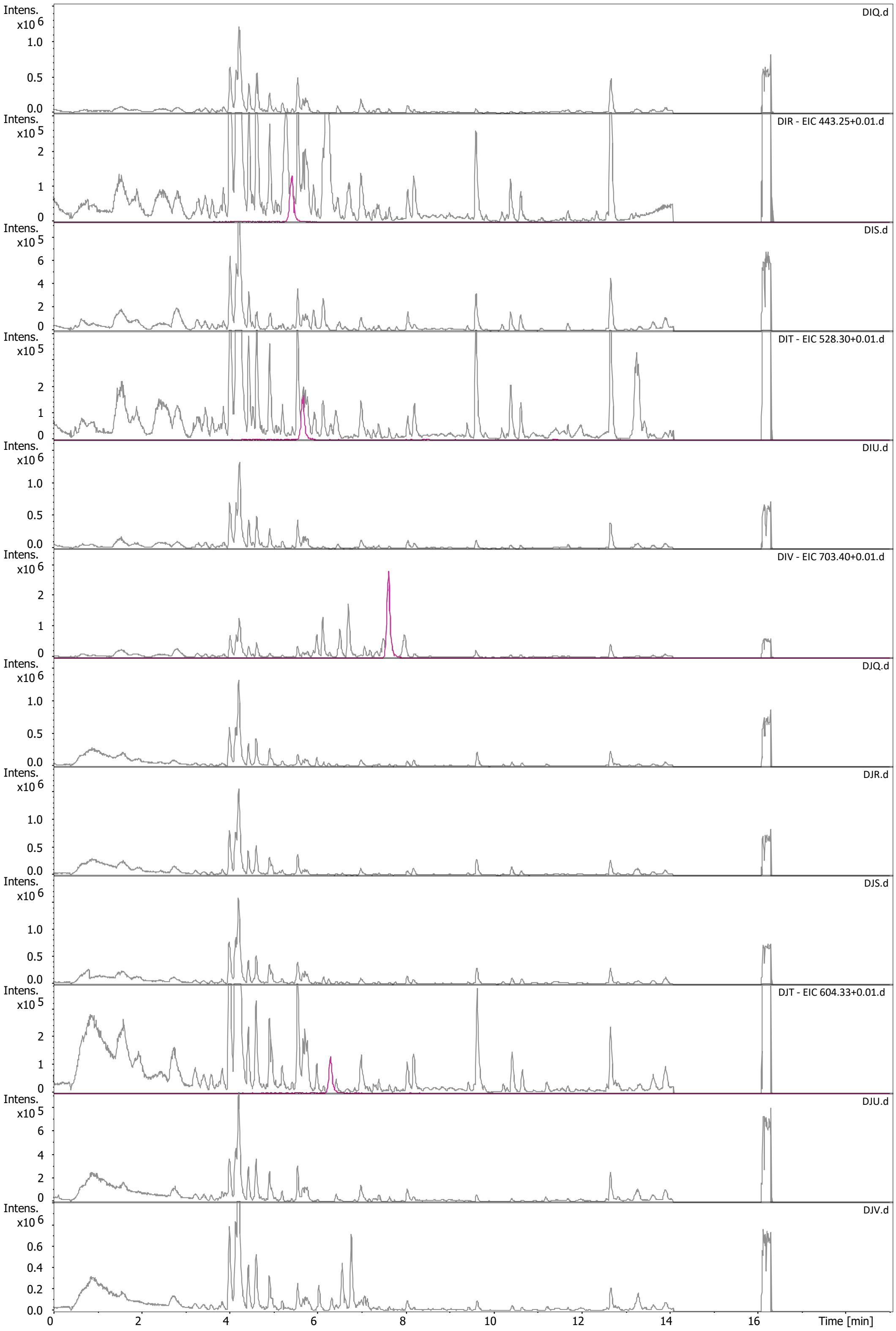
Intens.
x10 6
x10 5
DIQ.d
DIR - EIC 443.25+0.01.d
DIS.d
DIT - EIC 528.30+0.01.d
DIU.d
DIV - EIC 703.40+0.01.d
DJQ.d
DJR.d
DJS.d
DJT - EIC 604.33+0.01.d
DJU.d
DJV.d
Time [min]

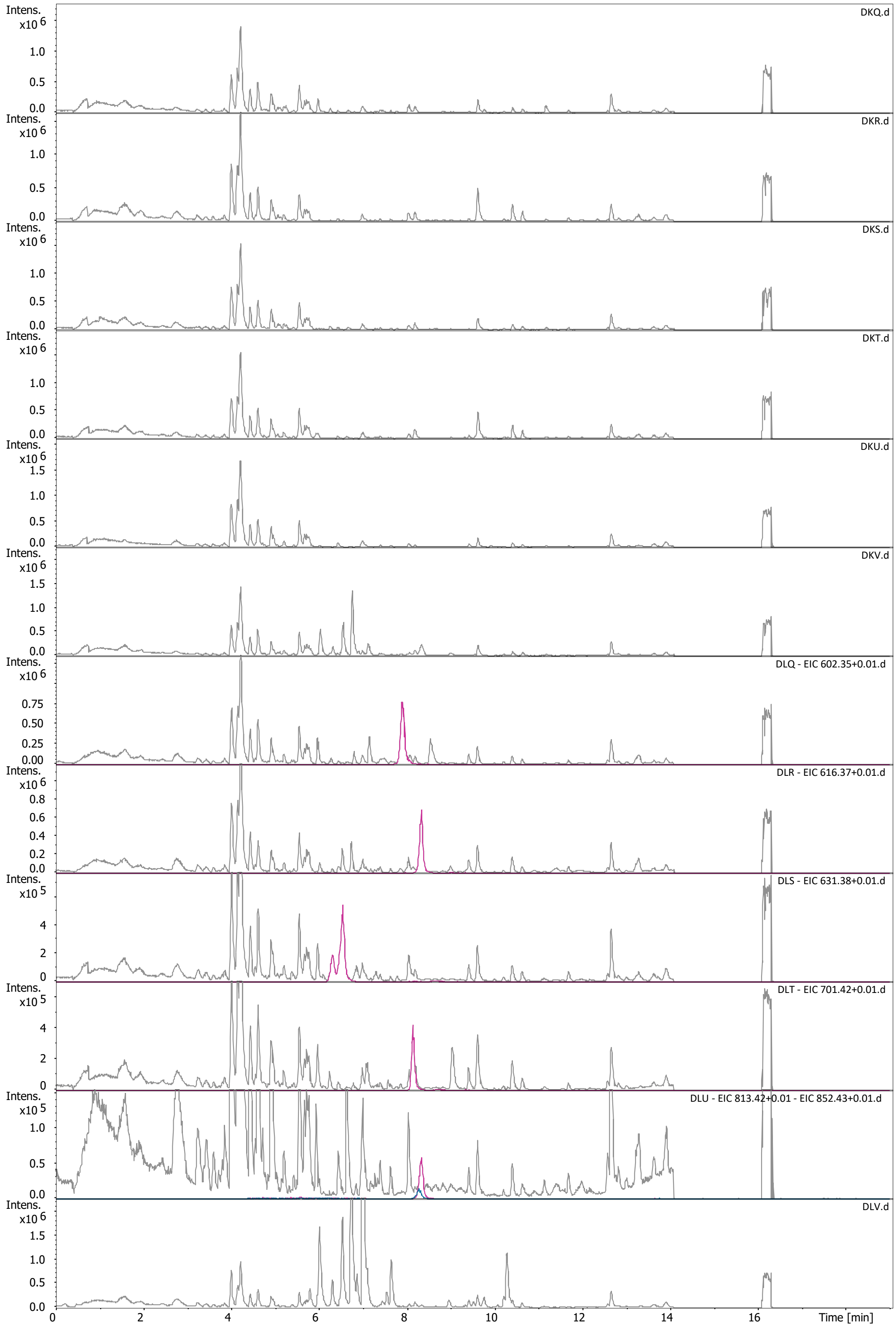
Intens.
x10 6
DKQ.d
DKR.d
DKS.d
DKT.d
DKU.d
DKV.d
DLQ - EIC 602.35+0.01.d
DLR - EIC 616.37+0.01.d
x10 5
DLS - EIC 631.38+0.01.d
DLT - EIC 701.42+0.01.d
DLU - EIC 813.42+0.01 - EIC 852.43+0.01.d
DLV.d
Time [min]

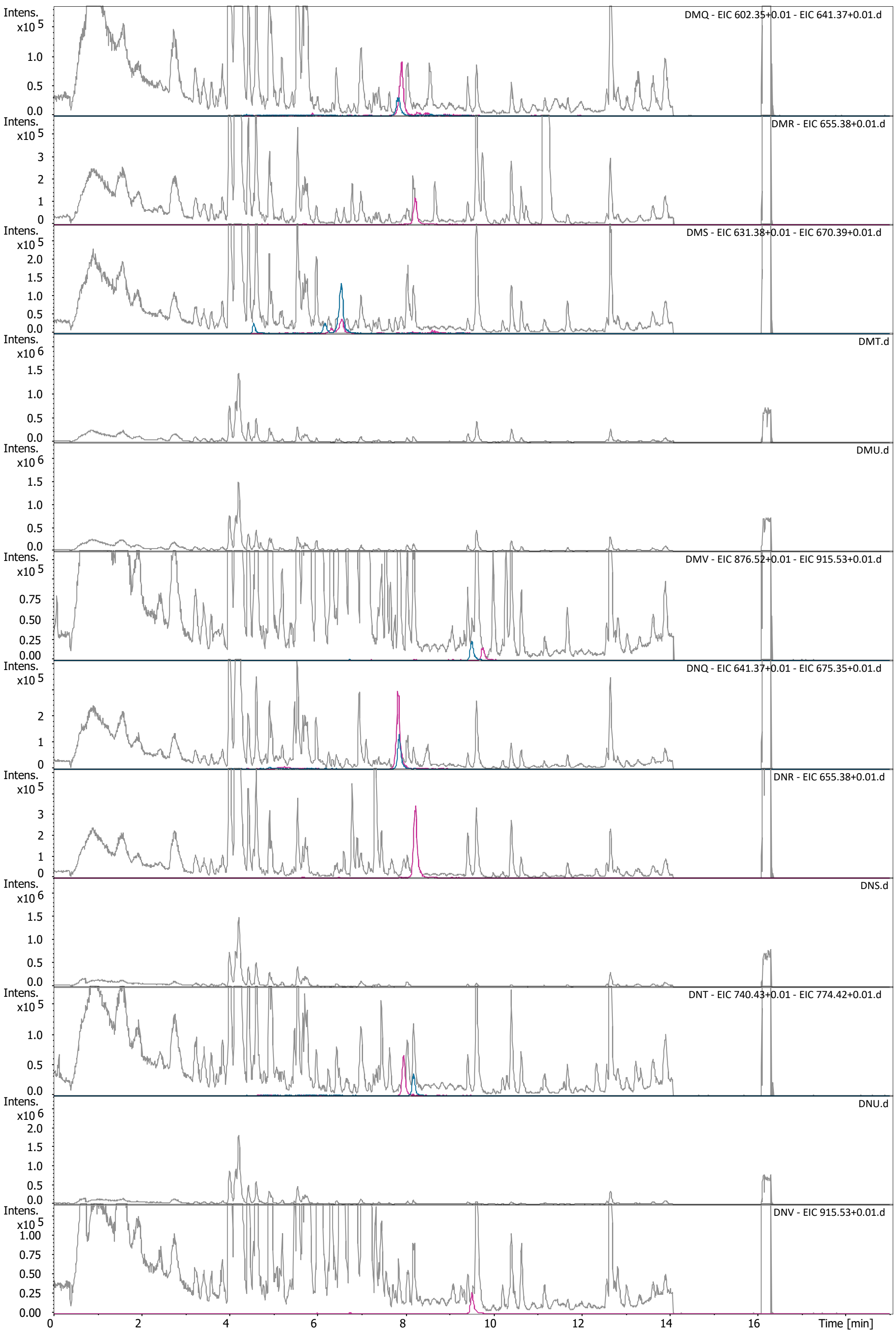

DMQ - EIC 602.35+0.01 - EIC 641.37+0.01.d
DMR - EIC 655.38+0.01.d
DMS - EIC 631.38+0.01 - EIC 670.39+0.01.d
DMT.d
DMU.d
DMV - EIC 876.52+0.01 - EIC 915.53+0.01.d
DNQ - EIC 641.37+0.01 - EIC 675.35+0.01.d
DNR - EIC 655.38+0.01.d
DNS.d
DNT - EIC 740.43+0.01 - EIC 774.42+0.01.d
DNU.d
DNV - EIC 915.53+0.01.d
Intens.
Time [min]

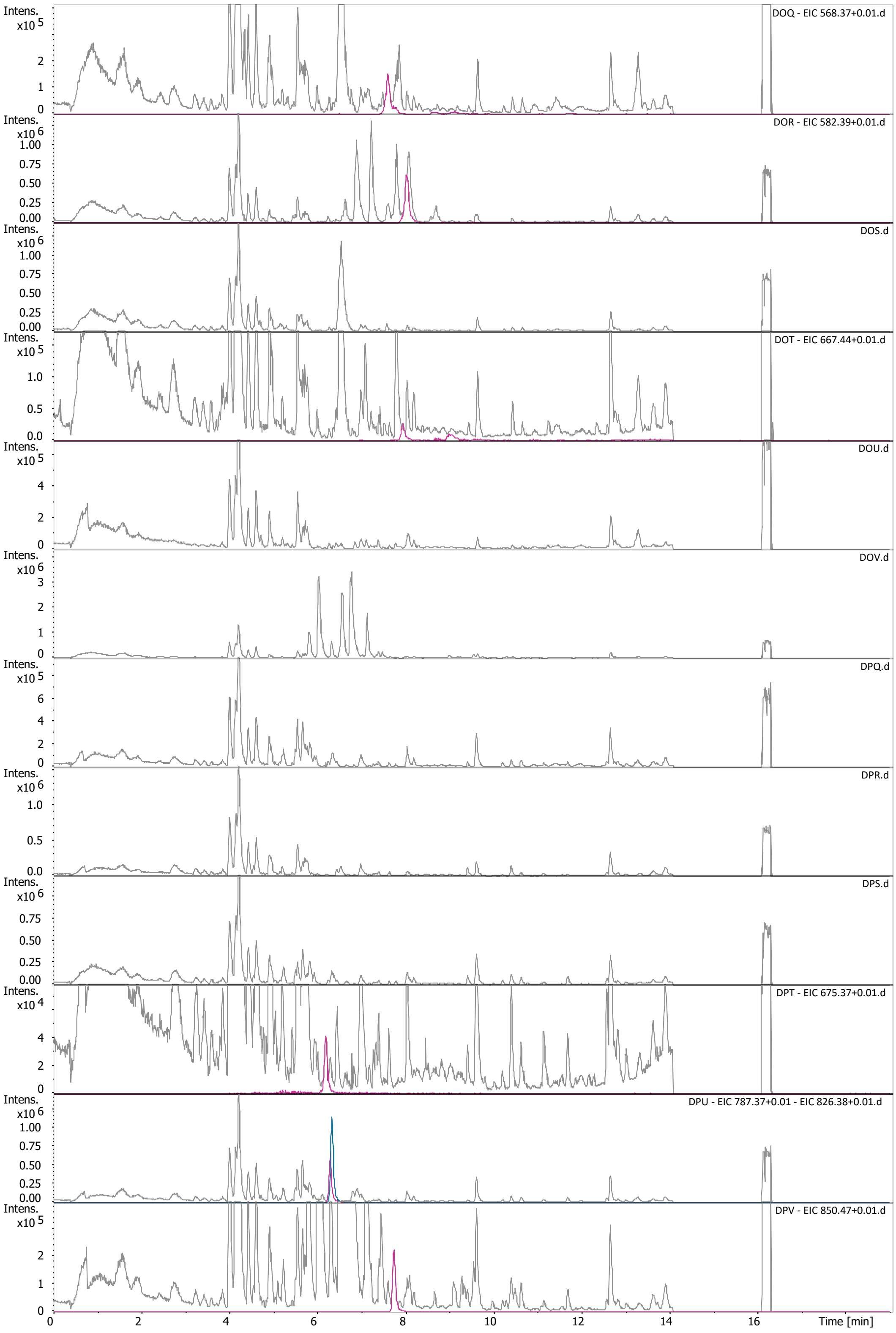
DOQ - EIC 568.37+0.01.d
DOR - EIC 582.39+0.01.d
DOS.d
DOT - EIC 667.44+0.01.d
DOU.d
DOV.d
DPQ.d
DPR.d
DPS.d
DPT - EIC 675.37+0.01.d
DPU - EIC 787.37+0.01 - EIC 826.38+0.01.d
DPV - EIC 850.47+0.01.d
Intens.
Time [min]

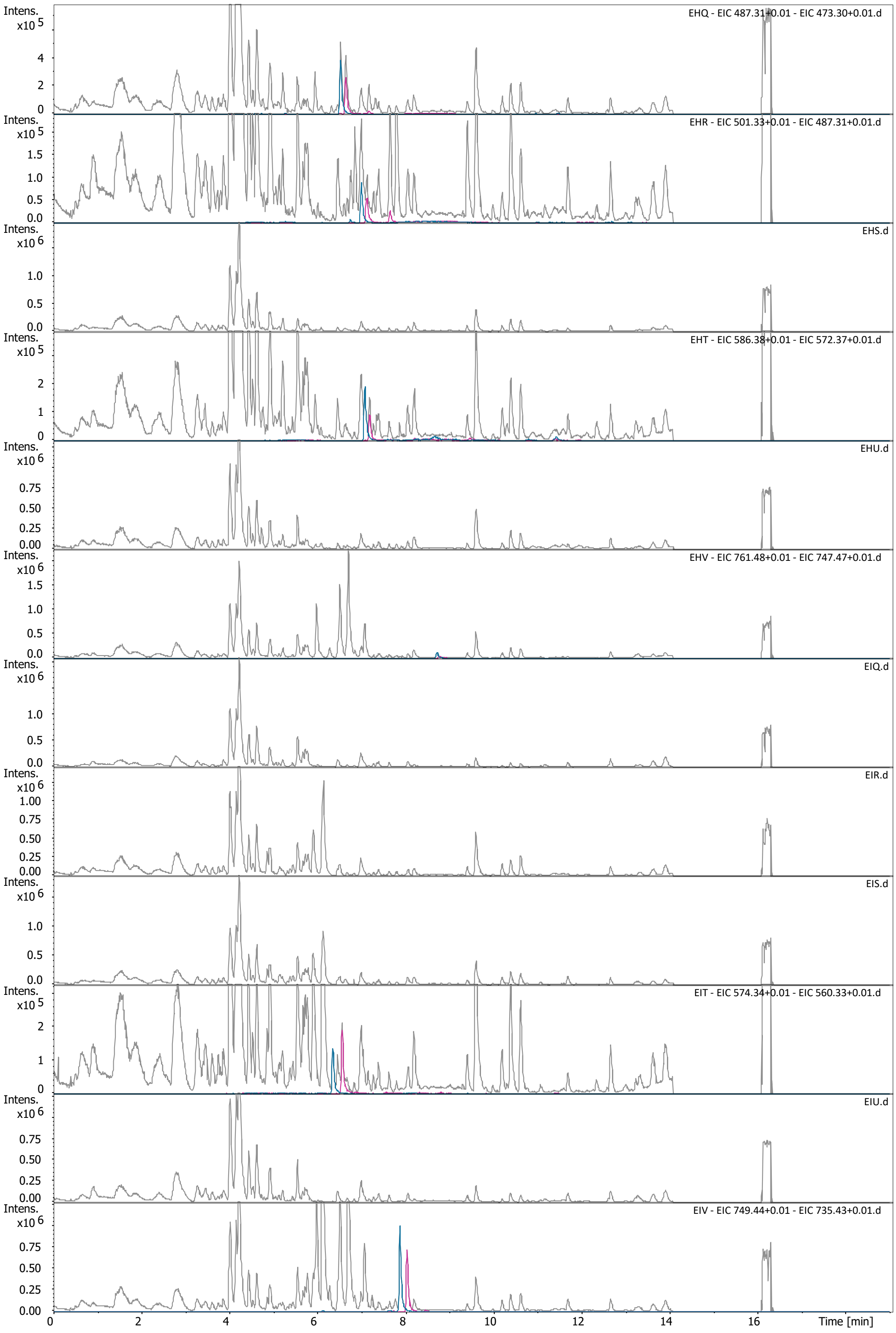
EHQ - EIC 487.31±0.01 - EIC 473.30±0.01.d
EHR - EIC 501.33±0.01 - EIC 487.31±0.01.d
EHS.d
EHT - EIC 586.38±0.01 - EIC 572.37±0.01.d
EHU.d
EHV - EIC 761.48±0.01 - EIC 747.47±0.01.d
EIQ.d
EIR.d
EIS.d
EIT - EIC 574.34±0.01 - EIC 560.33±0.01.d
EIU.d
EIV - EIC 749.44±0.01 - EIC 735.43±0.01.d
Intens.
Time [min]

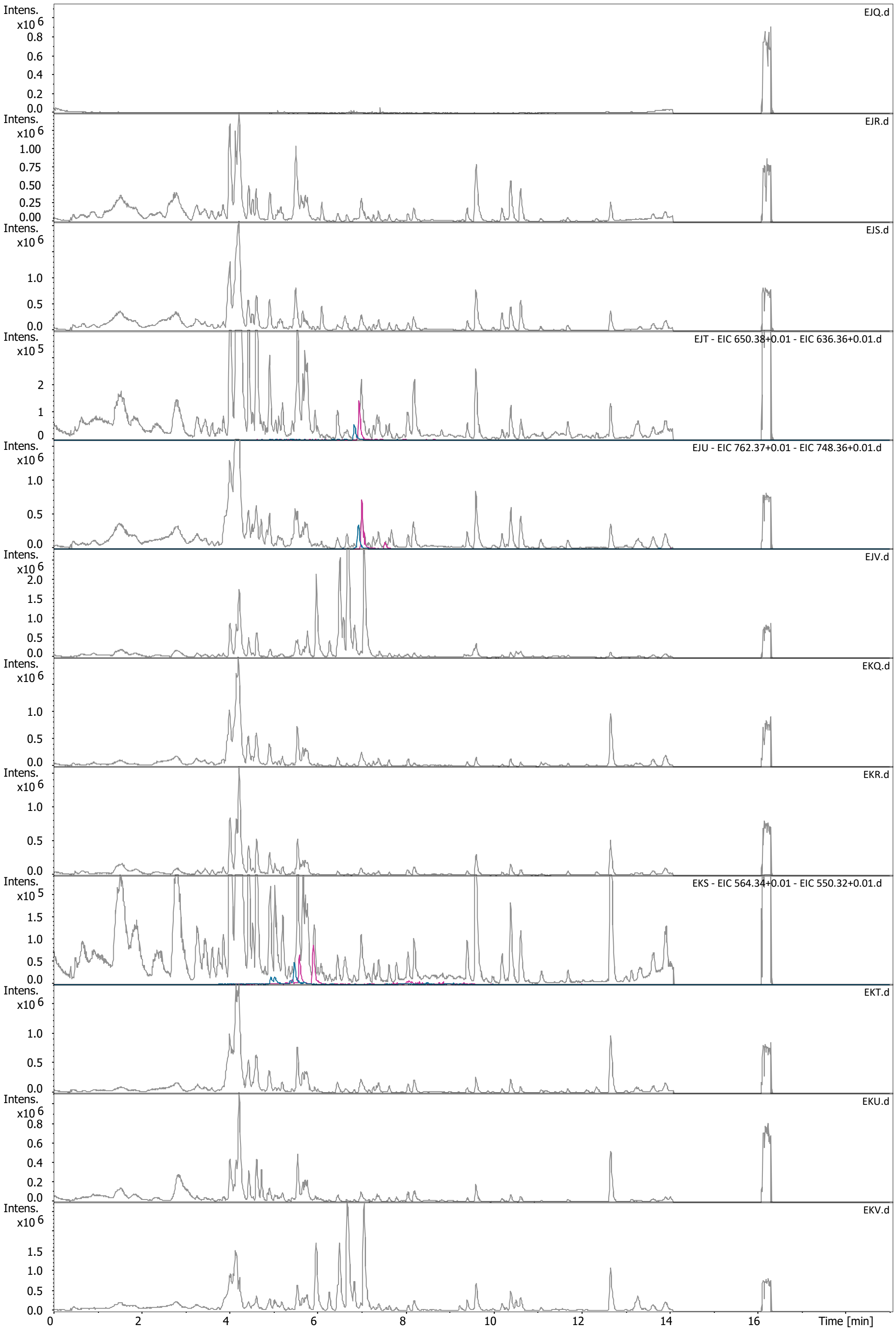

EJQ.d
EJR.d
EJS.d
EJT - EIC 650.38+0.01 - EIC 636.36+0.01.d
EJU - EIC 762.37+0.01 - EIC 748.36+0.01.d
EJV.d
EKQ.d
EKR.d
EKS - EIC 564.34+0.01 - EIC 550.32+0.01.d
EKT.d
EKU.d
EKV.d
Intens.
Time [min]

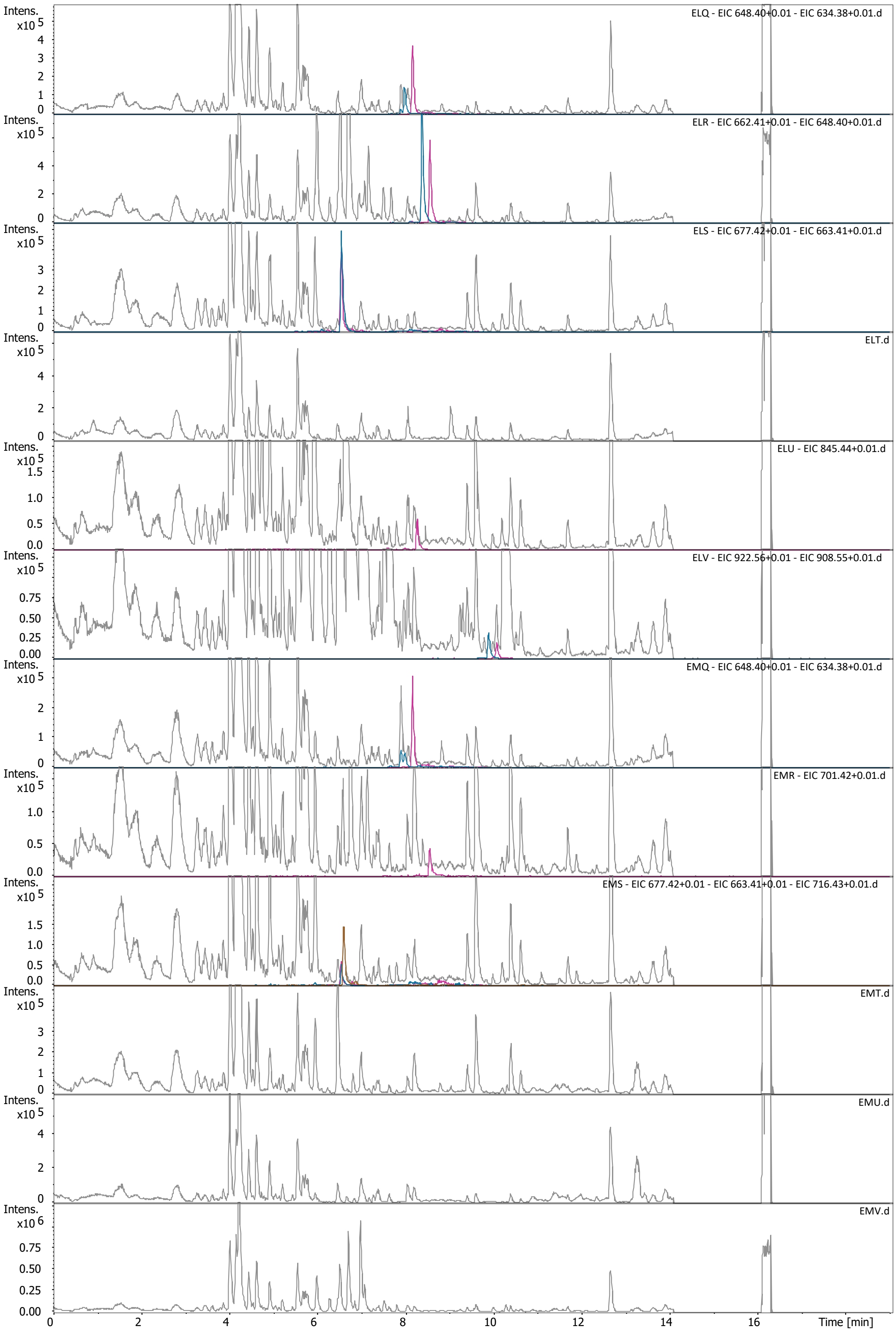

ELQ - EIC 648.40+0.01 - EIC 634.38+0.01.d
ELR - EIC 662.41+0.01 - EIC 648.40+0.01.d
ELS - EIC 677.42+0.01 - EIC 663.41+0.01.d
ELT.d
ELU - EIC 845.44+0.01.d
ELV - EIC 922.56+0.01 - EIC 908.55+0.01.d
EMQ - EIC 648.40+0.01 - EIC 634.38+0.01.d
EMR - EIC 701.42+0.01.d
EMS - EIC 677.42+0.01 - EIC 663.41+0.01 - EIC 716.43+0.01.d
EMT.d
EMU.d
EMV.d
Intens.
Time [min]

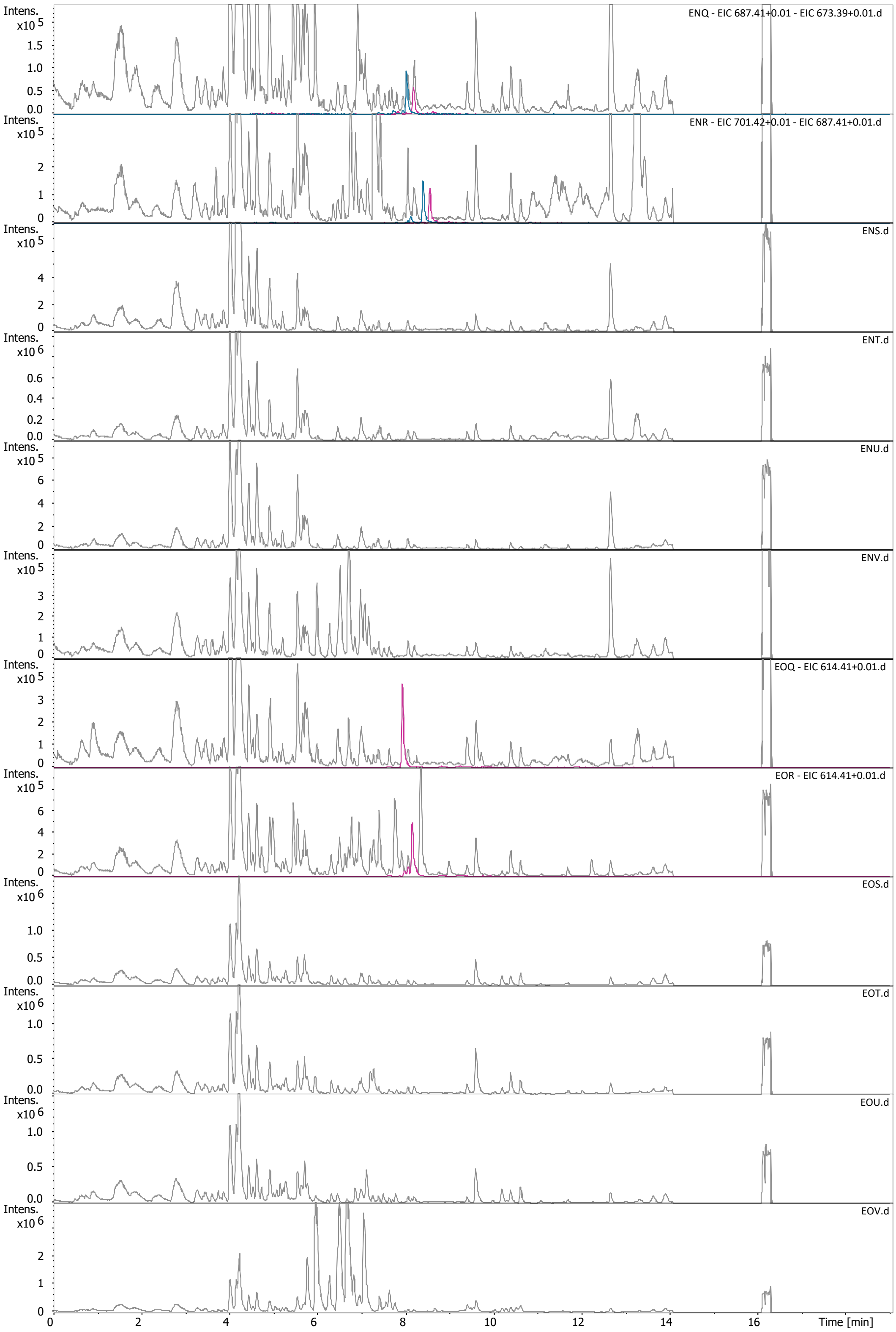

ENQ - EIC 687.41+0.01 - EIC 673.39+0.01.d
ENR - EIC 701.42+0.01 - EIC 687.41+0.01.d
ENS.d
ENT.d
ENU.d
ENV.d
EOQ - EIC 614.41+0.01.d
EOR - EIC 614.41+0.01.d
EOS.d
EOT.d
EOU.d
EOV.d
Intens.
Time [min]

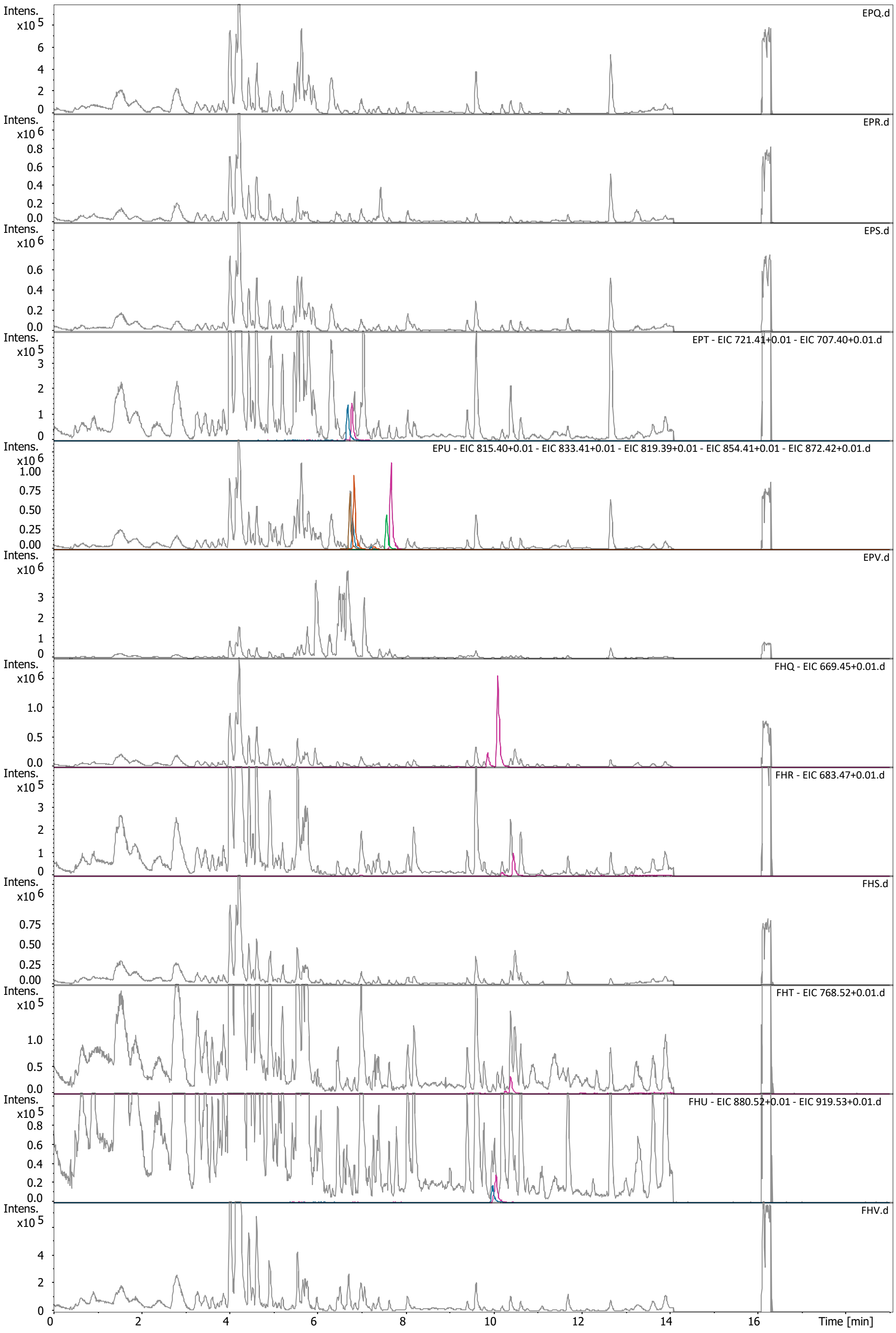
EPQ.d
EPR.d
EPS.d
EPT - EIC 721.41+0.01 - EIC 707.40+0.01.d
EPU - EIC 815.40+0.01 - EIC 833.41+0.01 - EIC 819.39+0.01 - EIC 854.41+0.01 - EIC 872.42+0.01.d
EPV.d
FHQ - EIC 669.45+0.01.d
FHR - EIC 683.47+0.01.d
FHS.d
FHT - EIC 768.52+0.01.d
FHU - EIC 880.52+0.01 - EIC 919.53+0.01.d
FHV.d
Intens.
Time [min]

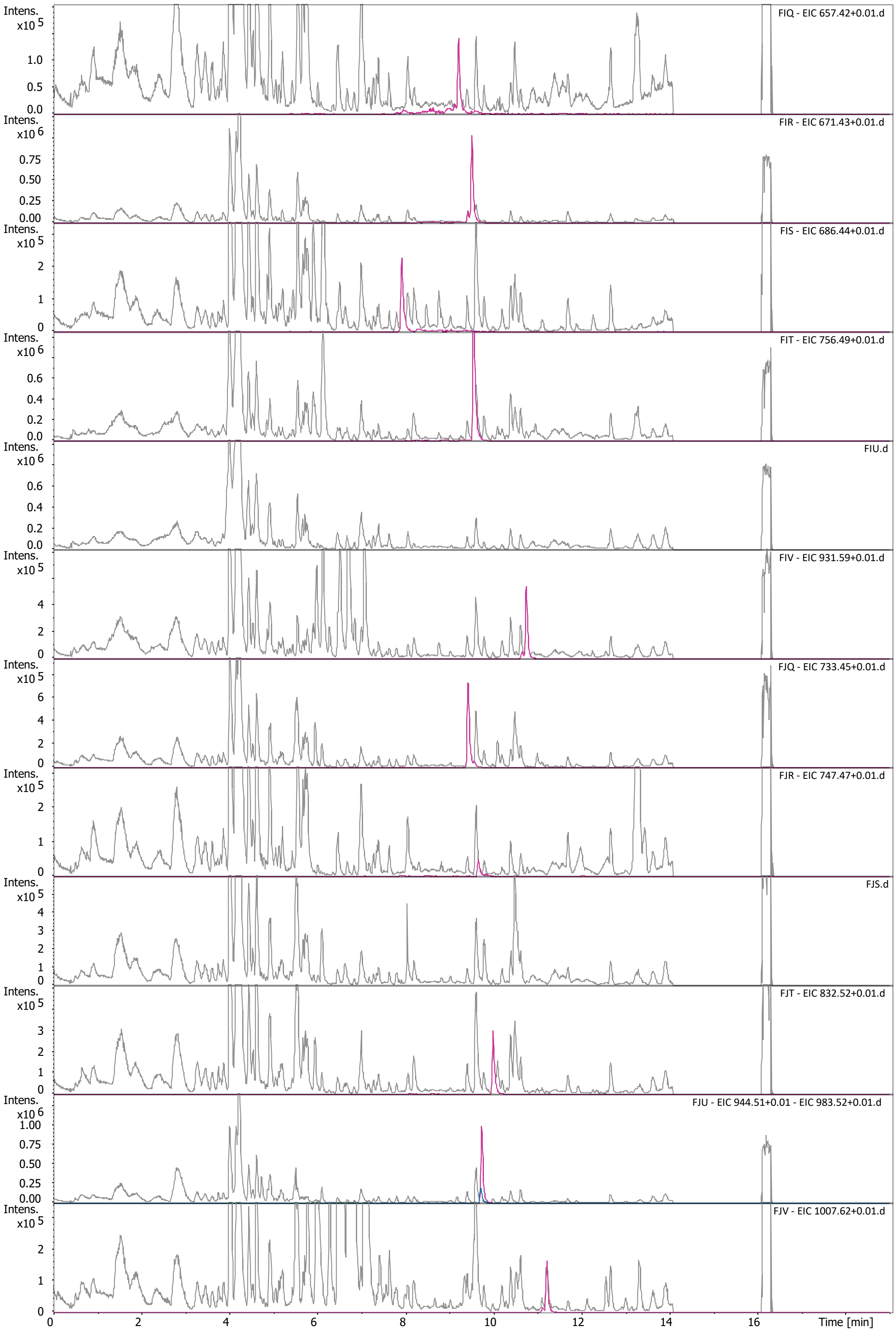

Intens. x10 5
FIQ - EIC 657.42+0.01.d
Intens. x10 6
FIR - EIC 671.43+0.01.d
Intens. x10 5
FIS - EIC 686.44+0.01.d
Intens. x10 6
FIT - EIC 756.49+0.01.d
Intens. x10 6
FIU.d
Intens. x10 5
FIV - EIC 931.59+0.01.d
Intens. x10 5
FJQ - EIC 733.45+0.01.d
Intens. x10 5
FJR - EIC 747.47+0.01.d
Intens. x10 5
FJS.d
Intens. x10 5
FJT - EIC 832.52+0.01.d
Intens. x10 6
FJU - EIC 944.51+0.01 - EIC 983.52+0.01.d
Intens. x10 5
FJV - EIC 1007.62+0.01.d
Time [min]

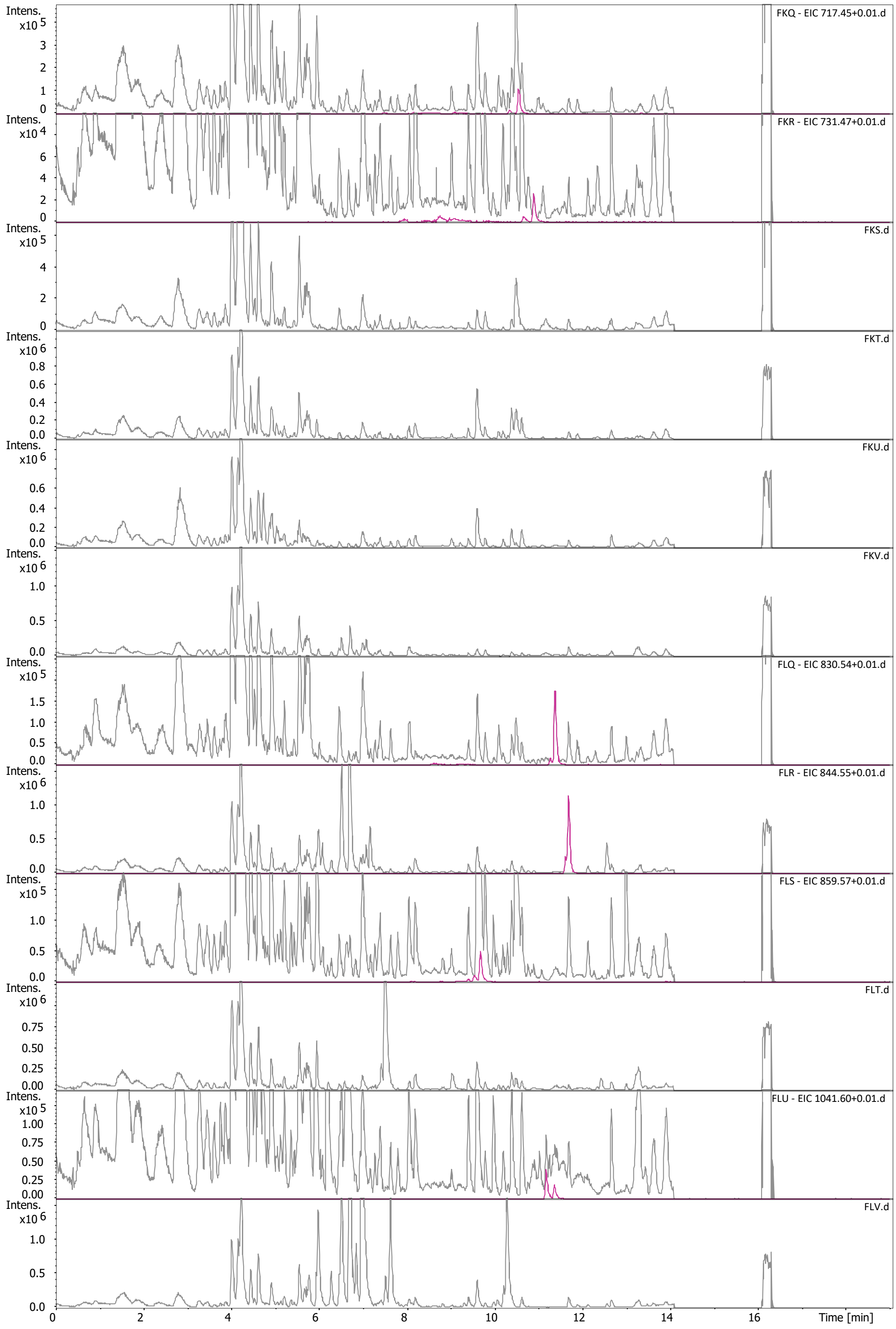

FKQ - EIC 717.45+0.01.d
FKR - EIC 731.47+0.01.d
FKS.d
FKT.d
FKU.d
FKV.d
FLQ - EIC 830.54+0.01.d
FLR - EIC 844.55+0.01.d
FLS - EIC 859.57+0.01.d
FLT.d
FLU - EIC 1041.60+0.01.d
FLV.d
Intens.
Time [min]

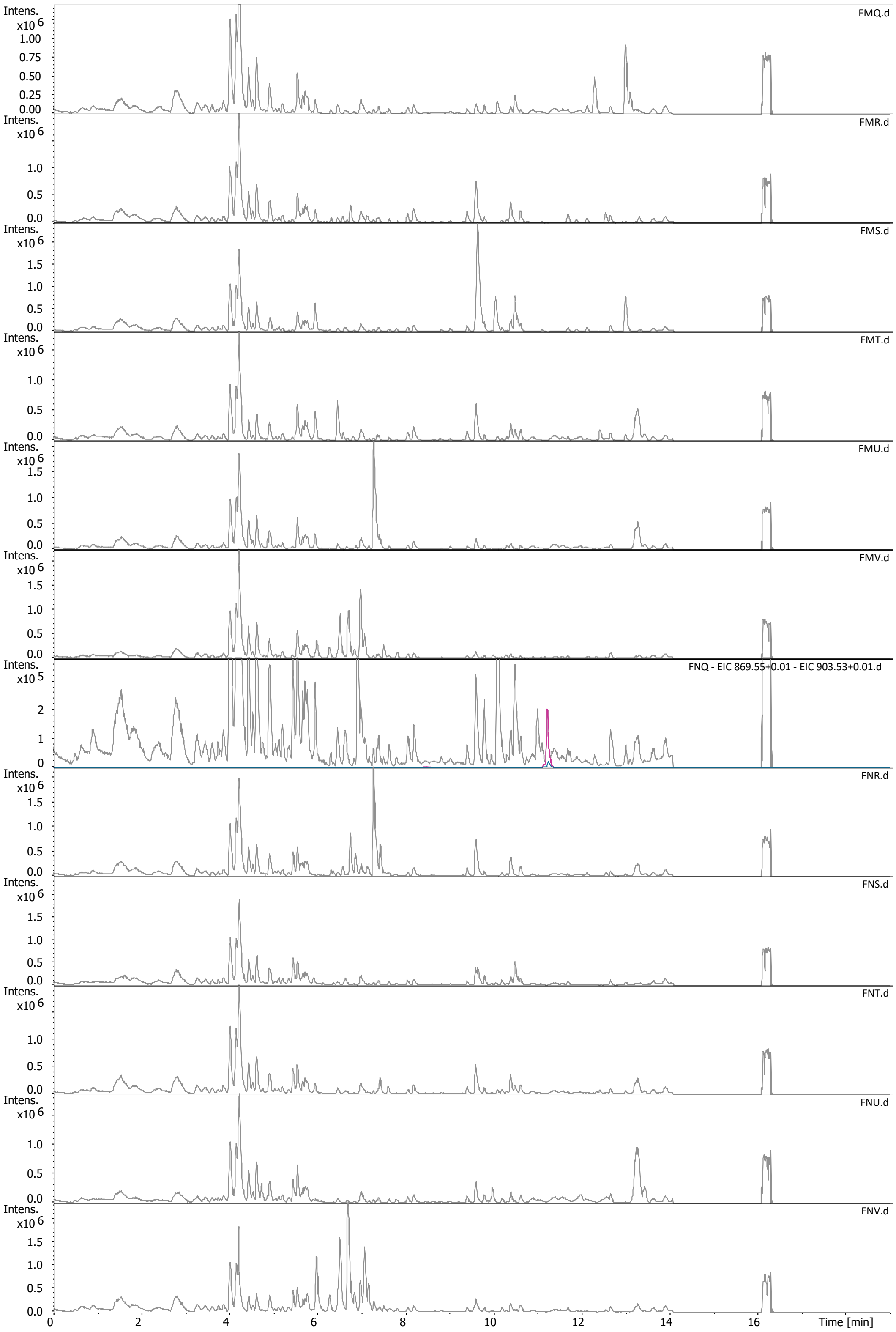
FMQ.d
FMR.d
FMS.d
FMT.d
FMU.d
FMV.d
FNQ - EIC 869.55+0.01 - EIC 903.53+0.01.d
FNR.d
FNS.d
FNT.d
FNU.d
FNV.d
Intens.
x10 6
x10 5
Time [min]

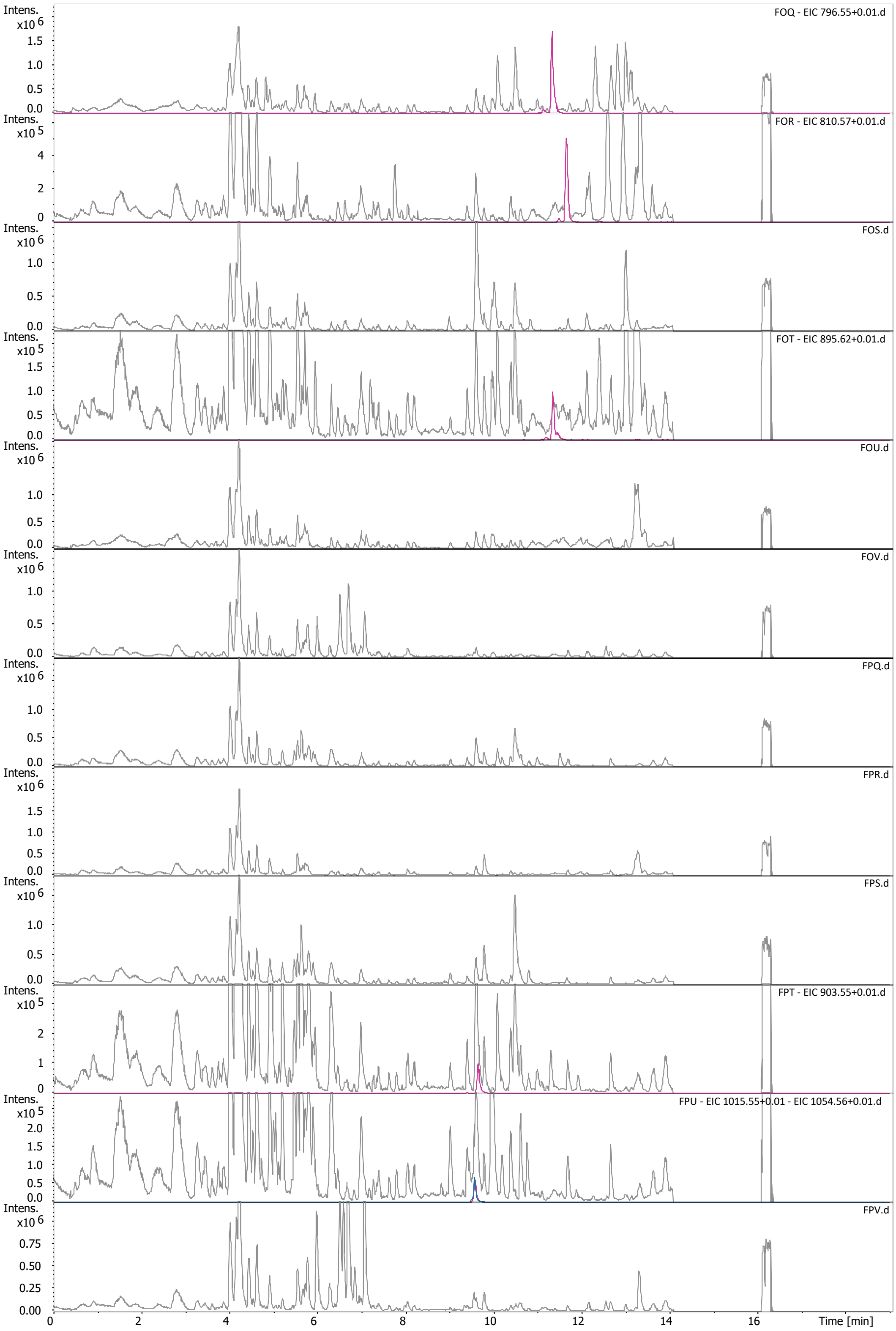
FOQ - EIC 796.55+0.01.d
FOR - EIC 810.57+0.01.d
FOS.d
FOT - EIC 895.62+0.01.d
FOU.d
FOV.d
FPQ.d
FPR.d
FPS.d
FPT - EIC 903.55+0.01.d
FPU - EIC 1015.55+0.01 - EIC 1054.56+0.01.d
FPV.d
Intens.
Time [min]

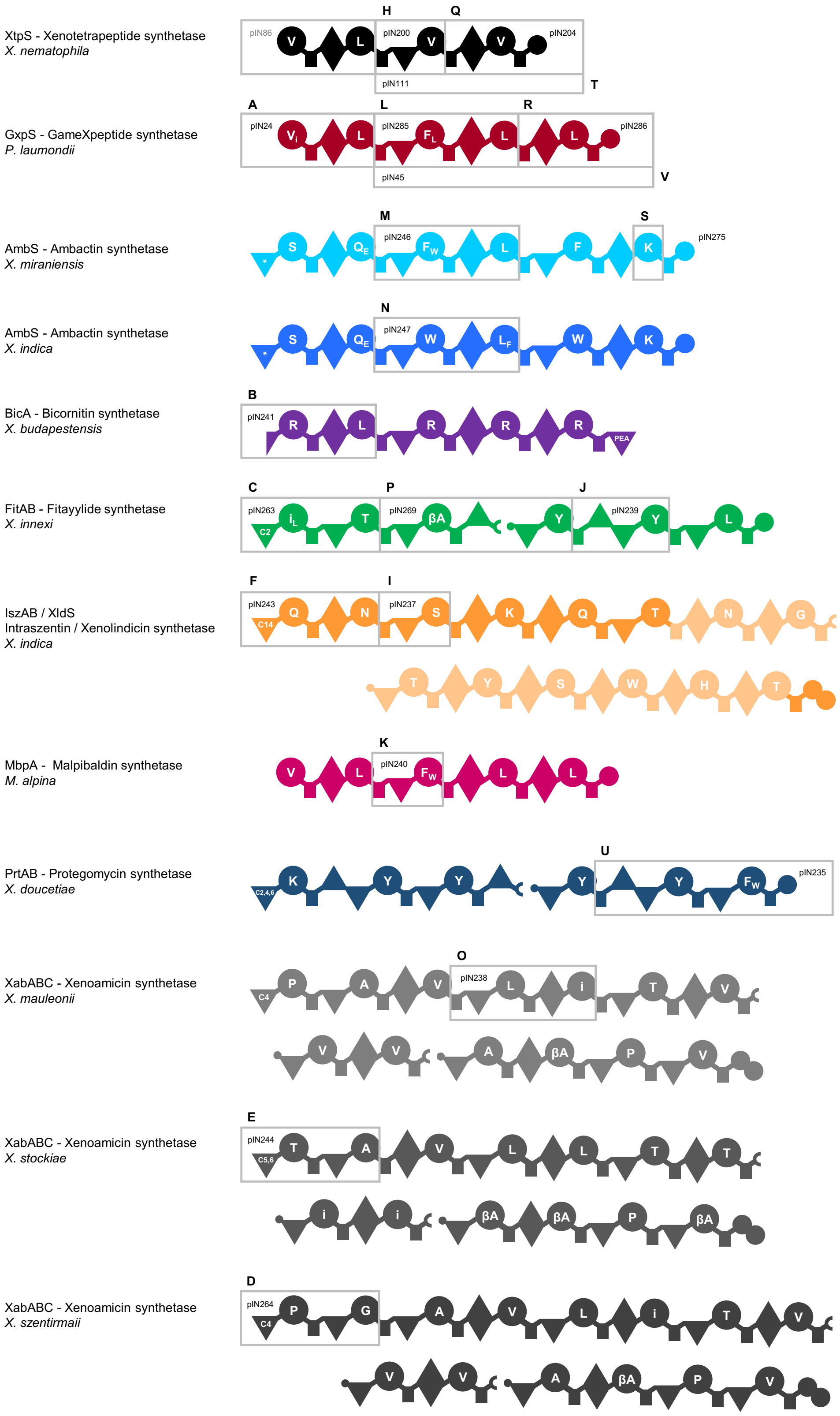

**Figure S3. | Overview of all native NRPS clusters used in this work.** Squares show XUT[I] units generated from these clusters with respective letters. Chemical structurs of corresponding products are shown in Figure S4.

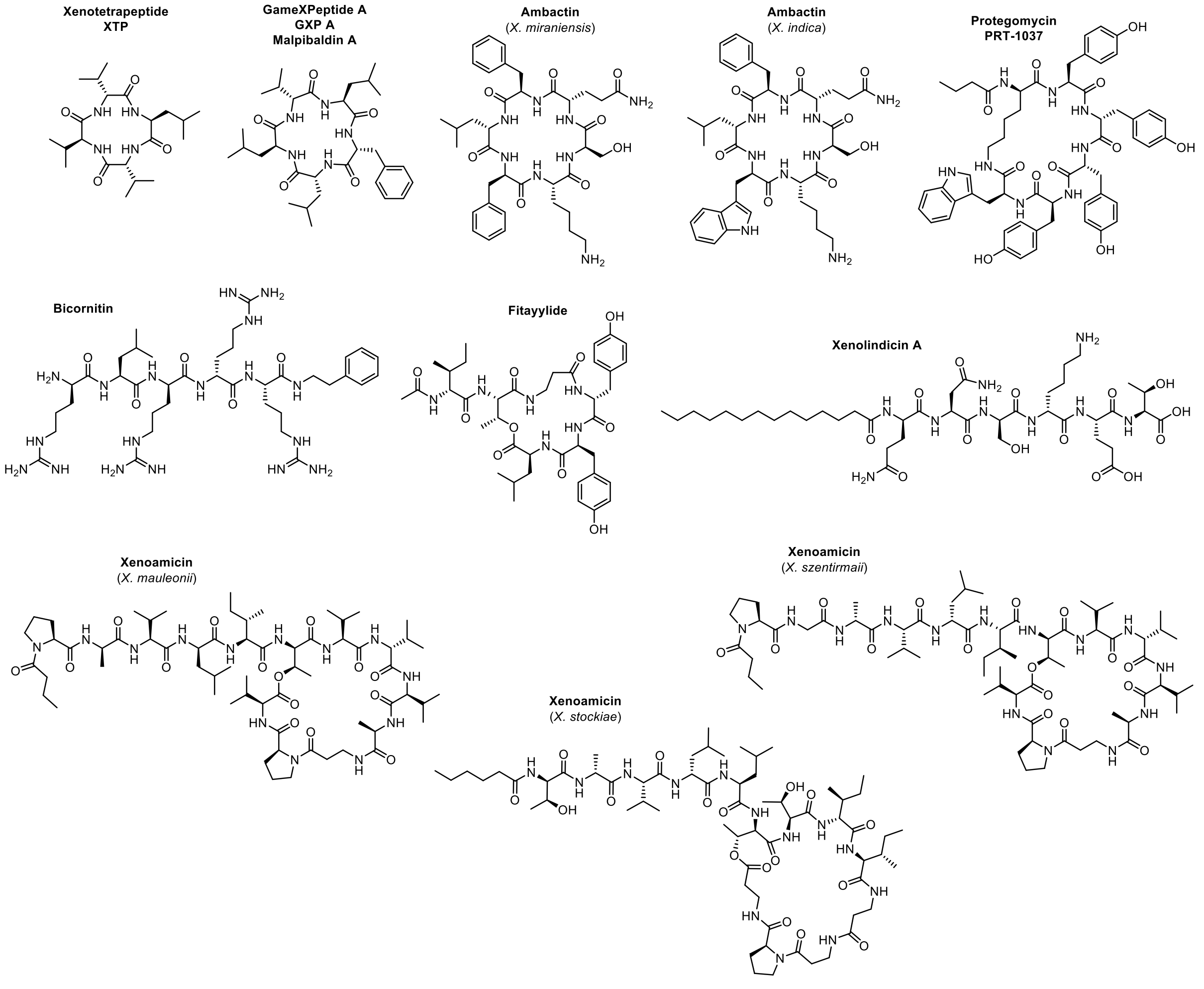

**Figure S4. | Chemical structures of natural products used in this work.**

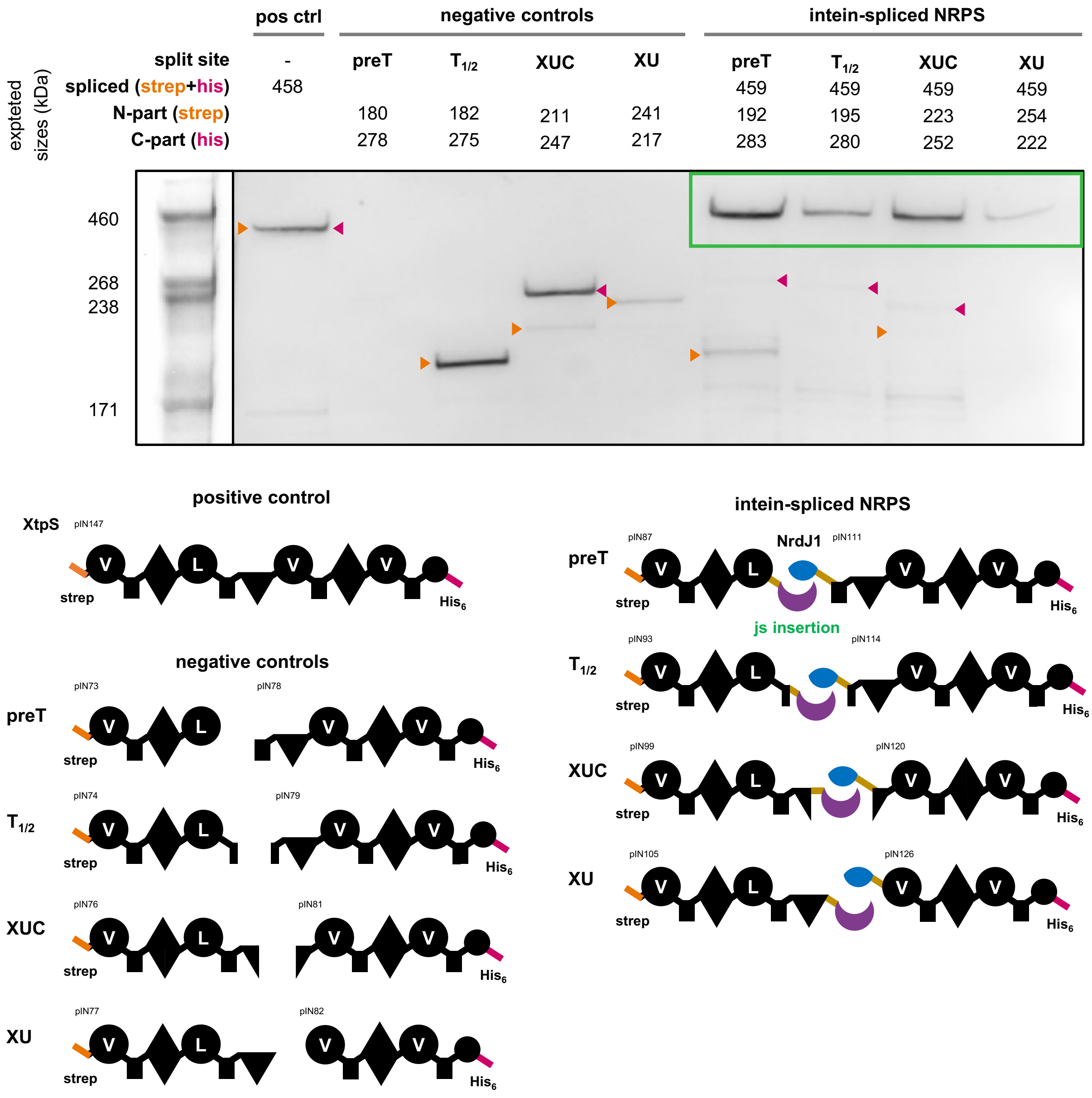

**Figure S5. | Western blot analysis.** XtpS was expressed with an N-terminal Strep-tag and a C-terminal His6-tag, either from a single plasmid (positive control) or split at one of four tested engineering sites. In the split constructs, expression occurred from two separate plasmids, either without inteins (negative controls) or with the corresponding halves of the split intein NrdsJ-1 fused to the protein fragments. Proteins were detected simultaneously with anti-poly-Histidine HRP conjugates and StrepTag II antibody HRP conjugates, allowing detection of both the N- and C-terminal half of the NRPS. N-terminal fragments of the expected sizes are marked with orange arrows, C-terminal fragments with red arrows, and bands corresponding to intein-spliced products are highlighted with a green box.

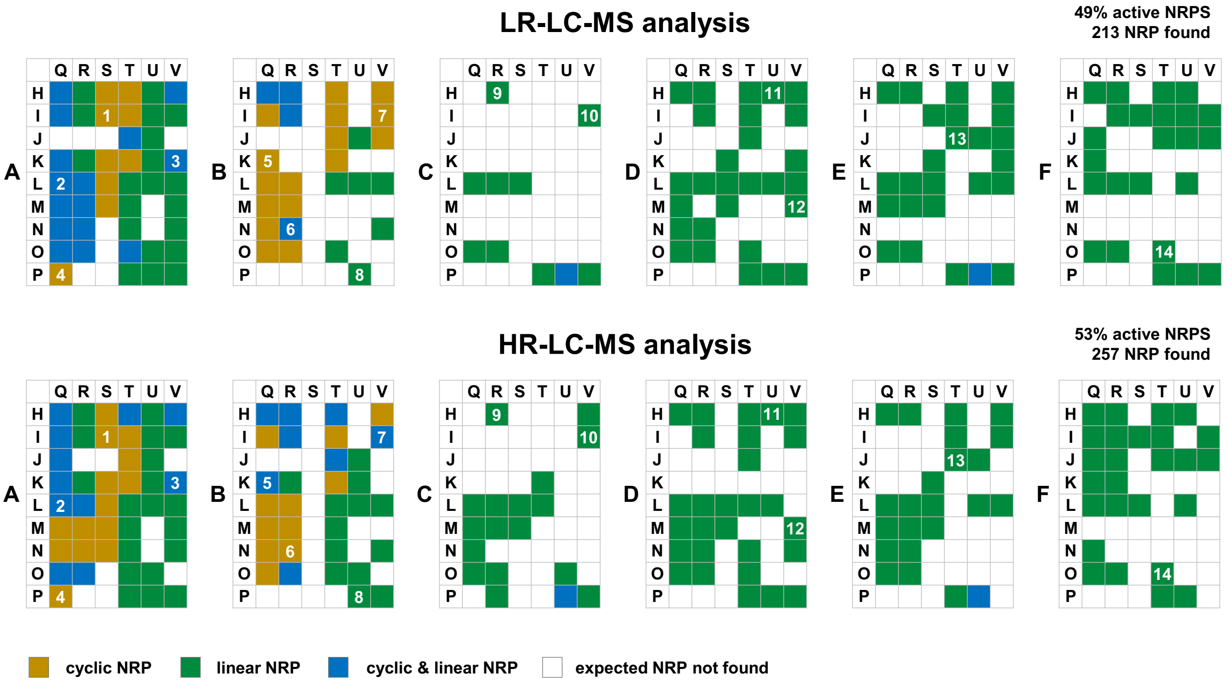

**Figure S6. | Results of LR-LC-MS analysis and comparison to HR results.**
